# iFLinkC-EZ: A scalable and automatable method for the assembly of complex fusion proteins and multi-gene expression constructs based on the iFLinkC framework

**DOI:** 10.1101/2025.10.08.680606

**Authors:** Philipp Kemp, Melinda Blat Belmonte, Max Schmidt, Klara Eisenhauer, Carolin Gebhardt, Kai Kabuth, Heinz Koeppl, Viktor Stein

## Abstract

Standardized methods for the assembly of DNA constructs have become indispensable in synthetic biology, biotechnology and basic research. The majority of frameworks – most prominently based on Golden Gate – have been developed and optimized for the assembly of transcriptional units (TUs) and thereof composed multi-gene expression constructs. This is significantly enabled by the modular organization of functional elements underlying any given TU. In contrast, the assembly of protein coding sequences typically necessitates tailored approaches. Addressing this technological gap, iterative functional linker cloning (iFLinkC) was recently developed to provide a standardized framework for assembling protein coding sequences of arbitrary size and complexity from the ground up based on pairwise ligations of any two DNA fragments *via* a two-base overlap. Yet, a need for physically purifying DNA fragments imposes significant procedural complexity and operator skill. Overcoming these limitations, a new assembly algorithm, termed iFLinkC-EZ, is presented. Crucially, the DNA assembly products are now purified by genetic means which greatly simplifies the underlying assembly process and also renders it compatible with robotic automation. Further, the potency of iFLinkC-EZ is demonstrated in the assembly of mono- and poly-chromatic poly-fluorescent fusion proteins as well as poly-cistronic expression constructs while examining the effect of poly-PT and poly-GGS linkers as well as ribosome binding sites on the functional expression of poly-fluorescent fusion protein constructs. Further, the utility of poly-fluorescent proteins is demonstrated for the enhanced labeling of surface displayed recombinant binders in yeast display. Given its ease and efficiency, iFLinkC-EZ is anticipated to be applicable to the assembly of many different types of fusion proteins and thereof based multi-gene expression constructs including but not limited to synthetic protein switches, sensors and multi-enzyme complexes as well as thereof composed genetic circuits and metabolic pathways.

## Introduction

Standardized methods for assembling DNA constructs from smaller DNA fragments have become indispensable in basic research, biotechnology and synthetic biology (Casini et al. 2015). Ideally, DNA assembly methods feature low cost, low skill, low effort, and high fidelity. Further, they are readily scalable and amenable to robotic automation. Mechanistically, DNA assembly can be broadly grouped into methods based on homologous (Bomfiglio et al. 2025) and non-homologous DNA recombination (Laborda-Mansilla and García-Ruiz 2025) depending whether they direct the ligation of >2 DNA fragments either based on 25-50 bp dsDNA or short ssDNA overhangs. From a functional perspective, homology-dependent DNA assembly methods typically afford the greatest flexibility while non-homology DNA assembly methods are most scalable.

In terms of non-homology-dependent DNA assembly methods, Golden Gate has proven the most versatile and standardizable approach (Engler et al. 2008) giving rise to numerous modular cloning (MoClo) frameworks (Bird et al. 2022). These typically feature toolboxes of well-defined parts that are stored in entry plasmids (Level 0) that are first assembled into a transcriptional unit (TUs) (Level 1) and then into a large multi gene expression constructs (Level 2). Crucially, the positioning of **(i)** individual parts (e.g. promoters, ribosome binding sites, open reading frames and terminators) within a TU, and the positioning of **(ii)** TUs within a multi gene expression construct is pre-defined *via* distinct 4 bp sequence tags. While affording scalability, a pre-defined positioning of individual parts renders conventional MoClo frameworks unsuitable for assembling modular fusion proteins which calls for flexibility and reusability in terms of DNA fragment size and the relative orientation of protein domains and connecting linkers.

Addressing these limitations, iterative functional linker cloning (iFLinkC) was recently developed for assembling fusion proteins (Gräwe et al. 2020, 2021, 2022). Unlike conventional Golden Gate and MoClo frameworks, iFLinkC restriction digests and ligates DNA fragments in a pairwise and sequential rather than parallel fashion. Further, recombination is directed *via* constant 2-base as opposed to variable 4-base overhangs (**Fig. 1A**). Crucially, this affords flexibility as DNA fragments of arbitrary size can be appended at either end of any given DNA construct. Crucially, the product of any given pairwise DNA assembly reaction readily serves as the input for the subsequent assembly step. In this way, DNA constructs of arbitrary size and complexity can be assembled in an iterative fashion from the ground up. iFLinkC has proven especially powerful in the assembly of complex fusion proteins – notably, protein switches and sensors – where the ligation of individual domains *via* defined linkers calls for flexibility in terms of **(i)** N- and C-terminal positioning, **(ii)** fragment size which can be as small as one amino acid and **(iii)** a capacity for combinatorial linker engineering (Gräwe et al. 2020, 2022; Weber et al. 2022; Kemp et al. 2023). Yet, the need for purifying DNA fragments by agarose gel electrophoresis currently imposes substantial procedural complexity and limits the scope for robotic automation.

**Figure 1:**
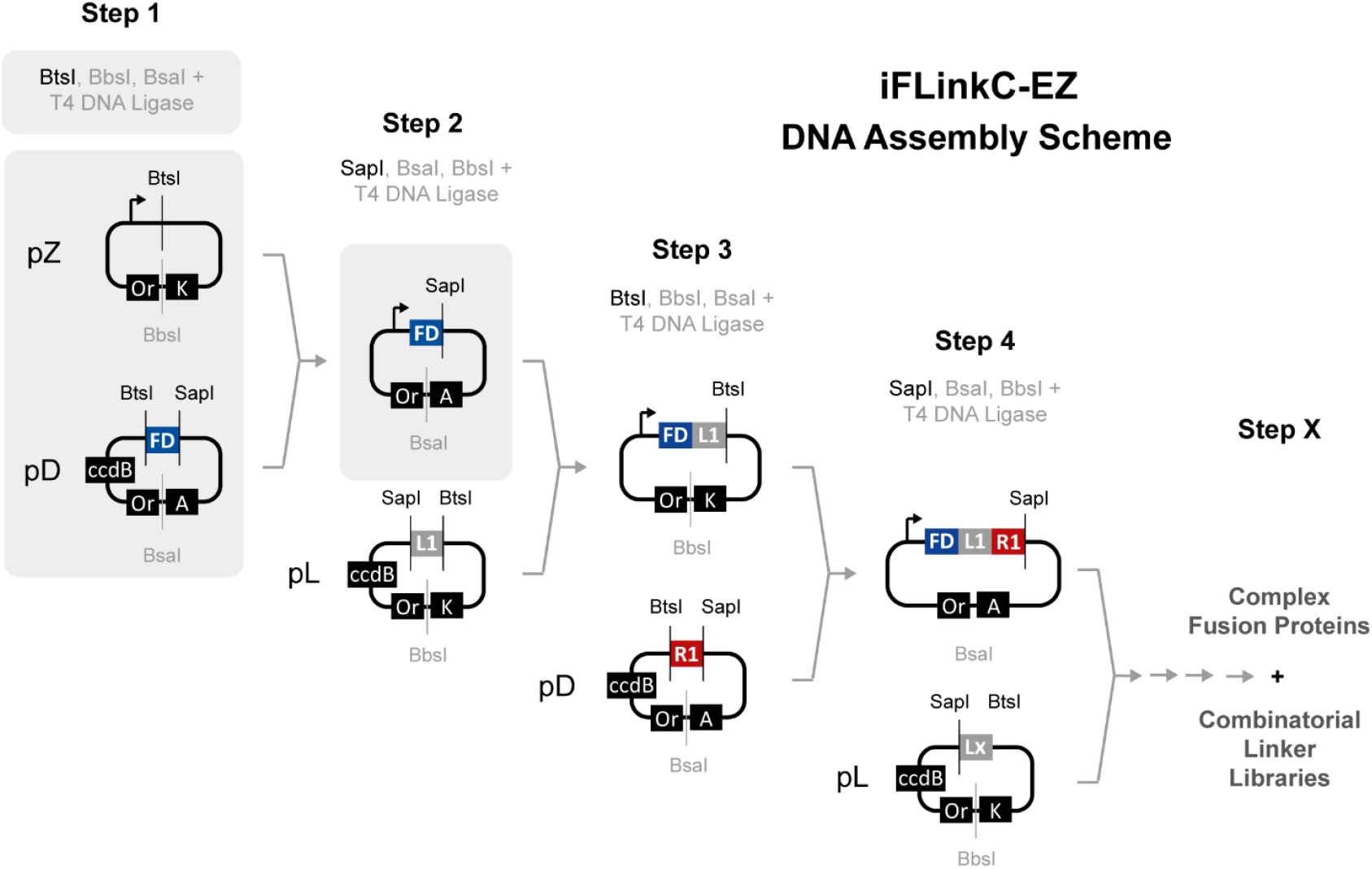

Schematic overview of iFLinkC-EZ mediated DNA assembly with an elementary assembly cycle highlighted in grey. DNA constructs are directly assembled in a destination plasmid pZ while pD and pL coding for either a functional domain or a linker element are iteratively recombined with pZ. To this end, pZ along with pD and pL are restriction digested with combinations of either BtsI or SapI and BbsI or BsaI. The resultant 2- or 3-base overhangs then direct recombination of either pD or pL with the destination plasmid pZ harboring the growing fusion protein. Adding T5 exonuclease removes linear DNA contaminants while the resultant output plasmid pZ is genetically purified upon transformation through a change in antibiotic resistance in pZ and ccdB counterselection against either pD or pL.

Overcoming these shortcomings, an improved assembly algorithm now renders iFLinkC very *easy* (EZ) to conduct. Crucially, any input DNA is removed genetically by means of a double negative selection step which in turn obviates physical purification by agarose gel electrophoresis. This reduces experimental complexity and shortens an elementary assembly cycle to one day. Further, it renders iFLinkC-EZ compatible with robotic automation. Proof-of-concept is demonstrated in the assembly of poly-fluorescent proteins (FP) *via* poly-PT and poly-GGS linkers while assessing their functional expression and functionality in yeast display. Further, dedicated adaptor modules can be used to construct large poly-cistronic expression constructs effectively interfacing iFLinkC-EZ with alternative DNA assembly strategies such as Golden Gate. Overall, we anticipate that iFLinkC-EZ will be applicable and greatly facilitate the DNA assembly of many different types of fusion protein constructs. In particular, this concerns synthetic protein switches and sensors (Gräwe and Stein 2021) but is also readily conceivable to multi-enzyme complexes as well as thereof based genetic circuits and metabolic pathways.

## Results and Discussion

### Conception of iFLinkC-EZ

iFLinkC comprises a scalable DNA assembly framework which recombines two dedicated input plasmids into one output plasmid based on the sequential action of two separate type IIS restriction enzymes, BtsI and BsrDI, and T4 DNA ligase (Gräwe et al. 2020, 2021, 2022). Crucially, the resultant output plasmid is regenerated upon completing an elementary iFLinkC DNA assembly cycle enabling individual steps to be performed iteratively and in parallel in order to build DNA constructs of arbitrary complexity from the ground up. iFLinkC has proven particular amenable in the construction of complex fusion proteins as any two (protein coding) DNA sequences are fused independent of length and sequence composition *via* a minimal two base overhang that, in the context of a protein, results in a minimal single amino acid scar (Gräwe et al. 2020). iFLinkC has proven particularly amenable in the assembly of protein switches, protein sensors and recombinant nanopores and thereof based combinatorial linker libraries (Gräwe et al. 2020, 2022; Weber et al. 2022; Kemp et al. 2023). Yet, the need for purifying DNA fragments by agarose gel electrophoresis currently restricts an elementary cycle to two days, adds procedural complexity and prohibits robotic automation.

Addressing these limitations, a simplified DNA assembly process is developed while preserving its exquisite compatibility with protein coding sequences and the iterative nature of iFLinkC mediated DNA assembly (**Fig. 1**). Notably, input DNA is now removed genetically by alternating antibiotic resistances and ccdB counter selection (Wang et al. 2014) (**Fig. 1**). In this way, the need for physically purifying DNA by agarose gel electrophoresis is obviated. Crucially, this reduces procedural complexity and shortens the duration of an elementary assembly cycle to one day. To realize iFLinkC-EZ, the 5’ restriction site is omitted while DNA constructs are assembled linearly 5’ to 3’ directly in a destination plasmid, termed pZ or pZHiX (**Fig. 1**). The latter comprise modified derivatives of pPro24 (Lee and Keasling 2005) and pET (Shilling et al. 2020) where type II restriction sites relevant for iFLinkC-EZ mediated assembly have been removed (**Supporting Information**). Further, for greater efficiency and reliability, BsrDI is also exchanged for SapI/BspQI compared to the original iFLinkC assembly process.

### Experimental Implementation of iFLinkC-EZ

For its practical implementation, iFLinkC-EZ was tested under different experimental conditions and application scenarios. First, the efficiency of an elementary assembly cycle was tested in a conventional, manually handled DNA assembly reaction (**Fig. 2A**). The efficiency and fidelity of a DNA assembly reaction is generally high yielding hundreds of colonies per 5 µL assembly reaction with 4 out 4 samples displaying the correct sequence. Crucially, no background colonies were observed when T4 DNA ligase was omitted as no functional output plasmid was formed in the first place and all input plasmids were effectively degraded by T5 exonuclease and stringently counter selected (**Fig. 2B**). Further, due to greatly reduced procedural complexity and the absence of an agarose gel purification step, an elementary iFLinkC-EZ DNA assembly can be fully automated on a robotic platform (**Fig. 2C** and **Fig. 3**). In this case, 10-fold diluted enzymes was employed to reduce viscosity of the enzyme stocks that were dispensed using an I.DOT nanodispenser. Crucially, no tradeoffs in the stringency of DNA assembly are observed demonstrating the robustness, scalability and cost-efficiency of iFLinkC-EZ.

**Figure 2:**
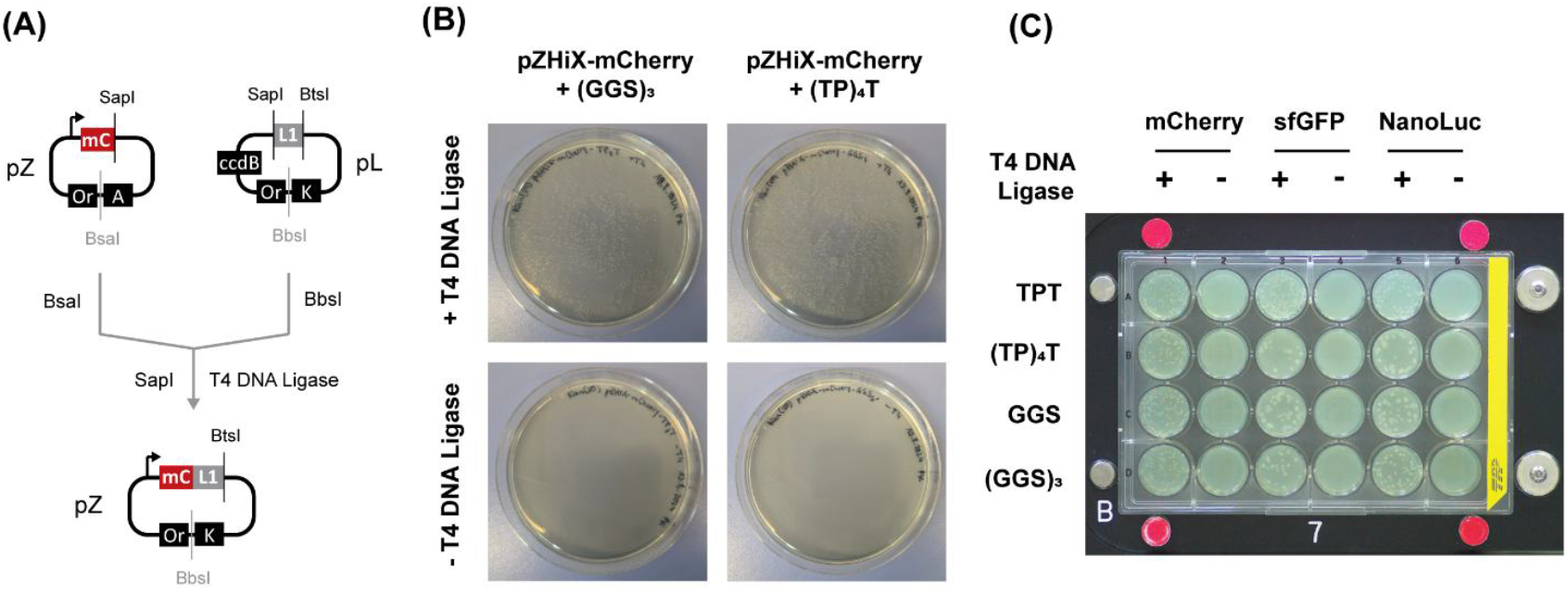

**(A)** Schematic of an elementary iFLinkC-EZ assembly cycle exemplified for a pZHiX and pL assembly step. The two input plasmids are separately digested using BsaI and BbsI before **(i)** the pL restriction digest is heat inactivated, **(ii)** the restriction digests pZHiX and pL combined and **(iii)** SapI and T4 DNA ligase added to complete an elementary DNA assembly step while any remaining DNA fragments are degraded by T5 exonuclease. Demonstrated efficiency and fidelity of an elementary iFLinkC-EZ assembly cycle for **(C)** manually-handled and **(C)** automated iFLinkC-EZ DNA assembly reactions. Transformation efficiencies constitute representative results and typically yield several hundred to thousands of colonies for a 5 µL manually-handled DNA assembly reaction, and tens of colonies for an automated DNA assembly reaction with 10× diluted enzyme dispensed with an I.DOT nanodispenser. In either case, no background colonies were observed when T4 DNA ligase was omitted. Detailed experimental protocols including sequences of pD, pL and pZ are enclosed in the Supporting Information.

**Figure 3:**
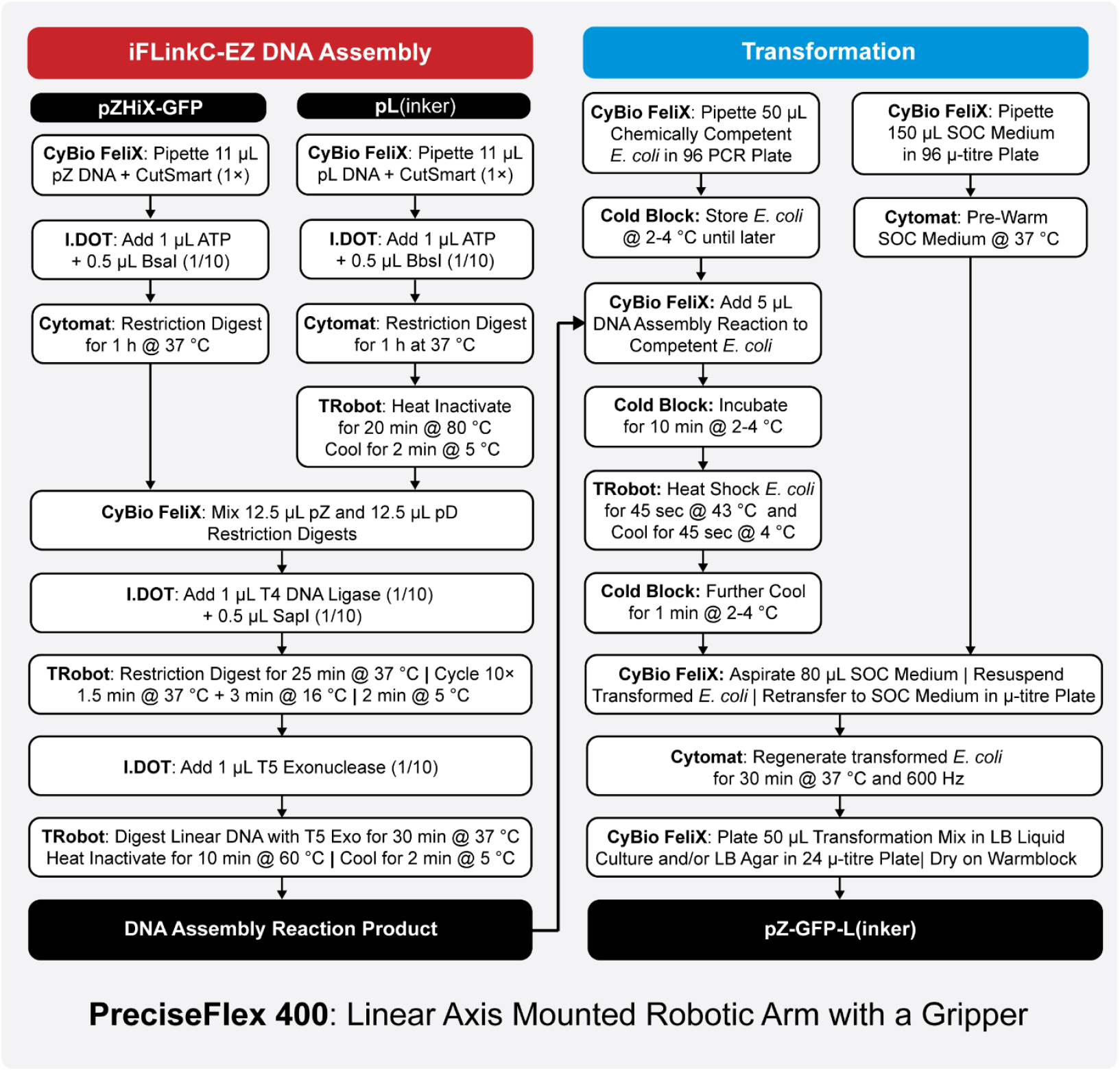

Schematic overview of an automated iFLinkC-EZ DNA assembly cycle exemplified for a pZ and pL recombination event. All devices are fully integrated and functionally connected through a linear axis mounted robotic arm equipped with a gripper (PreciseFlex 400) and orchestrated through the LARAsuit control software. A CyBio FeliX and an I.DOT are available for dispensing µL and nL volumes, respectively. A Cytomat is used to incubate the primary restriction digests at 37 ^°^C and regenerate transformed *E. coli* at 37 ^°^C after heat shock transformation. A Biometra TRobot is used for temperature cycling during the combined restriction-digestion-ligation step, the T5 exonuclease digest of linear DNA and heat shock of *E. coli* for transformation. A cold block cooled to 2-4 ^°^C is used for storing chemically competent *E. coli* and further cooling *E. coli* shortly after heat shock transformation.

### Iterative Assembly of Large Repetitive DNA Constructs with iFLinkC-EZ

Next, iFLinkC-EZ was assessed in the assembly of poly-fluorescent fusion proteins (FPs) which carry potential applications as fluorescent protein markers, labels and sensor. Yet, their assembly poses several technical challenges: In particular, from a technical perspective, the large homology of multiple identical copies of highly similar genes constitutes an exquisite challenge for any DNA assembly endeavor. Further, from a functional perspective, it is unclear how the identity of linkers impacts the functional expression of poly-fluorescent fusion proteins. In terms of linkers, we focus on a comparison of conventional poly-GGS and poly-PT linkers as the latter were recently shown to possess favorable biophysical properties compared to GGS-rich linkers in the context of Zn^2+^ responsive dual read-out FRET-BRET sensors (Gräwe et al. 2022).

To demonstrate the potency of iFLinkC-EZ, a set of monochromatic poly-fluorescent proteins (FP) composed of up to five copies of mCherry (Shaner et al. 2004; Carroll et al. 2014; Fages-Lartaud et al. 2022) was assembled by means of iFLinkC-EZ (**Fig. 4A**). Individual FPs were connected either by (GGS)_3_- or (TP)_4_T-linkers. The assembly of up to five copies of mCherry was achieved with high fidelity irrespective of the underlying FP or linker highlighting the potency of iFLinkC-EZ in the assembly of challenging complex fusion proteins (**Fig. 4B** and **Tab. S2**). Subsequent recombinant expression in BL21(DE3) *E. coli* yielded a red fluorescent signal for all mCherry constructs (**Fig. 4C**). With total fluorescence comparable or even slightly higher for the shortest constructs, this suggests that expression yields are limited by the total expression capacity of the cell rather than protein copies.

**Figure 4:**
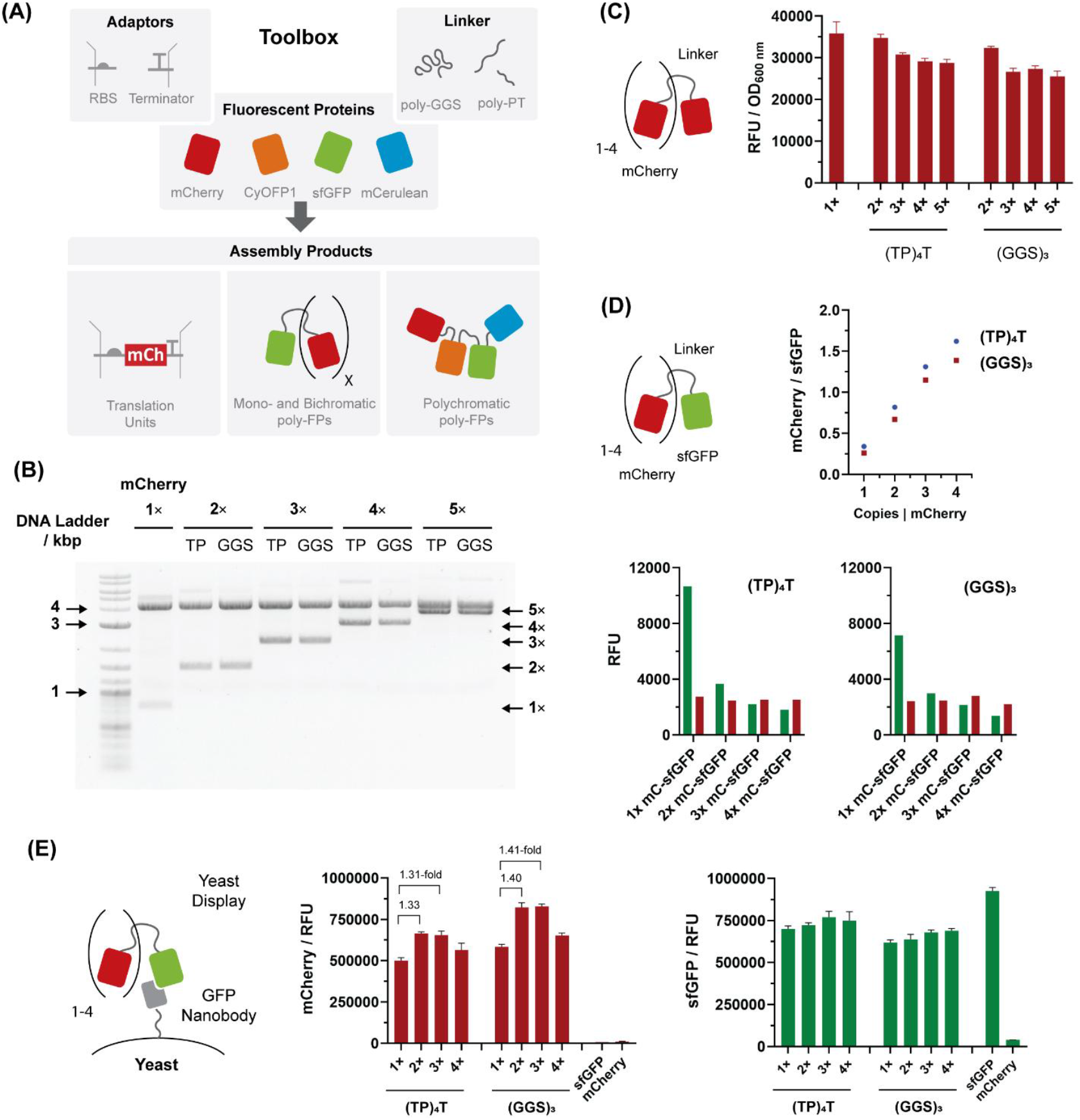

DNA assembly, recombinant expression and functional analysis of poly-fluorescent fusion proteins: **(A)** iFLinkC-EZ can be employed to assemble a wide range of different types of DNA constructs including fusion proteins, transcription and translation units based on modular parts; **(B)** The DNA assembly of poly-mCherry fusion proteins is stringent as judged by analytical DNA agarose gel electrophoresis of iteratively added copies of mCherry; **(C)** Spectrophotometric analysis of expressing poly-mCherry fusion proteins; **(D)** Spectrophotometric analysis of expressing mCherry_1-4_-sfGFP fusion proteins; **(E)** Antigen labelling using mCherry_1-4_-sfGFP fusion proteins (directly applied from cells lysates) in yeast display. sfGFP served as the antigen in order to label sfGFP specific nanobodies. The fluorescence per GFP-specific nanobody displaying yeast cells reaches a maximum for two copies of mCherry before leveling for three and dropping for four copies of mCherry.

While the total fluorescent signal gives an approximate indication for the functional expression of poly-FPs, it does not account for the efficiency of full-length protein synthesis. Namely, fluorescence could equally arise from pre-maturely truncated protein expression. Conversely, it is not known to what extent all copies of mCherry are functionally folded and thus contribute to fluorescence. To address this, sfGFP (Pédelacq et al. 2006) was fused to the C-terminus of up to four copies of mCherry and recombinantly expressed in HMS174(DE3). As before, both (GGS)_3_- or (TP)T_4_ linkers were used to connect individual FPs. Endpoint analysis after recombinant expression shows that the sfGFP signal drops with an increasing number of mCherry copies (**Fig. 4D**). While this trend can be accounted for by stoichiometry, the ratio of sfGFP and mCherry dependent fluorescence turned out greater with more favorable ratios for (GGS)_3_ compared to (TP)T_4_ linkers (**Fig. 4D**). This suggests an increasing, albeit small proportion of truncated protein contributes to the red fluorescent signal as the size of the poly-fluorescent fusion protein increases while (GGS)_3_ linkers yield slightly more functional full-length proteins compared to (TP)_4_T linkers. To dissect this further and demonstrate their functionality, poly-mCherry-sfGFP were used to label yeast cells displaying sfGFP-specific nanobodies (Rothbauer et al. 2006) *via* Aga2p fusion proteins on their surface (Chao et al. 2006) (**Fig. 4E**). To ensure comparable concentrations of sfGFP antigen, cell lysates were normalised over sfGFP fluorescence before they were applied to yeast cells. Yeast cells could be efficiently labelled with poly-mCherry-sfGFP as judged by single cell flow cytometric analysis (**Fig. 4E**). Notably, fluorescence increases for up to two copies of mCherry before leveling for three and dropping with a fourth copy of mCherry (**Fig. 4E**) which could either be due to fewer mCherry functionally maturing or a reduced binding affinity for larger poly-FP constructs. Further, binding was dependent on the display of a sfGFP-specific nanobody as displaying a non-specific binder did not yield a fluorescent signal (**Fig. S1**).

### Extending iFLinkC-EZ with a Capacity for Golden Gate Assembly

Compared to the original implementation, iFLinkC-EZ is restricted to linear assembly modes – i.e. for larger multi-gene expression constructs, the advantages of a technically simpler and especially shorter assembly cycle afforded by iFLinkC-EZ are mitigated by the greater number of steps imposed by a linear assembly scheme. To overcome these limitations, iFLinkC-EZ can be rendered compatible with alternative DNA assembly methods following the introduction of distinct adaptor modules that code for dedicated DNA recombination sites. To this end, translation units are first assembled individually with iFLinkC-EZ before being compiled into larger multi-gene expression constructs either based on homologous (Bomfiglio et al. 2025) and non-homologous DNA recombination (Laborda-Mansilla and García-Ruiz 2025).

For instance, additional sites for the type IIS restriction enzymes BsmBI can convey a capacity for Golden Gate assembly and thus assist in the construction of large poly-cistronic constructs expressing multiple FPs (**Fig. 5A**). To this end, individual translation units coding for different FPs were first assembled with iFLinkC-EZ and featured **(i)** a translation initiation module with a ribosome binding site (RBS) (Iverson et al. 2016), **(ii)** the protein coding sequence for mCherry (Shaner et al. 2004), CyOFP1 (Chu et al. 2016), sfGFP (Pédelacq et al. 2006) or mCerulean3 (Rizzo and Piston 2005), and **(iii)** a translation termination with a stop codon. Fluorescent proteins generally featured identical RBS. Crucially, translation initiation and translation termination modules feature a positioning module with a defined 4-base overhang generated upon restriction digestion with BsmBI in order to direct the assembly of individual translation units into a poly-cistronic expression construct. In representative results, the assembly efficiency was high yielding >50 colonies per 5 µL assembly reaction (**Fig. 5B**) with 2 out of 3 constructs displaying the correct sequence as confirmed by sequencing.

**Fig 5:**
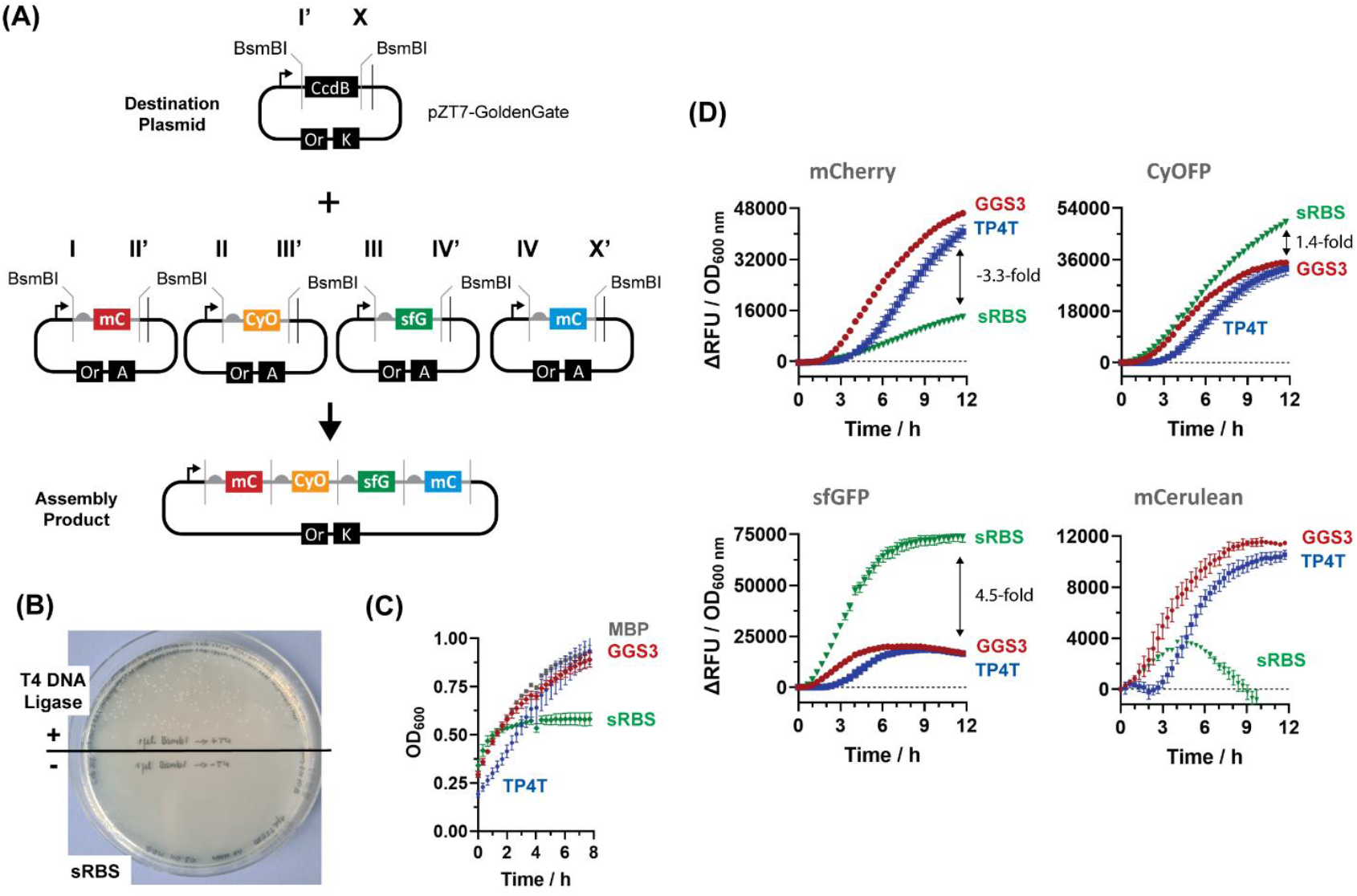

**(A)** iFLinkC-EZ mediated parallel assembly of multiple translation units into a poly-cistronic expression construct. Dedicated adaptor modules provide additional sites for type IIS restriction digestion by BsmBI. The resultant 4-base overhangs subsequently direct the assembly of individual translation units into a poly-cistronic expression constructs *via* Golden Gate. Adaptors can be introduced as independent modules or combined with functional modules for translation initiation and translation termination; **(B)** Successful transformation of a poly-cistronic multi-gene expression cassette coding for multiple FPs *via* BsmBI mediated Golden Gate assembly using adaptor modules. The negative control features no T4 DNA ligase; **(C)** Poly-chromatic FPs display differential growth in HMS174(DE3). Compared to a MBP control and a poly-fluorescent protein (GGS)_3_ construct, the poly-cistronic construct initially displays comparable growth rates before moving early into stationary phase. Conversely, the poly-FP fusion proteins separated by (TP)_4_T linkers initially displays slower growth but ultimately reach comparable OD_600_ compared to (GGS)_3_ and the MBP control; **(D)** Functional expression of poly-FP poly-cistronic expression cassette and its comparison with an equivalent poly-FP fusion protein. A poly-fluorescent fusion protein construct displays more even expression levels compared to a poly-cistronic expression construct as judged by the relative expression levels for CyOFP1, sfGFP and mCerulean3 expressed from identical RBSs.

Further, comparing the expression of FPs in the context of a poly-cistronic construct and poly-fusion proteins separated by (GGS)_3_- and (TP)_4_T-linkers yielded the following observations: First of all, constructs turned out functional as up to 4 different FPs could be expressed in *E. coli* (**Fig. 5D**). However, relative expression levels turn out substantially more variable for poly-cistronic constructs compared to poly-fluorescent fusion proteins: i.e. the fluorescence for mCherry, CyOFP1 and sfGFP respectively turns out 3.3-fold lesser, 1.4-fold greater and 4.5-fold greater in the context of a poly-cistronic construct compared a poly-fluorescent fusion protein. Further, mCerulean3 is only marginally expressed in the poly-cistronic construct. Thus, care must be taken when balancing the expression of multiple proteins in the context of a poly-cistronic expression construct as identical RBS do not necessarily ensure comparable expression levels. Secondly, constructs displayed varying levels of toxicity: To start with, the functional analysis was restricted to the K12 derivative HMS174(DE3) as RBS separated poly-cistronic construct and (TP)_4_T but not the (GGS)_3_ turned out toxic upon transformation into the B strain derivative BL21(DE3). Further, toxicity also reflected by variable growth rates in the K12 strain derivative HMS174(DE3) as the poly-cistronic construct pre-maturely reached stationary phase and the poly-(TP)_4_T construct showed delayed growth before reaching stationary phase (**Fig. 5C**). Given the high degree of sequence identity and similarity in function as well as the fact this was not observed for poly-mCherry constructs, this appears to be an idiosyncratic feature of the resultant expression constructs suggesting that even comparatively small changes can potentially impact cell growth.

## Conclusions

With iFLinkC-EZ, a simplified and automatable method for protein-centered DNA assembly is developed. Crucially, it preserves key features of the original iFLinkC DNA assembly algorithm as protein coding sequences can be iteratively and arbitrarily recombined independent of orientation, length and sequence complexity in order to build complex fusion proteins. Key advances compared to the original iFLinkC implementation include shortening the DNA assembly cycle to one day along with a significantly reduced procedural complexity, lower operator skill and a capacity for robotic automation. This is achieved by genetically purifying the output plasmid DNA by exploiting a change in antibiotic resistance and counter selection with ccdB which obviates the need DNA agarose gel electrophoresis. Crucially, this results in a very high stringency of DNA assembly under a variety of experimental scenarios and also confers a capacity for robotic automation. Notably, the material cost of an elementary iFLinkC-EZ assembly cycle in a volume of 12.5 µL equates to 2.5 - 4 EUR for a manually-handled DNA assembly reaction depending whether a pD or pL module is fused to pZ (**Supporting Information**). Costs can be further reduced to less than 1 EUR per assembly step by diluting enzymes 10-fold as required by I.DOT nanodispenser during robotic automation.

In proof-of-concept, the potency of iFLinkC-EZ was demonstrated in the assembly of monochromatic and poly-chromatic poly-fluorescent fusion proteins while testing the impact of GGS- and poly-TP linkers on their underlying functional expression. Applications are demonstrated in labelling surface displayed antigens on yeast cells. Despite the highly repetitive nature of poly-fluorescent fusion proteins, full length protein expression is readily achieved. Otherwise, the impact of (GGS)_3_ and (TP)_4_T linkers is comparatively small as (GGS)_3_ linkers display slightly more favorable properties compared to (TP)_4_T linkers both in terms of full-length expression in lysates and subsequent labelling in yeast cells. Therefore, the main difference between poly-GGS and poly-TP linkers can be attributed to their biophysical properties as the latter better separates two domains as demonstrated in the context of Zn^2+^-responsive dual read-out FRET-BRET sensor (Gräwe et al. 2022).

Finally, iFLinkC-EZ – which is especially useful for compiling (complex) fusion proteins and translational units – can be readily combined with alternative DNA assembly frameworks in order to build large poly-cistronic expression constructs. This can be readily achieved by introducing dedicated adaptors during an iFLinkC-EZ mediated assembly process. Depending on the expression construct, adaptors can either be introduced on their own or be integrated with functional modules, for instance, to initiate and terminate translation and potentially transcription as well. Such a capacity is experimentally demonstrated for Golden Gate assembly with dedicated adaptors coding for additional type IIS restriction sites. Proof-of-concept is achieved in the expression of poly-chromatic fluorescent proteins. Notably, despite identical RBS preceding individual FPs, their functional expression turns out substantially more variable compared to a poly-chromatic fluorescent fusion protein indicating strong context dependencies when expressing proteins from a poly-cistronic construct. Beyond Golden Gate, alternative DNA assembly strategies, including homology-based DNA recombination methods, are equally conceivable and can be readily enabled by introducing dedicated adaptors adding to the ease, flexibility and interoperability of iFLinkC-EZ mediated assembly further adding to its general utility.

## Materials and Methods

A detailed account of the materials including sequences of DNA constructs and experimental methods used in this study are provided in the Supporting Information. iFLinkC-EZ was automated on the CompuGene robotic platform (Spannenkrebs et al. 2025) and orchestrated using the LARAsuit control software (https://gitlab.com/LARAsuite).

## Acknowledgements

The authors like to thank Mark Doerr, University of Greifswald, for assistance with the implementation LARAsuit on the CompuGene robotic platform and Beatrix Suess for access to cytometry equipment.

