## Supporting Information for "iFLinkC-EZ: A scalable and automatable method for the assembly of complex fusion proteins and multi-gene expression constructs based on the iFLinkC framework"

### Materials and Methods

*General*

DNA assembly reactions for iFLinkC-EZ were chemically transformed in DH10B and protein expression conducted in either BL21(DE3) and HMS174(DE3) as indicated. Lysogeny broth (LB) was supplemented with the relevant antibiotics as indicated. Kanamycin was used at a concentration of 50 µg/mL and ampicillin at a concentration of 100 µg/mL. DNA plasmids were purified using commercial DNA purification kits (Macherey & Nagel). Assembled DNA constructs were sequence verified using a commercial provider (Microsynth DNA). The DNA used to redesign plasmid backbones (e.g. pD, pL, pZHiX and pZT7-Golden Gate) and the component parts (e.g. linker elements, functional domains and adaptor modules) were commercially sourced (Sigma-Aldrich, Integrated DNA Technologies) before being subcloned into suitable input plasmids. To render plasmid backbones and components compatible with different assembly modes, any restriction sites for BtsI, SapI, BsaI and BpsI required for iFLinkC-EZ mediated assembly and restriction sites BsmbI required for Golden Gate mediated assembly were removed from the DNA sequences and replaced with synonymous codons by TEDA (Xia et al. 2019). Plasmids pD and pL constitute modified of pPro24 rendered compatible with iFLinkC-EZ mediated DNA assembly (Lee and Keasling 2005). Plasmid pZHiX and pZT7-Golden-Gate is a modified derivative of pET (Shilling et al. 2020). The detailed DNA sequences of different plasmids are given below. Reagents for iFLinkC-EZ – i.e. different restriction enzymes, T4 DNA ligase and T5 exonuclease were generally purchased from New England Biolabs.

*Elementary iFLinkC-EZ Assembly Cycle*

iFlinkC-EZ assembly is based on two different types of assembly reactions as either pD or pL are iteratively fused to pZHiX (**Tab. S1**). The two assembly reactions are denoted as Version 1 and Version 2 and require input plasmids to be treated with different combinations of restriction enzymes before chemically transformed *E. coli* are plated on LB agar supplemented with different antibiotics depending on the resistance genes in the respective plasmids – i.e. plasmids harboring KAN resistances (i.e. pL-vectors, the original pZ destination vector, or pZ-vectors with a 3' terminal linker element) are strictly restriction digested with BbsI-HF. Conversely, plasmids harboring AMP resistances (e.g. pD-vectors, or pZ-vectors with a 3' terminal functional domain) are always restriction digested using BsaI-HF. Further, when a functional domain is added to a pZ destination vector (e.g. the original pZ destination vector or a pZ destination vector with a 3' terminal linker element) both plasmids are cleaved using BtsI-HF (Version 1). Conversely, when a linker element is added to the pZ destination vector both plasmids are cleaved using SapI (Version 2). A step wise experimental protocol is outlined on the following pages. For manually handled reactions, volumes can either comprise 12.5 µL or 25 µL using undiluted enzymes.

*Cost breakdown for iFLinkC-EZ assembly reactions*

The primary costs for an elementary iFLinkC-EZ assembly cycle stem from **(i)** commercial plasmid preparation kits used to prepare input plasmids (e.g. 2× 0.68 EUR for a commercial plasmid purification kit from Macherey & Nagel for purifying pZ and either pD or pL) and **(ii)** enzymes (e.g. per 12.5 µL reaction: BsaI-HF @ 0.30 EUR, BbsI-HF @ 0.13 EUR, T4 DNA ligase @ 0.29 EUR, T5 exonuclease @ 0.15 EUR and either BtsI-HF @ 0.32 EUR or SapI @ 0.15 EUR for a pD or pL fusion, respectively). Further, for **(iii)** self-made competent cells, 1 EUR is calculated per transformation. Overall, the material costs for an elementary assembly cycle thus approximate 3.50-5.00 EUR depending whether iFLinkC-EZ is performed in a volume of 12.5 µL or 25 µL.

*iFLinkC-EZ Step-by-Step Protocol*

- Depending whether pD or pL is attached to the growing destination plasmid pZHiX, two separate combinations of restrictions digests (Version 1 or Version 2) are prepared:

**Table S1.** Reaction parameters for adding either a pD or pL vector

**Version 1 (→ pZHiX with pD later to be plated on ampicillin)**

| **pZHiX-Vector (Restriction Digest #1)** | **pD-Vector (Restriction Digest #2)** |
|---|---|
| 100 ng DNA | 500 ng DNA |
| 2.5 µL CutSmart Buffer | 2.5 µL CutSmart Buffer |
| 2.5 µL ATP (10 mM) | 2.5 µL ATP (10 mM) |
| 0.5 µL BbsI-HF | 0.5 µL BsaI-HF |
| Fill up to 25 µL $ddH_2O$ | Fill up to 25 µL $ddH_2O$ |

**Version 2 (→ pZHiX with pL later to be plated on kanamycin)**

| **pZHiX-Vector (Restriction Digest #1)** | **pL-Vector (Restriction Digest #2)** |
|---|---|
| 100 ng DNA | 500 ng DNA |
| 2.5 µL CutSmart Buffer | 2.5 µL CutSmart Buffer |
| 2.5 µL ATP (10 mM) | 2.5 µL ATP (10 mM) |
| 0.5 µL BsaI-HF | 0.5 µL BbsI-HF |
| Fill up to 25 µL $ddH_2O$ | Fill up to 25 µL $ddH_2O$ |

- The two restriction digests are then incubated for 30 min to 60 min at 37°C. Then the reaction restriction digesting either pD (Version 1) or pL (Version 2) is inactivated at 80°C for 20 min – i.e. the reaction which restriction digests pZ remains active.
- The contents of both restriction digests are then mixed to a total volume of 50 µL and 0.5 µL T4 DNA Ligase is added together with either 0.5 µL BtsI-HF (Version 1) or 0.5 µL of SapI (Version 2) before being subject to the following temperature cycling protocol.

| **Temperature** | **Duration** | **Number of cycles** |
|---|---|---|
| 37 °C | 25:00 min | 1 |
| 37 °C | 1:30 min | 10 |
| 16 °C | 3:00 min | |

- Upon completion of temperature cycling, 0.5 µL of T5 exonuclease is added to the assembly reaction. This step is optional but removes any remaining linear DNA fragments and thus reduces any background which may potentially arise from unligated plasmid or thereof based recombinant products.

| | | |
|---|---|---|
| 37 °C | Pause | Add 0.5 µL of T5 Exonuclease |
| 37 °C | 30:00 min | 1 |
| 65 °C | 10:00 min | 1 |

- Afterwards, 5 µL of the reaction mixture is used to transform 50 µL of chemically competent *E. coli* cells
- Transformed *E. coli* are then plated on LB agar plates containing the appropriate antibiotic and left to incubate overnight at 37 °C – i.e. the antibiotic comprises **ampicillin** when adding a functional domain through recombination *via* pD (**Version 1**), and **kanamycin** when adding a linker element through recombination *via* pL (**Version 2**)
- The following day, after an efficient transformation with >100 colonies per plate, cells are harvested by rinsing agar plates with 2 mL dd$H_2O$ while scrubbing colonies off using a Drigalski spatula. It is recommended that a portion of the resulting cell suspension is used to prepare glycerol stocks and thus store assembly intermediates.

- Then, cells are sedimented by centrifugation for 1 min at 11,000 g and the newly recombined pZHiX is extracted with a plasmid purification kit. The resultant extracted plasmid can then serve as an input DNA for the next iFLinkC-EZ assembly cycle.

Alternatively, for further cost reduction, iFLinkC-EZ with 10-fold diluted enzymes as required when using an I.DOT nanodispenser. This readily saves costs and renders iFLinkC-EZ compatible with robotic automation. Schematic workflows for **(i)** iFLinkC-EZ mediated assembly along with **(ii)** transformation on the robotic platform are depicted (see **Fig. 3**).

**Table S2.** Summary of DNA fragment sizes generated using a BamHI / SalI analytical restriction digest with BamHI and SalI of poly-fluorescent protein constructs in Figure 3B

| Lane | DNA Construct | DNA Fragments Sizes |
|---|---|---|
| 1 | pZHiX-$His_6$-$TVMV_{cs}$-mCherry | 4253 bp + 746 bp |
| 2 | pZHiX-$His_6$-$TVMV_{cs}$-[mCherry-$(TP)_4T]_1$-mCherry | 4253 bp + 1484 bp |
| 3 | pZHiX-$His_6$-$TVMV_{cs}$-[mCherry-$(TP)_4T]_2$-mCherry | 4253 bp + 2222 bp |
| 4 | pZHiX-$His_6$-$TVMV_{cs}$-[mCherry-$(TP)_4T]_3$-mCherry | 4253 bp + 2960 bp |
| 5 | pZHiX-$His_6$-$TVMV_{cs}$-[mCherry-$(TP)_4T]_4$-mCherry | 4253 bp + 3698 bp |

*Functional Expression of poly-FP Fusion Proteins in Microtitre Plates*

Chemically competent *E. coli* BL21(DE3) were transformed with fully assembled expression constructs – i.e. $pZ^{Kan}$ plasmids coding for monochromatic and polychromatic FP fusions as indicated – before being plated on LB agar plates (+ KAN) and incubated overnight at 37 °C. Single colonies were then used to inoculate 5 mL LB medium (+ KAN) and grown overnight at 37 °C by shaking at 180 rpm. The resultant preculture was then used to inoculate a 96-well plate with 200 µL LB medium (+ KAN) to an $OD_{600}$ 0.05 before being shaken at 37 °C for 3 hours. Protein expression was then induced with 0.5 mM IPTG and the culture further shaken at 30 °C in TECAN Spark while measuring the fluorescence of mono- and polychromatic fluorescent fusion proteins (mCherry: ex. 575 nm, em. 620 nm).

*Expression of poly-Fluorescent Fusion Proteins for Yeast Display Selection*

Chemically competent *E. coli* HMS174(DE3) were transformed with pZHiX expressing either coding mono- or poly-fluorescent fusion proteins by heat-shock. Cells were plated on LB agar plates (+ 100 µg/mL AMP) and incubated overnight at 37 °C. To prepare proteins for yeast display selections, single colonies were used to inoculate 5 mL LB medium (+ 100 µg/mL AMP) and incubated overnight at 37 °C and 200 rpm. The pre-culture was then used to inoculate 50 mL LB (+ 100 µg/mL AMP) to an initial $OD_{600}$ 0.05 and incubated at 37 °C and 200 rpm. When the $OD_{600}$ reached 0.6, protein expression was induced with 0.5 mM IPTG and cells were incubated overnight at 30 °C and 200 rpm. The following day cells were harvested and washed twice in ice-cold 1× PBS. Cells were then resuspended in ice-cold 1× PBS to reach an $OD_{600}$ equivalent of 10. Aliquots of 500 µL of cell suspension were lysed by sonification (QSonica Q125) using the following intensity: 50 % amplitude, 10 sec pulses with 10 sec breaks for 1 min of active sonication). Cell debris was removed by centrifugation (2 min, 17.000 g at room temperature) the supernatant was stored on ice in a new reaction tube until further application to yeast cells.

*Yeast Display with poly-Fluorescent Fusion Proteins*

Yeast display selections were performed according to previous protocols plates (Chao et al. 2006). Chemically competent yeast cells (EBY100) were transformed with plasmids pCTCon2-Aga2P-scFv and pCTCon2-Aga2P-GFP_nb using the Frozen-EZ Yeast Transformation II kit (Zymo Research) according to manufacturer's instructions. Cells were plated on SCDAA agar and incubated for 2 days at 30 °C. Single colonies were picked and used to inoculate 2 mL of SCDAA medium. Pre-cultures were incubated overnight at 30 °C and 200 rpm. The next day, 10 mL of SDCAA medium was inoculated to an $OD_{600}$ of 0.05. Cells were incubated for 6 h at 30 °C and 200 rpm. Induction of gene expression through medium exchange: Cells were harvested at 2,500 g for 5 min and resuspended in SGCAA medium. Cells were incubated at 30 °C, 200 rpm for 2 days. Yeast cells were harvested and washed and resuspended in 1× PBSF. 50 µL of yeast cells were mixed with 50 µL of *E. coli* lysates and incubated for 20 min at 30 °C and 300 rpm. Cells were harvested at 2,500 g for 5 min and washed twice with ice-cold 1× PBSF. Cells were resuspended in 200 µL ice-cold, sterile-filtrated 1×PBSF and then diluted 1:20 in the same buffer in black bottom 96 well plates. The fluorescent staining efficiency of yeast cells was analysed by flow cytometry (CytoFlex S, Beckman Coulter). sfGFP was excited using a 488 nm laser and emission was filtered through a 525/40 nm filter, while mCherry was excited using a 561 nm laser and emission was filtered through a 610/20 filter. Cells were analysed using CytExpert (Beckman Coulter) and gates were set as to only include singlet yeast cells. Mean values and standard deviations were calculated from triplicates.

**Aga2P-GFP_nb**

Aga2P, Factor Xa site, HA epitope tag, GFP_nb, Myc epitope tag

MQLLRCFSIFSVIASVLAQELTTICEQIPSPTLESTPYSLSTTTILANGKAMQGVFEYYKSVTFVSNCGSHPSTTSKGSPIN
TQYVFKDNSSTIEGRYPYDVPDYALQASGGGGSGGGGSGGGGSASQVQLVESGGALVQPGGSLRLSCAASGFPVNRYSMRWY
RQAPGKEREWVAGMSSAGDRSSYEDSVKGRFTISRDDARNTVYLQMNSLKPEDTAVYYCNVNVGFEYWGQGTQVTVSSGSEQ
KLISEEDL

**Aga2P-scFv (non-specific vector control)**

Aga2P, Factor Xa site, HA (human influenza hemagglutinin) epitope tag, scFv, Myc epitope tag

MQLLRCFSIFSVIASVLAQELTTICEQIPSPTLESTPYSLSTTTILANGKAMQGVFEYYKSVTFVSNCGSHPSTTSKGSPIN
TQYVFKDNSSTIEGRYPYDVPDYALQASGGGGSGGGGSGGGGSASCGGGGTSKISHFLKMESLNFIRAHTPYINIYNCEPAN
PSEKNSPSTQYCYSIQSSQVDCGGGSEQKLISEEDL

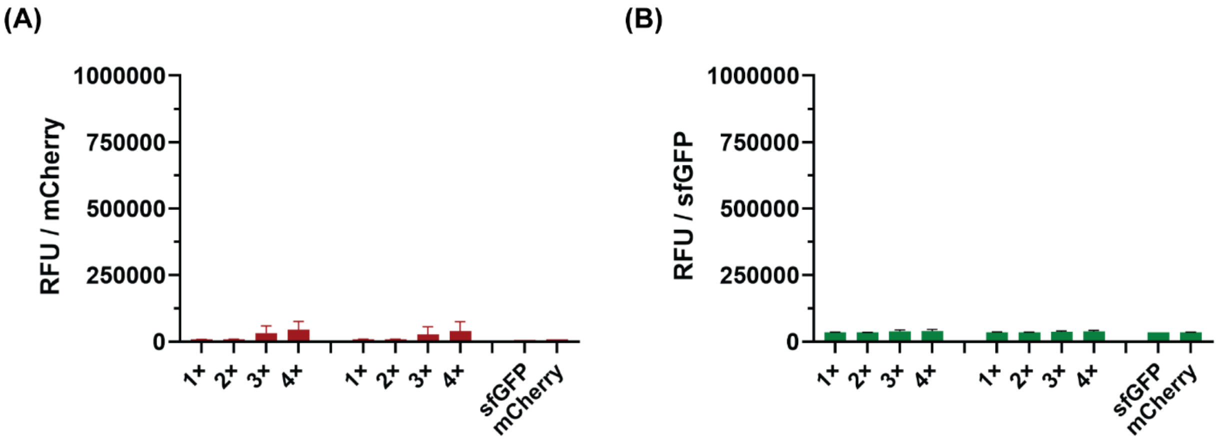


**Fig. S1:** Testing for non-specific binding of poly-fluorescent fusion proteins in yeast display. Summary of fluorescence following labelling yeast cells displaying a non-specific scFv with mCherry$_{1\text{-}4}$-sfGFP fusion proteins (directly applied from cell lysates). Neither **(A)** GFP nor **(B)** mCherry dependent fluorescence can be detected.

### Summary of DNA Sequences for iFLinkC-EZ Mediated DNA Assembly

**pZHiX-6×His-TVMV$^{CS}$-ccdB |** 6×His, BtsI, ccdB, KanR, **BbsI**, ColE1 ori, LacI

GGTGCATGCAAGGAGATGGTAGAGGATCGAGATCTCGATCCCGCGAAATTAATACGACTCACTATAGGGAGAGGAATTGTGA
GCGGATAACAATTCCCCTCTAGAAATAATTTTGTTTAACTTTAAGAAGGAGAGCAGCTATGCAGCTTAGCCATCACCATCAT
CATCACAGCAGCGGATCCGAAACCGTGCGTTTCCAGTCTGGGCACTGCATGCAGTTTAAGGTTTACACCTATAAAAGAGAGA
GCCGTTATCGTCTGTTTGTGGATGTACAGAGTGATATTATTGACACGCCCGGGCGACGGATGGTGATCCCCCTGGCCAGTGC
ACGTCTGCTGTCAGATAAAGTCTCCCGTGAACTTTACCCGGTGGTGCATATCGGGGATGAAAGCTGGCGCATGATGACCACC
GATATGGCCAGTGTGCCGGTCTCCGTTATCGGGGAAGAAGTGGCTGATCTCAGCCACCGCGAAAATGACATCAAAAACGCCA
TTAACCTGATGTTCTGGGGAATATAAACTAGTGGTACCGCTGCTAACAAAGCCACAAAGCCCGAAAGGAAGCTGAGTTGGCT
GCTGCCACCGCTGAGCAATAACTAGCATAACCCCTTGGGGCCTCTAAACGGGTCTTGAGGGGTTTTTTGCTGAAAGGAGGAA
CTATATCCGGATTGGCGAATGGGGCCATAAACTGCCAGGCATCAAATTAAGCAGAAGGCCATCCTGACGGATGGCCTTTTTG
CGTTTCTACAAACTCTCTGATCCTTCAACTCAGCAAAAGTTCGATTTATTCAACAAAGCCACGTTGTGTCTCAAAATCTCTG
ATGTTACATTGCACAAGATAAAAATATATCATCATGAACAATAAAACTGTCTGCTTACATAAACAGTAATACAAGGGGTGTT
ATGAGCCATATTCAACGGGAAACGTCTTGCTCTAGGCCGCGATTAAATTCCAACATGGATGCTGATTTATATGGGTATAAAT
GGGCTCGCGATAATGTCGGGCAATCAGGTGCGACAATCTATCGATTGTATGGGAAGCCCGATGCGCCAGAGTTGTTTCTGAA
ACATGGCAAAGGTAGCGTTGCCAATGATGTTACAGATGAGATGGTCAGACTAAACTGGCTGACGGAATTTATGCCTCTGCCG
ACCATCAAGCATTTTATCCGTACTCCTGATGATGCATGGTTACTCACCACGGCGATCCCAGGGAAAACAGCATTCCAGGTAT
TAGAAGAATATCCTGATTCAGGTGAAAATATTGTTGATGCGCTGGCGGTGTTCCTGCGCCGGTTGCATTCGATTCCTGTTTG
TAATTGTCCTTTTAACAGCGACCGCGTATTTCGTCTGGCTCAGGCGCAATCACGAATGAATAACGGTTTGGTTGATGCGAGT
GATTTTGATGACGAGCGTAATGGCTGGCCTGTTGAACAAGTCTGGAAAGAAATGCATAAACTTTTGCCATTCTCACCGGATT
CAGTCGTCACTCATGGTGATTTCTCACTTGATAACCTTATTTTTGACGAGGGGAAATTAATAGGTTGTATTGATGTTGGACG
AGTCGGAATCGCAGACCGATACCAGGATCTTGCCATCCTATGGAACTGCCTCGGTGAGTTTTCTCCTTCATTACAGAAACGG
CTTTTTCAAAAATATGGTATTGATAATCCTGATATGAATAAATTGCAGTTTCATTTGATGCTCGATGAGTTTTTCTAACTGT
CAGACCAAGTTTACTCATATATACTTTAGATTGATTT**GAAGAC**TACGCGCCCTGTAGCGGCGCATTAAGCGCGGCGGGTGTG
GTGGTTACGCGCAGCGTGACCGCTACACTTGCCAGCGCCCTAGCGCCCGCTCCTTTCGCTTTCTTCCCTTCCTTTCTCGCCA
CGTTCGCCGGCTTTCCCCGTCAAGCTCTAAATCGGGGGCTCCCTTTAGGGTTCCGATTTAGTGCTTTACGGCACCTCGACCC
CAAAAAACTTGATTTGGGTGATGGTTCACGTAGTGGGCCATCGCCCTGATAGACGGTTTTTCGCCCTTTGACGTTGGAGTCC
ACGTTCTTTAATAGTGGACTCTTGTTCCAAACTGGAACAACACTCAACCCTATCTCGGGCTATTCTTTTGATTTATAAGGGA
TTTTGCCGATTTCGGCCTATTGGTTAAAAAATGAGCTGATTTAACAAAAATTTAACGCGAATTTTAACAAAATATTAACGTT
TACAATTTAAAAGGATCTAGGTGAAGATCCTTTTTGATAATCTCATGACCAAAATCCCTTAACGTGAGTTTTCGTTCCACTG
AGCGTCAGACCCCGTAGAAAAGATCAAAGGATCTTCTTGAGATCCTTTTTTTCTGCGCGTAATCTGCTGCTTGCAAACAAAA
AAACCACCGCTACCAGCGGTGGTTTGTTTGCCGGATCAAGAGCTACCAACTCTTTTTCCGAAGGTAACTGGCTTCAGCAGAG
CGCAGATACCAAATACTGTCCTTCTAGTGTAGCCGTAGTTAGGCCACCACTTCAAGAACTCTGTAGCACCGCCTACATACCT
CGCTCTGCTAATCCTGTTACCAGTGGCTGCTGCCAGTGGCGATAAGTCGTGTCTTACCGGGTTGGACTCAAGACGATAGTTA
CCGGATAAGGCGCAGCGGTCGGGCTGAACGGGGGGTTCGTGCACACAGCCCAGCTTGGAGCGAACGACCTACACCGAACTGA
GATACCTACAGCGTGAGCTATGAGAAAGCGCCACGCTTCCCGAAGGGAGAAAGGCGGACAGGTATCCGGTAAGCGGCAGGGT
CGGAACAGGAGAGCGCACGAGGGAGCTTCCAGGGGGAAACGCCTGGTATCTTTATAGTCCTGTCGGGTTTCGCCACCTCTGA
CTTGAGCGTCGATTTTTGTGATGCTCGTCAGGGGGGCGGAGCCTATGGAAAAACGCCAGCAACGCGGCCTTTTTACGGTTCC
TGGCCTTTTGCTGGCCTTTTGCTCACATGTTCTTTCCTGCGTTATCCCCTGATTCTGTGGATAACCGTATTACCGCCTTTGA
GTGAGCTGATACCGCTCGCCGCAGCCGAACGACCGAGCGCAGCGAGTCAGTGAGCGAGGAAGCGGTAGAGCGCCTGATGCGG
TATTTTCTCCTTACGCATCTGTGCGGGAGATCCCGGTGCCTAATGAGTGAGCTAACTTACATTAATTGCGTTGCGCTCATTG
ACCGCTTTCCAGTCGGGAAACCTGTCGTGCCAGCTGCATTAATGAATCGGCCAACGCGCGGGGAGAGGCGGTTTGCGTATTG
GGCGCCAGGGTGGTTTTTCTTTTCACCAGTGAGACGGGCAACAGCTGATTGCCCTTCACCGCCTGGCCCTGAGAGAGTTGCA
GCAAGCGGTCCACGCTGGTTTGCCCCAGCAGGCGAAAATCCTGTTTGATGGTGGTTAACGGCGGGATATAACATGAGCTGTC
CTCGGTATCGTCGTATCCCACTACCGAGATGTCCGCACCAACGCGCAGCCCGGACTCGGTAATGGCGCGCATTGCGCCCAGC
GCCATCTGATCGTTGGCAACCAGCATCGCGGTTGGAACGATGCCCTCATTCAGCATTTGCATGGTTTGTTGAAAACCGGACA
TGGCACTAAAGTCGCCTTCCCGTTCCGCTATCGGCTGAATTTGATTGCGAGTGAGATATTTATGCCAGCCAGCCAGACGCAG
ACGCGCCGAGACAGAACTTAATGGGCCCGCTAACAGCGCGATTTGCTGGTGACCCAATGCGACCAGATGCTCCACGCCCAGT
CGCGTACCATCTTCATGGGAGAAAATAATACTGTTGATGGGTGTCTGGTCAGAGACATCAAGAAATAACGCCGGAACATTAG
TGCAGGCAGCTTCCACAGCAATGGCATCCTGGTCATCCAGCGGATAGTTAATGATCAGCCCACTGACGCGTTGCGCGAGAAG
ATTGTGCACCGCCGCTTTACAGGCTTCGACGCCGCTTCGTTCTACCATCGACACCACCACGCTGGCACCCAGTTGATCGGCG
CGAGATTTAATCGCCGCGACAATTTGCGACGGCGCGTGCAGGGCCAGACTGGAGGTGGCAACGCCAATCAGCAACGACTGTT
TGCCCGCCAGTTGTTGTGCCACGCGGTTGGGAATGTAATTCAGCTCCGCCATCGCCGCTTCCACTTTTTCCCGCGTTTTCGC
AGAAACGTGGCTGGCCTGGTTCACCACGCGGGAAACGGTCTGATAAGAGACACCGGCATACTCTGCGACATCGTATAACGTT
ACTGGTTTCACATTCACCACCCTGAATTGACTCTCTTCCGGGCGCTATCATGCCATACCGCGAAAGGTTTTGCGCCATTCGA
TGGTGTCCGGGATCTCGACGCTCTCCCTTATGCGACTCCTGCATTAGG

**pL-(GGS)$_3$ |** SapI, (GGS)$_3$, BtsI, KanR, ColE1 ori, ccdB

GAATATTGGGTTTAGTCTTGTTTCATAATTGTTGCAATGAAACGCGGTGAAACATTGCCTGAAACGTTAACTGAAACGCATA
TTTGCGGATTAGTTCATGACTTTATCTCTAACAAATTGAAATTAAACATTTAATTTTATTAAGGCAATTGTGGCACACCCCT
TGCTTTGTCTTTATCAACGCAAATAACAAGTTGATAACAAGCTAGCAGGAGGAATTCCATATGGGCTCTTCAGGGGGAGGGT
CAGGCGGTTCAGGGGGATCAGGGCACTGCTGAACTAGTCTGATACAGTCGACCTGCAGGCATGCAAGCTTGGCTGTTTTGGC
GGATGAGAGAAGATTTTCAGCCTGATACAGATTAAATCAGAACGCAGAAGCGGTCTGATAAAACAGAATTTGCCTGGCGGCA
GTAGCGCGGTGGTCCCACCTGACCCCATGCCGAACTCAGAAGTGAAACGCCGTAGCGCCGATGGTAGTGTGGCCAGAGCCCA
TGCGAGAGTAGGGAACTGCCAGGCATCAAATAAAACGAAAGGCTCAGTCGAAAGACTGGGCCTTTCGTTTTATCTGTTGTTT
GTCGGTGAACGCTCTCCTGAGTAGGACAAATCCGCCGGGAGCGGATTTGAACGTTGCGAAGCAACGGCCCGGAGGGTGGCGG
GCAGGACGCCCGCCATAAACTGCCAGGCATCAAATTAAGCAGAAGGCCATCCTGACGGATGGCCTTTTTGCGTTTCTACAAA
CTCTCTGATCCTTCAACTCAGCAAAAGTTCGATTTATTCAACAAAGCCACGTTGTGTCTCAAAATCTCTGATGTTACATTGC
ACAAGATAAAAATATATCATCATGAACAATAAAACTGTCTGCTTACATAAACAGTAATACAAGGGGTGTTATGAGCCATATT
CAACGGGAAACGTCTTGCTCTAGGCCGCGATTAAATTCCAACATGGATGCTGATTTATATGGGTATAAATGGGCTCGCGATA
ATGTCGGGCAATCAGGTGCGACAATCTATCGATTGTATGGGAAGCCCGATGCGCCAGAGTTGTTTCTGAAACATGGCAAAGG
TAGCGTTGCCAATGATGTTACAGATGAGATGGTCAGACTAAACTGGCTGACGGAATTTATGCCTCTGCCGACCATCAAGCAT
TTTATCCGTACTCCTGATGATGCATGGTTACTCACCACGGCGATCCCAGGGAAAACAGCATTCCAGGTATTAGAAGAATATC
CTGATTCAGGTGAAAATATTGTTGATGCGCTGGCGGTGTTCCTGCGCCGGTTGCATTCGATTCCTGTTTGTAATTGTCCTTT
TAACAGCGACCGCGTATTTCGTCTGGCTCAGGCGCAATCACGAATGAATAACGGTTTGGTTGATGCGAGTGATTTTGATGAC
GAGCGTAATGGCTGGCCTGTTGAACAAGTCTGGAAAGAAATGCATAAACTTTTGCCATTCTCACCGGATTCAGTCGTCACTC
ATGGTGATTTCTCACTTGATAACCTTATTTTTGACGAGGGGAAATTAATAGGTTGTATTGATGTTGGACGAGTCGGAATCGC
AGACCGATACCAGGATCTTGCCATCCTATGGAACTGCCTCGGTGAGTTTTCTCCTTCATTACAGAAACGGCTTTTTCAAAAA
TATGGTATTGATAATCCTGATATGAATAAATTGCAGTTTCATTTGATGCTCGATGAGTTTTTCTAACTGTCAGACCAAGTTT
ACTCATATATACTTTAGATTGATTT**GAAGAC**TACGCGCCCTGTAGCGGCGCATTAAGCGCGGCGGGTGTGGTGGTTACGCGC
AGCGTGACCGCTACACTTGCCAGCGCCCTAGCGCCCGCTCCTTTCGCTTTCTTCCCTTCCTTTCTCGCCACGTTCGCCGGCT
TTCCCCGTCAAGCTCTAAATCGGGGGCTCCCTTTAGGGTTCCGATTTAGTGCTTTACGGCACCTCGACCCCAAAAAACTTGA
TTTGGGTGATGGTTCACGTAGTGGGCCATCGCCCTGATAGACGGTTTTTCGCCCTTTGACGTTGGAGTCCACGTTCTTTAAT
AGTGGACTCTTGTTCCAAACTGGAACAACACTCAACCCTATCTCGGGCTATTCTTTTGATTTATAAGGGATTTTGCCGATTT
CGGCCTATTGGTTAAAAAATGAGCTGATTTAACAAAAATTTAACGCGAATTTTAACAAAATATTAACGTTTACAATTTAAAA
GGATCTAGGTGAAGATCCTTTTTGATAATCTCATGACCAAAATCCCTTAACGTGAGTTTTCGTTCCACTGAGCGTCAGACCC
CGTAGAAAAGATCAAAGGATCTTCTTGAGATCCTTTTTTTCTGCGCGTAATCTGCTGCTTGCAAACAAAAAAACCACCGCTA
CCAGCGGTGGTTTGTTTGCCGGATCAAGAGCTACCAACTCTTTTTCCGAAGGTAACTGGCTTCAGCAGAGCGCAGATACCAA
ATACTGTCCTTCTAGTGTAGCCGTAGTTAGGCCACCACTTCAAGAACTCTGTAGCACCGCCTACATACCTCGCTCTGCTAAT
CCTGTTACCAGTGGCTGCTGCCAGTGGCGATAAGTCGTGTCTTACCGGGTTGGACTCAAGACGATAGTTACCGGATAAGGCG
CAGCGGTCGGGCTGAACGGGGGGTTCGTGCACACAGCCCAGCTTGGAGCGAACGACCTACACCGAACTGAGATACCTACAGC
GTGAGCTATGAGAAAGCGCCACGCTTCCCGAAGGGAGAAAGGCGGACAGGTATCCGGTAAGCGGCAGGGTCGGAACAGGAGA
GCGCACGAGGGAGCTTCCAGGGGGAAACGCCTGGTATCTTTATAGTCCTGTCGGGTTTCGCCACCTCTGACTTGAGCGTCGA
TTTTTGTGATGCTCGTCAGGGGGGCGGAGCCTATGGAAAAACGCCAGCAACGCGGCCTTTTTACGGTTCCTGGCCTTTTGCT
GGCCTTTTGCTCACATGTTCTTTCCTGCGTTATCCCCTGATTCTGTGGATAACCGTATTACCGCCTTTGAGTGAGCTGATAC
CGCTCGCCGCAGCCGAACGACCGAGCGCAGCGAGTCAGTGAGCGAGGAAGCGGTAGAGCGCCTGATGCGGTATTTTCTCCTT
ACGCATCTGTGCGGTATTTCACACCGCATAGGGTCATGGCTGCGCCCCGACACCCGCCAACACCCGCTGACGCGCCCTGACG
GGCTTGTCTGCTCCCGGCATCCGCTTACAGACAAGCTGTGACCGTGTCCGGGAGCTGCATGTGTCAGAGGTTTTCACCGTCA
TCACCGAAACGCGCGAGGCAGAAGGAGATGGCGCCCAACAGTCCCCCGGCCACGGGGCCTGCCACCATACCCACGCCGAAAC
AAGCGCTCATGAGCCCGAAGTGGCGAGCCCGATCTTCCCCATCGGTGATGTCGGCGATATAGGCGCCAGCAACCGCACCTGT
GGCGCCGGTGATGCCGGCCACGATGCGTCCGGCGTAGAGGATCTGCTCATGTTTGACAGCTTATCATCGATTTATATTCCCC
AGAACATCAGGTTAATGGCGTTTTTGATGTCATTTTCGCGGTGGCTGAGATCAGCCACTTCTTCCCCGATAACGGAGACTGG
CACACTGGCCATATCGGTGGTCATCATGCGCCAGCTTTCATCCCCGATATGCACCACCGGGTAAAGTTCACGGGAGACTTTA
TCTGACAGCAGACGTGCACTGGCCAGGGGGATCACCATCCGTCGCCCGGGCGTGTCAATAATATCACTCTGTACATCCACAA
ACAGACGATAACGGCTCTCTCTTTTATAGGTGTAAACCTTAAACTGCATAGCGCACCGCAAAGTTAAGAAACC

**pL-$(TP)_4T$ | SapI, $(TP)_4T$, BtsI, KanR, BbsI, ColE1 ori, ccdB**

GAATATTGGGTTTAGTCTTGTTTCATAATTGTTGCAATGAAACGCGGTGAAACATTGCCTGAAACGTTAACTGAAACGCATA
TTTGCGGATTAGTTCATGACTTTATCTCTAACAAATTGAAATTAAACATTTAATTTTATTAAGGCAATTGTGGCACACCCCT
TGCTTTGTCTTTATCAACGCAAATAACAAGTTGATAACAAGCTAGCAGGAGGAATTCCATATGGGCTCTTCAGGGACTCCCA
CTCCTACACCTACTCCAACCGGGCACTGCTGAACTAGTCTGATACAGTCGACCTGCAGGCATGCAAGCTTGGCTGTTTTGGC
GGATGAGAGAAGATTTTCAGCCTGATACAGATTAAATCAGAACGCAGAAGCGGTCTGATAAAACAGAATTTGCCTGGCGGCA
GTAGCGCGGTGGTCCCACCTGACCCCATGCCGAACTCAGAAGTGAAACGCCGTAGCGCCGATGGTAGTGTGGCCAGAGCCCA
TGCGAGAGTAGGGAACTGCCAGGCATCAAATAAAACGAAAGGCTCAGTCGAAAGACTGGGCCTTTCGTTTTATCTGTTGTTT
GTCGGTGAACGCTCTCCTGAGTAGGACAAATCCGCCGGGAGCGGATTTGAACGTTGCGAAGCAACGGCCCGGAGGGTGGCGG
GCAGGACGCCCGCCATAAACTGCCAGGCATCAAATTAAGCAGAAGGCCATCCTGACGGATGGCCTTTTTGCGTTTCTACAAA
CTCTCTGATCCTTCAACTCAGCAAAAGTTCGATTTATTCAACAAAGCCACGTTGTGTCTCAAAATCTCTGATGTTACATTGC
ACAAGATAAAAATATATCATCATGAACAATAAAACTGTCTGCTTACATAAACAGTAATACAAGGGGTGTTATGAGCCATATT
CAACGGGAAACGTCTTGCTCTAGGCCGCGATTAAATTCCAACATGGATGCTGATTTATATGGGTATAAATGGGCTCGCGATA
ATGTCGGGCAATCAGGTGCGACAATCTATCGATTGTATGGGAAGCCCGATGCGCCAGAGTTGTTTCTGAAACATGGCAAAGG
TAGCGTTGCCAATGATGTTACAGATGAGATGGTCAGACTAAACTGGCTGACGGAATTTATGCCTCTGCCGACCATCAAGCAT
TTTATCCGTACTCCTGATGATGCATGGTTACTCACCACGGCGATCCCAGGGAAAACAGCATTCCAGGTATTAGAAGAATATC
CTGATTCAGGTGAAAATATTGTTGATGCGCTGGCGGTGTTCCTGCGCCGGTTGCATTCGATTCCTGTTTGTAATTGTCCTTT
TAACAGCGACCGCGTATTTCGTCTGGCTCAGGCGCAATCACGAATGAATAACGGTTTGGTTGATGCGAGTGATTTTGATGAC
GAGCGTAATGGCTGGCCTGTTGAACAAGTCTGGAAAGAAATGCATAAACTTTTGCCATTCTCACCGGATTCAGTCGTCACTC
ATGGTGATTTCTCACTTGATAACCTTATTTTTGACGAGGGGAAATTAATAGGTTGTATTGATGTTGGACGAGTCGGAATCGC
AGACCGATACCAGGATCTTGCCATCCTATGGAACTGCCTCGGTGAGTTTTCTCCTTCATTACAGAAACGGCTTTTTCAAAAA
TATGGTATTGATAATCCTGATATGAATAAATTGCAGTTTCATTTGATGCTCGATGAGTTTTTCTAACTGTCAGACCAAGTTT
ACTCATATATACTTTAGATTGATTT**GAAGAC**TACGCGCCCTGTAGCGGCGCATTAAGCGCGGCGGGTGTGGTGGTTACGCGC
AGCGTGACCGCTACACTTGCCAGCGCCCTAGCGCCCGCTCCTTTCGCTTTCTTCCCTTCCTTTCTCGCCACGTTCGCCGGCT
TTCCCCGTCAAGCTCTAAATCGGGGGCTCCCTTTAGGGTTCCGATTTAGTGCTTTACGGCACCTCGACCCCAAAAAACTTGA
TTTGGGTGATGGTTCACGTAGTGGGCCATCGCCCTGATAGACGGTTTTTCGCCCTTTGACGTTGGAGTCCACGTTCTTTAAT
AGTGGACTCTTGTTCCAAACTGGAACAACACTCAACCCTATCTCGGGCTATTCTTTTGATTTATAAGGGATTTTGCCGATTT
CGGCCTATTGGTTAAAAAATGAGCTGATTTAACAAAAATTTAACGCGAATTTTAACAAAATATTAACGTTTACAATTTAAAA
GGATCTAGGTGAAGATCCTTTTTGATAATCTCATGACCAAAATCCCTTAACGTGAGTTTTCGTTCCACTGAGCGTCAGACCC
CGTAGAAAAGATCAAAGGATCTTCTTGAGATCCTTTTTTTCTGCGCGTAATCTGCTGCTTGCAAACAAAAAAACCACCGCTA
CCAGCGGTGGTTTGTTTGCCGGATCAAGAGCTACCAACTCTTTTTCCGAAGGTAACTGGCTTCAGCAGAGCGCAGATACCAA
ATACTGTCCTTCTAGTGTAGCCGTAGTTAGGCCACCACTTCAAGAACTCTGTAGCACCGCCTACATACCTCGCTCTGCTAAT
CCTGTTACCAGTGGCTGCTGCCAGTGGCGATAAGTCGTGTCTTACCGGGTTGGACTCAAGACGATAGTTACCGGATAAGGCG
CAGCGGTCGGGCTGAACGGGGGGTTCGTGCACACAGCCCAGCTTGGAGCGAACGACCTACACCGAACTGAGATACCTACAGC
GTGAGCTATGAGAAAGCGCCACGCTTCCCGAAGGGAGAAAGGCGGACAGGTATCCGGTAAGCGGCAGGGTCGGAACAGGAGA
GCGCACGAGGGAGCTTCCAGGGGGAAACGCCTGGTATCTTTATAGTCCTGTCGGGTTTCGCCACCTCTGACTTGAGCGTCGA
TTTTTGTGATGCTCGTCAGGGGGGCGGAGCCTATGGAAAAACGCCAGCAACGCGGCCTTTTTACGGTTCCTGGCCTTTTGCT
GGCCTTTTGCTCACATGTTCTTTCCTGCGTTATCCCCTGATTCTGTGGATAACCGTATTACCGCCTTTGAGTGAGCTGATAC
CGCTCGCCGCAGCCGAACGACCGAGCGCAGCGAGTCAGTGAGCGAGGAAGCGGTAGAGCGCCTGATGCGGTATTTTCTCCTT
ACGCATCTGTGCGGTATTTCACACCGCATAGGGTCATGGCTGCGCCCCGACACCCGCCAACACCCGCTGACGCGCCCTGACG
GGCTTGTCTGCTCCCGGCATCCGCTTACAGACAAGCTGTGACCGTGTCCGGGAGCTGCATGTGTCAGAGGTTTTCACCGTCA
TCACCGAAACGCGCGAGGCAGAAGGAGATGGCGCCCAACAGTCCCCCGGCCACGGGGCCTGCCACCATACCCACGCCGAAAC
AAGCGCTCATGAGCCCGAAGTGGCGAGCCCGATCTTCCCCATCGGTGATGTCGGCGATATAGGCGCCAGCAACCGCACCTGT
GGCGCCGGTGATGCCGGCCACGATGCGTCCGGCGTAGAGGATCTGCTCATGTTTGACAGCTTATCATCGATTTATATTCCCC
AGAACATCAGGTTAATGGCGTTTTTGATGTCATTTTCGCGGTGGCTGAGATCAGCCACTTCTTCCCCGATAACGGAGACTGG
CACACTGGCCATATCGGTGGTCATCATGCGCCAGCTTTCATCCCCGATATGCACCACCGGGTAAAGTTCACGGAGACTTTA
TCTGACAGCAGACGTGCACTGGCCAGGGGGATCACCATCCGTCGCCCGGGCGTGTCAATAATATCACTCTGTACATCCACAA
ACAGACGATAACGGCTCTCTCTTTTATAGGTGTAAACCTTAAACTGCATAGCGCACCGCAAAGTTAAGAAACC

**pD-mCherry |** BtsI, mCherry, SapI, AmpR, **BsaI**, ColE1 ori, ccdB

GAATATTGGGTTTAGTCTTGTTTCATAATTGTTGCAATGAAACGCGGTGAAACATTGCCTGAAACGTTAACTGAAACGCATA
TTTGCGGATTAGTTCATGACTTTATCTCTAACAAATTGAAATTAAACATTTAATTTTATTAAGGCAATTGTGGCACACCCCT
TGCTTTGTCTTTATCAACGCAAATAACAAGTTGATAACAAGCTAGCAGGAGGAATTCCATATGGGCAGTGGGGTGAGCAAGG
GCGAGGAGGATAACCTGGCCATCATCAAGGAGTTCATGCGCTTCAAGGTTCACATGGAGGGCTCCGTGAACGGCCACGAGTT
CGAGATCGAGGGCGAGGGCGAGGGCCGCCCCTACGAGGGCACCCAGACCGCCAAGCTGAAGGTGACCAAGGGTGGCCCCCTG
CCCTTCGCCTGGGACATCCTGTCCCCTCAGTTCATGTACGGCTCCAAGGCCTACGTGAAGCACCCCGCCGACATCCCCGACT
ACTTGAAGCTGTCCTTCCCCGAGGGCTTCAAGTGGGAGCGCGTGATGAACTTCGAGGACGGCGGCGTGGTGACCGTGACCCA
GGACTCCTCCCTGCAAGACGGCGAGTTCATCTACAAGGTGAAGCTGCGCGGCACCAACTTCCCCTCCGACGGCCCCGTAATG
CAGAAAAAGACCATGGGCTGGGAGGCCTCCTCCGAGCGGATGTACCCCGAGGACGGCGCGCTGAAGGGCGAGATCAAGCAGA
GGCTGAAGCTGAAGGACGGCGGCCACTACGACGCTGAGGTCAAGACCACCTACAAGGCCAAGAAGCCCGTGCAACTGCCCGG
CGCGTACAACGTCAACATCAAGTTGGACATCACCTCCCACAACGAGGACTACACCATCGTGGAACAGTACGAACGCGCCGAG
GGCCGCCACTCCACCGGCGGCATGGACGAGCTGTACAAGGGGTGAAGAGCGTCGACCTGCAGGCATGCAAGCTTGGCTGTTT
TGGCGGATGAGAGAAGATTTTCAGCCTGATACAGATTAAATCAGAACGCAGAAGCGGTCTGATAAAACAGAATTTGCCTGGC
GGCAGTAGCGCGGTGGTCCCACCTGACCCCATGCCGAACTCAGAAGTGAAACGCCGTAGCGCCATGGTAGTGTGGCCAGAGC
CCATGCGAGAGTAGGGAACTGCCAGGCATCAAATAAAACGAAAGGCTCAGTCGAAAGACTGGGCCTTTCGTTTTATCTGTTG
TTTGTCGGTGAACGCTCTCCTGAGTAGGACAAATCCGCCGGGAGCGGATTTGAACGTTGCGAAGCAACGGCCCGGAGGGTGG
CGGGCAGGACGCCCGCCATAAACTGCCAGGCATCAAATTAAGCAGAAGGCCATCCTGACGGATGGCCTTTTTGCGTTTCTAC
AAACTCTTTTGTTTATTTTTCTAAATACATTCAAATATGTATCCGCTCATGAGACAATAACCCTGATAAATGCTTCAATAAT
ATTGAAAAAGGAAGTGTATGAGTATTCAACATTTCCGTGTCGCCCTTATTCCCTTTTTTGCGGCATTTTGCCTTCCTGTTTT
TGCTCACCCAGAAACGCTGGTGAAAGTAAAAGATGCTGAAGATCAGTTGGGTGCACGAGTGGGTTACATCGAACTGGATCTC
AACAGCGGTAAGATCCTTGAGAGTTTTCGCCCCGAAGAACGTTTTCCAATGATGAGCACTTTTAAAGTTCTGCTATGTGGCG
CGGTATTATCCCGTATTGACGCCGGGCAAGAGCAACTCGGTCGCCGCATACACTATTCTCAGAATGACTTGGTTGAGTACTC
ACCAGTCACAGAAAAGCATCTTACGGATGGCATGACAGTAAGAGAATTATGTAGTGCTGCCATAACCATGAGTGATAACACA
GCGGCCAACTTACTTCTGACAACGATCGGAGGACCGAAGGAGCTAACCGCTTTTTTGCACAACATGGGGGATCATGTAACTC
GCCTTGATCGTTGGGAACCGGAGCTGAATGAAGCCATACCAAACGACGAGCGTGACACCACGATGCCTGTAGCGATGGCAAC
AACGTTGCGCAAACTATTAACTGGCGAACTACTTACTCTAGCTTCCCGGCAACAATTAATAGACTGGATGGAGGCGGATAAA
GTTGCAGGACCACTTCTTCGCTCAGCACTTCCAGCTGGTTGGTTTATTGCTGATAAATCTGGAGCCGGTGAGCGTGGCTCTC
GCGGTATCATTGCAGCACTGGGGCCAGATGGTAAGCCCTCCCGTATCGTAGTTATCTACACGACGGGGAGTCAGGCAACTAT
GGATGAACGAAATAGACAGATCGCTGAGATAGGTGCCTCACTGATTAAGCATTGGTAACTGTCAGACCAAGTTTACTCATAT
ATACTTTAGATTGATTT**GGTCTC**ACGCGCCCTGTAGCGGCGCATTAAGCGCGGCGGGTGTGGTGGTTACGCGCAGCGTGACC
GCTACACTTGCCAGCGCCCTAGCGCCCGCTCCTTTCGCTTTCTTCCCTTCCTTTCTCGCCACGTTCGCCGGCTTTCCCCGTC
AAGCTCTAAATCGGGGGCTCCCTTTAGGGTTCCGATTTAGTGCTTTACGGCACCTCGACCCCAAAAAACTTGATTTGGGTGA
TGGTTCACGTAGTGGGCCATCGCCCTGATAGACGGTTTTTCGCCCTTTGACGTTGGAGTCCACGTTCTTTAATAGTGGACTC
TTGTTCCAAACTGGAACAACACTCAACCCTATCTCGGGCTATTCTTTTGATTTATAAGGGATTTTGCCGATTTCGGCCTATT
GGTTAAAAAATGAGCTGATTTAACAAAAATTTAACGCGAATTTTAACAAAATATTAACGTTTACAATTTAAAAGGATCTAGG
TGAAGATCCTTTTTGATAATCTCATGACCAAAATCCCTTAACGTGAGTTTTCGTTCCACTGAGCGTCAGACCCCGTAGAAAA
GATCAAAGGATCTTCTTGAGATCCTTTTTTTCTGCGCGTAATCTGCTGCTTGCAAACAAAAAAACCACCGCTACCAGCGGTG
GTTTGTTTGCCGGATCAAGAGCTACCAACTCTTTTTCCGAAGGTAACTGGCTTCAGCAGAGCGCAGATACCAAATACTGTCC
TTCTAGTGTAGCCGTAGTTAGGCCACCACTTCAAGAACTCTGTAGCACCGCCTACATACCTCGCTCTGCTAATCCTGTTACC
AGTGGCTGCTGCCAGTGGCGATAAGTCGTGTCTTACCGGGTTGGACTCAAGACGATAGTTACCGGATAAGGCGCAGCGGTCG
GGCTGAACGGGGGGTTCGTGCACACAGCCCAGCTTGGAGCGAACGACCTACACCGAACTGAGATACCTACAGCGTGAGCTAT
GAGAAAGCGCCACGCTTCCCGAAGGGAGAAAGGCGGACAGGTATCCGGTAAGCGGCAGGGTCGGAACAGGAGAGCGCACGAG
GGAGCTTCCAGGGGGAAACGCCTGGTATCTTTATAGTCCTGTCGGGTTTCGCCACCTCTGACTTGAGCGTCGATTTTTGTGA
TGCTCGTCAGGGGGGCGGAGCCTATGGAAAAACGCCAGCAACGCGGCCTTTTTACGGTTCCTGGCCTTTTGCTGGCCTTTTG
CTCACATGTTCTTTCCTGCGTTATCCCCTGATTCTGTGGATAACCGTATTACCGCCTTTGAGTGAGCTGATACCGCTCGCCG
CAGCCGAACGACCGAGCGCAGCGAGTCAGTGAGCGAGGAAGCGGTAGAGCGCCTGATGCGGTATTTTCTCCTTACGCATCTG
TGCGGTATTTCACACCGCATAGGGTCATGGCTGCGCCCCGACACCCGCCAACACCCGCTGACGCGCCCTGACGGGCTTGTCT
GCTCCCGGCATCCGCTTACAGACAAGCTGTGACCGTGTCCGGGAGCTGCATGTGTCAGAGGTTTTCACCGTCATCACCGAAA
CGCGCGAGGCAGAAGGAGATGGCGCCCAACAGTCCCCCGGCCACGGGGCCTGCCACCATACCCACGCCGAAACAAGCGCTCA
TGAGCCCGAAGTGGCGAGCCCGATCTTCCCCATCGGTGATGTCGGCGATATAGGCGCCAGCAACCGCACCTGTGGCGCCGGT
GATGCCGGCCACGATGCGTCCGGCGTAGAGGATCTGCTCATGTTTGACAGCTTATCATCGATTTATATTCCCCAGAACATCA
GGTTAATGGCGTTTTTGATGTCATTTTCGCGGTGGCTGAGATCAGCCACTTCTTCCCCGATAACGGAGACTGGCACACTGGC
CATATCGGTGGTCATCATGCGCCAGCTTTCATCCCCGATATGCACCACCGGGTAAAGTTCACGGGAGACTTTATCTGACAGC
AGACGTGCACTGGCCAGGGGGATCACCATCCGTCGCCCGGGCGTGTCAATAATATCACTCTGTACATCCACAAACAGACGAT
AACGGCTCTCTCTTTTATAGGTGTAAACCTTAAACTGCATAGCGCACCGCAAAGTTAAGAAACC

**pD-CyOFP1 | BtsI, CyOFP1, SapI, AmpR, BsaI, ColE1 ori, ccdB**

GAATATTGGGTTTAGTCTTGTTTCATAATTGTTGCAATGAAACGCGGTGAAACATTGCCTGAAACGTTAACTGAAACGCATA
TTTGCGGATTAGTTCATGACTTTATCTCTAACAAATTGAAATTAAACATTTAATTTTATTAAGGCAATTGTGGCACACCCCT
TGCTTTGTCTTTATCAACGCAAATAACAAGTTGATAACAAGCTAGCAGGAGGAATTCCATATGGGCAGTGGGGTCAGTAAGG
GCGAGGAACTGATTAAGGAGAATATGCGCTCTAAGTTATATCTGGAAGGGTCGGTTAATGGCCATCAGTTCAAATGTACCCA
CGAAGGCGAAGGAAAACCTTACGAAGGGAAACAGACAAATCGCATTAAAGTTGTGGAGGGGGGTCCTTTACCATTTGCGTTC
GATATTCTTGCTACTCACTTTATGTATGGAAGCAAAGTGTTCATTAAATACCCCGCCGATCTTCCAGACTATTTCAAGCAGT
CATTCCCCGAGGGTTTTACATGGGAACGTGTGATGGTGTTCGAGGATGGAGGCGTCCTTACCGCGACACAAGACACATCCCT
CCAGGATGGAGAGCTTATTTACAATGTTAAGGTCCGCGGCGTGAACTTCCCAGCTAACGGTCCGGTTATGCAGAAAAAGACG
TTAGGCTGGGAGCCTTCGACAGAAACTATGTATCCGGCTGACGGCGGGCTTGAGGGTCGTTGCGACAAAGCGCTTAAGTTAG
TAGGCGGAGGTCACCTTCATGTGAATTTTAAGACCACATATAAAAGTAAAAAACCCGTTAAGATGCCTGGCGTCCACTATGT
GGATCGTCGCCTGGAGCGTATTAAGGAAGCTGATAACGAGACTTATGTAGAGCAGTACGAACATGCCGTGGCACGTTATTCC
AATTTGGGAGGGGGGATGGACGAGCTTTATAAGGGGTGAAGAGCGTCGACCTGCAGGCATGCAAGCTTGGCTGTTTTGGCGG
ATGAGAGAAGATTTTCAGCCTGATACAGATTAAATCAGAACGCAGAAGCGGTCTGATAAAACAGAATTTGCCTGGCGGCAGT
AGCGCGGTGGTCCCACCTGACCCCATGCCGAACTCAGAAGTGAAACGCCGTAGCGCCATGGTAGTGTGGCCAGAGCCCATGC
GAGAGTAGGGAACTGCCAGGCATCAAATAAAACGAAAGGCTCAGTCGAAAGACTGGGCCTTTCGTTTTATCTGTTGTTTGTC
GGTGAACGCTCTCCTGAGTAGGACAAATCCGCCGGGAGCGGATTTGAACGTTGCGAAGCAACGGCCCGGAGGGTGGCGGGCA
GGACGCCCGCCATAAACTGCCAGGCATCAAATTAAGCAGAAGGCCATCCTGACGGATGGCCTTTTTGCGTTTCTACAAACTC
TTTTGTTTATTTTTCTAAATACATTCAAATATGTATCCGCTCATGAGACAATAACCCTGATAAATGCTTCAATAATATTGAA
AAAGGAAGTGTATGAGTATTCAACATTTCCGTGTCGCCCTTATTCCCTTTTTTGCGGCATTTTGCCTTCCTGTTTTTGCTCA
CCCAGAAACGCTGGTGAAAGTAAAAGATGCTGAAGATCAGTTGGGTGCACGAGTGGGTTACATCGAACTGGATCTCAACAGC
GGTAAGATCCTTGAGAGTTTTCGCCCCGAAGAACGTTTTCCAATGATGAGCACTTTTAAAGTTCTGCTATGTGGCGCGGTAT
TATCCCGTATTGACGCCGGGCAAGAGCAACTCGGTCGCCGCATACACTATTCTCAGAATGACTTGGTTGAGTACTCACCAGT
CACAGAAAAGCATCTTACGGATGGCATGACAGTAAGAGAATTATGTAGTGCTGCCATAACCATGAGTGATAACACAGCGGCC
AACTTACTTCTGACAACGATCGGAGGACCGAAGGAGCTAACCGCTTTTTTGCACAACATGGGGGATCATGTAACTCGCCTTG
ATCGTTGGGAACCGGAGCTGAATGAAGCCATACCAAACGACGAGCGTGACACCACGATGCCTGTAGCGATGGCAACAACGTT
GCGCAAACTATTAACTGGCGAACTACTTACTCTAGCTTCCCGGCAACAATTAATAGACTGGATGGAGGCGGATAAAGTTGCA
GGACCACTTCTTCGCTCAGCACTTCCAGCTGGTTGGTTTATTGCTGATAAATCTGGAGCCGGTGAGCGTGGCTCTCGCGGTA
TCATTGCAGCACTGGGGCCAGATGGTAAGCCCTCCCGTATCGTAGTTATCTACACGACGGGGAGTCAGGCAACTATGGATGA
ACGAAATAGACAGATCGCTGAGATAGGTGCCTCACTGATTAAGCATTGGTAACTGTCAGACCAAGTTTACTCATATATACTT
TAGATTGATTT**GGTCTC**ACGCGCCCTGTAGCGGCGCATTAAGCGCGGCGGGTGTGGTGGTTACGCGCAGCGTGACCGCTACA
CTTGCCAGCGCCCTAGCGCCCGCTCCTTTCGCTTTCTTCCCTTCCTTTCTCGCCACGTTCGCCGGCTTTCCCCGTCAAGCTC
TAAATCGGGGGCTCCCTTTAGGGTTCCGATTTAGTGCTTTACGGCACCTCGACCCCAAAAAACTTGATTTGGGTGATGGTTC
ACGTAGTGGGCCATCGCCCTGATAGACGGTTTTTCGCCCTTTGACGTTGGAGTCCACGTTCTTTAATAGTGGACTCTTGTTC
CAAACTGGAACAACACTCAACCCTATCTCGGGCTATTCTTTTGATTTATAAGGGATTTTGCCGATTTCGGCCTATTGGTTAA
AAAATGAGCTGATTTAACAAAAATTTAACGCGAATTTTAACAAAATATTAACGTTTACAATTTAAAAGGATCTAGGTGAAGA
TCCTTTTTGATAATCTCATGACCAAAATCCCTTAACGTGAGTTTTCGTTCCACTGAGCGTCAGACCCCGTAGAAAAGATCAA
AGGATCTTCTTGAGATCCTTTTTTTCTGCGCGTAATCTGCTGCTTGCAAACAAAAAAACCACCGCTACCAGCGGTGGTTTGT
TTGCCGGATCAAGAGCTACCAACTCTTTTTCCGAAGGTAACTGGCTTCAGCAGAGCGCAGATACCAAATACTGTCCTTCTAG
TGTAGCCGTAGTTAGGCCACCACTTCAAGAACTCTGTAGCACCGCCTACATACCTCGCTCTGCTAATCCTGTTACCAGTGGC
TGCTGCCAGTGGCGATAAGTCGTGTCTTACCGGGTTGGACTCAAGACGATAGTTACCGGATAAGGCGCAGCGGTCGGGCTGA
ACGGGGGGTTCGTGCACACAGCCCAGCTTGGAGCGAACGACCTACACCGAACTGAGATACCTACAGCGTGAGCTATGAGAAA
GCGCCACGCTTCCCGAAGGGAGAAAGGCGGACAGGTATCCGGTAAGCGGCAGGGTCGGAACAGGAGAGCGCACGAGGGAGCT
TCCAGGGGGAAACGCCTGGTATCTTTATAGTCCTGTCGGGTTTCGCCACCTCTGACTTGAGCGTCGATTTTTGTGATGCTCG
TCAGGGGGGCGGAGCCTATGGAAAAACGCCAGCAACGCGGCCTTTTTACGGTTCCTGGCCTTTTGCTGGCCTTTTGCTCACA
TGTTCTTTCCTGCGTTATCCCCTGATTCTGTGGATAACCGTATTACCGCCTTTGAGTGAGCTGATACCGCTCGCCGCAGCCG
AACGACCGAGCGCAGCGAGTCAGTGAGCGAGGAAGCGGTAGAGCGCCTGATGCGGTATTTTCTCCTTACGCATCTGTGCGGT
ATTTCACACCGCATAGGGTCATGGCTGCGCCCCGACACCCGCCAACACCCGCTGACGCGCCCTGACGGGCTTGTCTGCTCCC
GGCATCCGCTTACAGACAAGCTGTGACCGTGTCCGGGAGCTGCATGTGTCAGAGGTTTTCACCGTCATCACCGAAACGCGCG
AGGCAGAAGGAGATGGCGCCCAACAGTCCCCCGGCCACGGGGCCTGCCACCATACCCACGCCGAAACAAGCGCTCATGAGCC
CGAAGTGGCGAGCCCGATCTTCCCCATCGGTGATGTCGGCGATATAGGCGCCAGCAACCGCACCTGTGGCGCCGGTGATGCC
GGCCACGATGCGTCCGGCGTAGAGGATCTGCTCATGTTTGACAGCTTATCATCGATTTATATTCCCCAGAACATCAGGTTAA
TGGCGTTTTTGATGTCATTTTCGCGGTGGCTGAGATCAGCCACTTCTTCCCCGATAACGGAGACTGGCACACTGGCCATATC
GGTGGTCATCATGCGCCAGCTTTCATCCCCGATATGCACCACCGGGTAAAGTTCACGGGAGACTTTATCTGACAGCAGACGT
GCACTGGCCAGGGGGATCACCATCCGTCGCCCGGGCGTGTCAATAATATCACTCTGTACATCCACAAACAGACGATAACGGC
TCTCTCTTTTATAGGTGTAAACCTTAAACTGCATAGCGCACCGCAAAGTTAAGAAACC

**pD-sfGFP | BtsI, sfGFP, SapI, AmpR, BsaI, ColE1 ori, ccdB**

GAATATTGGGTTTAGTCTTGTTTCATAATTGTTGCAATGAAACGCGGTGAAACATTGCCTGAAACGTTAACTGAAACGCATA
TTTGCGGATTAGTTCATGACTTTATCTCTAACAAATTGAAATTAAACATTTAATTTTATTAAGGCAATTGTGGCACACCCCT
TGCTTTGTCTTTATCAACGCAAATAACAAGTTGATAACAAGCTAGCAGGAGGAATTCCATATGGGCAGTGGGAGCAAAGGAG
AAGAACTTTTCACTGGGGTAGTCCCCATTTTGGTTGAATTAGACGGGGACGTGAATGGTCACAAATTTAGTGTTCGCGGGGA
GGGGGAGGGGGATGCCACCAATGGTAAGTTGACTTTAAAGTTCATCTGCACTACTGGGAAATTGCCAGTTCCGTGGCCAACT
CTTGTAACCACTTTAACCTATGGAGTTCAGTGCTTCAGCCGTTATCCGGATCACATGAAGCGCCACGACTTCTTCAAGAGTG
CGATGCCGGAGGGATACGTTCAAGAACGTACAATTAGCTTCAAGGATGATGGTACTTACAAAACCCGTGCCGAGGTTAAGTT
TGAAGGCGATACCTTGGTCAACCGCATTGAATTGAAAGGGATTGATTTTAAAGAAGATGGCAACATCTTGGGACACAAGCTG
GAGTACAATTTTAACAGTCACAACGTGTATATCACcGCCGACAAGCAAAAGAATGGTATCAAGGCGAACTTTAAAATTCGTC
ATAATGTTGAGGATGGGTCAGTGCAGCTGGCTGATCACTACCAACAAAACACGCCAATCGGAGATGGACCAGTATTACTTCC
GGACAATCACTACTTATCTACACAATCTGTCTTGTCCAAGGACCCAAATGAAAAACGTGATCACATGGTGTTGCTTGAATTT
GTCACCGCAGCAGGGATCACGCATGGAATGGACGAGCTTTATAAAGGGTGAAGAGCGTCGACCTGCAGGCATGCAAGCTTGG
CTGTTTTGGCGGATGAGAGAAGATTTTCAGCCTGATACAGATTAAATCAGAACGCAGAAGCGGTCTGATAAAACAGAATTTG
CCTGGCGGCAGTAGCGCGGTGGTCCCACCTGACCCCATGCCGAACTCAGAAGTGAAACGCCGTAGCGCCATGGTAGTGTGGC
CAGAGCCCATGCGAGAGTAGGGAACTGCCAGGCATCAAATAAAACGAAAGGCTCAGTCGAAAGACTGGGCCTTTCGTTTTAT
CTGTTGTTTGTCGGTGAACGCTCTCCTGAGTAGGACAAATCCGCCGGGAGCGGATTTGAACGTTGCGAAGCAACGGCCCGGA
GGGTGGCGGGCAGGACGCCCGCCATAAACTGCCAGGCATCAAATTAAGCAGAAGGCCATCCTGACGGATGGCCTTTTTGCGT
TTCTACAAACTCTTTTGTTTATTTTTCTAAATACATTCAAATATGTATCCGCTCATGAGACAATAACCCTGATAAATGCTTC
AATAATATTGAAAAAGGAAGTGTATGAGTATTCAACATTTCCGTGTCGCCCTTATTCCCTTTTTTGCGGCATTTTGCCTTCC
TGTTTTTGCTCACCCAGAAACGCTGGTGAAAGTAAAAGATGCTGAAGATCAGTTGGGTGCACGAGTGGGTTACATCGAACTG
GATCTCAACAGCGGTAAGATCCTTGAGAGTTTTCGCCCCGAAGAACGTTTTCCAATGATGAGCACTTTTAAAGTTCTGCTAT
GTGGCGCGGTATTATCCCGTATTGACGCCGGGCAAGAGCAACTCGGTCGCCGCATACACTATTCTCAGAATGACTTGGTTGA
GTACTCACCAGTCACAGAAAAGCATCTTACGGATGGCATGACAGTAAGAGAATTATGTAGTGCTGCCATAACCATGAGTGAT
AACACAGCGGCCAACTTACTTCTGACAACGATCGGAGGACCGAAGGAGCTAACCGCTTTTTTGCACAACATGGGGGATCATG
TAACTCGCCTTGATCGTTGGGAACCGGAGCTGAATGAAGCCATACCAAACGACGAGCGTGACACCACGATGCCTGTAGCGAT
GGCAACAACGTTGCGCAAACTATTAACTGGCGAACTACTTACTCTAGCTTCCCGGCAACAATTAATAGACTGGATGGAGGCG
GATAAAGTTGCAGGACCACTTCTTCGCTCAGCACTTCCAGCTGGTTGGTTTATTGCTGATAAATCTGGAGCCGGTGAGCGTG
GCTCTCGCGGTATCATTGCAGCACTGGGGCCAGATGGTAAGCCCTCCCGTATCGTAGTTATCTACACGACGGGGAGTCAGGC
AACTATGGATGAACGAAATAGACAGATCGCTGAGATAGGTGCCTCACTGATTAAGCATTGGTAACTGTCAGACCAAGTTTAC
TCATATATACTTTAGATTGATTT**GGTCTC**ACGCGCCCTGTAGCGGCGCATTAAGCGCGGCGGGTGTGGTGGTTACGCGCAGC
GTGACCGCTACACTTGCCAGCGCCCTAGCGCCCGCTCCTTTCGCTTTCTTCCCTTCCTTTCTCGCCACGTTCGCCGGCTTTC
CCCGTCAAGCTCTAAATCGGGGGCTCCCTTTAGGGTTCCGATTTAGTGCTTTACGGCACCTCGACCCCAAAAAACTTGATTT
GGGTGATGGTTCACGTAGTGGGCCATCGCCCTGATAGACGGTTTTTCGCCCTTTGACGTTGGAGTCCACGTTCTTTAATAGT
GGACTCTTGTTCCAAACTGGAACAACACTCAACCCTATCTCGGGCTATTCTTTTGATTTATAAGGGATTTTGCCGATTTCGG
CCTATTGGTTAAAAAATGAGCTGATTTAACAAAAATTTAACGCGAATTTTAACAAAATATTAACGTTTACAATTTAAAAGGA
TCTAGGTGAAGATCCTTTTTGATAATCTCATGACCAAAATCCCTTAACGTGAGTTTTCGTTCCACTGAGCGTCAGACCCCGT
AGAAAAGATCAAAGGATCTTCTTGAGATCCTTTTTTTCTGCGCGTAATCTGCTGCTTGCAAACAAAAAAACCACCGCTACCA
GCGGTGGTTTGTTTGCCGGATCAAGAGCTACCAACTCTTTTTCCGAAGGTAACTGGCTTCAGCAGAGCGCAGATACCAAATA
CTGTCCTTCTAGTGTAGCCGTAGTTAGGCCACCACTTCAAGAACTCTGTAGCACCGCCTACATACCTCGCTCTGCTAATCCT
GTTACCAGTGGCTGCTGCCAGTGGCGATAAGTCGTGTCTTACCGGGTTGGACTCAAGACGATAGTTACCGGATAAGGCGCAG
CGGTCGGGCTGAACGGGGGGTTCGTGCACACAGCCCAGCTTGGAGCGAACGACCTACACCGAACTGAGATACCTACAGCGTG
AGCTATGAGAAAGCGCCACGCTTCCCGAAGGGAGAAAGGCGGACAGGTATCCGGTAAGCGGCAGGGTCGGAACAGGAGAGCG
CACGAGGGAGCTTCCAGGGGGAAACGCCTGGTATCTTTATAGTCCTGTCGGGTTTCGCCACCTCTGACTTGAGCGTCGATTT
TTGTGATGCTCGTCAGGGGGGCGGAGCCTATGGAAAAACGCCAGCAACGCGGCCTTTTTACGGTTCCTGGCCTTTTGCTGGC
CTTTTGCTCACATGTTCTTTCCTGCGTTATCCCCTGATTCTGTGGATAACCGTATTACCGCCTTTGAGTGAGCTGATACCGC
TCGCCGCAGCCGAACGACCGAGCGCAGCGAGTCAGTGAGCGAGGAAGCGGTAGAGCGCCTGATGCGGTATTTTCTCCTTACG
CATCTGTGCGGTATTTCACACCGCATAGGGTCATGGCTGCGCCCCGACACCCGCCAACACCCGCTGACGCGCCCTGACGGGC
TTGTCTGCTCCCGGCATCCGCTTACAGACAAGCTGTGACCGTGTCCGGGAGCTGCATGTGTCAGAGGTTTTCACCGTCATCA
CCGAAACGCGCGAGGCAGAAGGAGATGGCGCCCAACAGTCCCCCGGCCACGGGGCCTGCCACCATACCCACGCCGAAACAAG
CGCTCATGAGCCCGAAGTGGCGAGCCCGATCTTCCCCATCGGTGATGTCGGCGATATAGGCGCCAGCAACCGCACCTGTGGC
GCCGGTGATGCCGGCCACGATGCGTCCGGCGTAGAGGATCTGCTCATGTTTGACAGCTTATCATCGATTTATATTCCCCAGA
ACATCAGGTTAATGGCGTTTTTGATGTCATTTTCGCGGTGGCTGAGATCAGCCACTTCTTCCCCGATAACGGAGACTGGCAC
ACTGGCCATATCGGTGGTCATCATGCGCCAGCTTTCATCCCCGATATGCACCACCGGGTAAAGTTCACGGGAGACTTTATCT
GACAGCAGACGTGCACTGGCCAGGGGGATCACCATCCGTCGCCCGGGCGTGTCAATAATATCACTCTGTACATCCACAAACA
GACGATAACGGCTCTCTCTTTTATAGGTGTAAACCTTAAACTGCATAGCGCACCGCAAAGTTAAGAAACC

**pD-mCerulean | BtsI, mCerulean, SapI, AmpR, BsaI, ColE1 ori, ccdB**

GAATATTGGGTTTAGTCTTGTTTCATAATTGTTGCAATGAAACGCGGTGAAACATTGCCTGAAACGTTAACTGAAACGCATA
TTTGCGGATTAGTTCATGACTTTATCTCTAACAAATTGAAATTAAACATTTAATTTTATTAAGGCAATTGTGGCACACCCCT
TGCTTTGTCTTTATCAACGCAAATAACAAGTTGATAACAAGCTAGCAGGAGGAATTCCATATGGGCAGTGGGGTGAGCAAGG
GCGAGGAGCTGTTCACCGGGGTGGTGCCCATCCTGGTCGAGCTGGACGGCGACGTAAACGGCCACAAGTTCAGCGTGTCCGG
CGAGGGCGAGGGCGATGCCACCTACGGCAAGCTGACCCTGAAGTTCATCTGCACCACCGGCAAGCTGCCCGTGCCCTGGCCC
ACCCTCGTGACCACCCTGAGCTGGGGCGTtCAGTGCTTCGCCCGCTACCCCGACCACATGAAGCAGCACGACTTCTTCAAGT
CCGCCATGCCCGAAGGCTACGTCCAGGAGCGCACCATCTTCTTCAAGGACGACGGCAACTACAAGACCCGCGCCGAGGTGAA
GTTCGAGGGCGACACCCTGGTGAACCGCATCGAGCTGAAGGGCATCGACTTCAAGGAGGACGGCAACATCCTGGGGCACAAG
CTGGAGTACAACGCCATCCACGGCAACGTCTATATCACCGCCGACAAGCAGAAGAACGGCATCAAGGCCAACTTCGGCCTCA
ACTGCAACATCGAGGACGGCAGCGTGCAGCTCGCCGACCACTACCAGCAGAACACCCCCATCGGCGACGGCCCCGTGCTGCT
GCCCGACAACCACTACCTGAGCACCCAGTCCAAGCTGAGCAAAGACCCCAACGAGAAGCGCGATCACATGGTCCTGCTGGAG
TTCGTGGGGTGAAGAGCGTCGACCTGCAGGCATGCAAGCTTGGCTGTTTTGGCGGATGAGAGAAGATTTTCAGCCTGATACA
GATTAAATCAGAACGCAGAAGCGGTCTGATAAAACAGAATTTGCCTGGCGGCAGTAGCGCGGTGGTCCCACCTGACCCCATG
CCGAACTCAGAAGTGAAACGCCGTAGCGCCATGGTAGTGTGGCCAGAGCCCATGCGAGAGTAGGGAACTGCCAGGCATCAAA
TAAAACGAAAGGCTCAGTCGAAAGACTGGGCCTTTCGTTTTATCTGTTGTTTGTCGGTGAACGCTCTCCTGAGTAGGACAAA
TCCGCCGGGAGCGGATTTGAACGTTGCGAAGCAACGGCCCGGAGGGTGGCGGGCAGGACGCCCGCCATAAACTGCCAGGCAT
CAAATTAAGCAGAAGGCCATCCTGACGGATGGCCTTTTTGCGTTTCTACAAACTCTTTTGTTTATTTTTCTAAATACATTCA
AATATGTATCCGCTCATGAGACAATAACCCTGATAAATGCTTCAATAATATTGAAAAAGGAAGTGTATGAGTATTCAACATT
TCCGTGTCGCCCTTATTCCCTTTTTTGCGGCATTTTGCCTTCCTGTTTTTGCTCACCCAGAAACGCTGGTGAAAGTAAAAGA
TGCTGAAGATCAGTTGGGTGCACGAGTGGGTTACATCGAACTGGATCTCAACAGCGGTAAGATCCTTGAGAGTTTTCGCCCC
GAAGAACGTTTTCCAATGATGAGCACTTTTAAAGTTCTGCTATGTGGCGCGGTATTATCCCGTATTGACGCCGGGCAAGAGC
AACTCGGTCGCCGCATACACTATTCTCAGAATGACTTGGTTGAGTACTCACCAGTCACAGAAAAGCATCTTACGGATGGCAT
GACAGTAAGAGAATTATGTAGTGCTGCCATAACCATGAGTGATAACACAGCGGCCAACTTACTTCTGACAACGATCGGAGGA
CCGAAGGAGCTAACCGCTTTTTTGCACAACATGGGGGATCATGTAACTCGCCTTGATCGTTGGGAACCGGAGCTGAATGAAG
CCATACCAAACGACGAGCGTGACACCACGATGCCTGTAGCGATGGCAACAACGTTGCGCAAACTATTAACTGGCGAACTACT
TACTCTAGCTTCCCGGCAACAATTAATAGACTGGATGGAGGCGGATAAAGTTGCAGGACCACTTCTTCGCTCAGCACTTCCA
GCTGGTTGGTTTATTGCTGATAAATCTGGAGCCGGTGAGCGTGGCTCTCGCGGTATCATTGCAGCACTGGGGCCAGATGGTA
AGCCCTCCCGTATCGTAGTTATCTACACGACGGGGAGTCAGGCAACTATGGATGAACGAAATAGACAGATCGCTGAGATAGG
TGCCTCACTGATTAAGCATTGGTAACTGTCAGACCAAGTTTACTCATATATACTTTAGATTGATTT**GGTCTC**ACGCGCCCTG
TAGCGGCGCATTAAGCGCGGCGGGTGTGGTGGTTACGCGCAGCGTGACCGCTACACTTGCCAGCGCCCTAGCGCCCGCTCCT
TTCGCTTTCTTCCCTTCCTTTCTCGCCACGTTCGCCGGCTTTCCCCGTCAAGCTCTAAATCGGGGGCTCCCTTTAGGGTTCC
GATTTAGTGCTTTACGGCACCTCGACCCCAAAAAACTTGATTTGGGTGATGGTTCACGTAGTGGGCCATCGCCCTGATAGAC
GGTTTTTCGCCCTTTGACGTTGGAGTCCACGTTCTTTAATAGTGGACTCTTGTTCCAAACTGGAACAACACTCAACCCTATC
TCGGGCTATTCTTTTGATTTATAAGGGATTTTGCCGATTTCGGCCTATTGGTTAAAAAATGAGCTGATTTAACAAAAATTTA
ACGCGAATTTTAACAAAATATTAACGTTTACAATTTAAAAGGATCTAGGTGAAGATCCTTTTTGATAATCTCATGACCAAAA
TCCCTTAACGTGAGTTTTCGTTCCACTGAGCGTCAGACCCCGTAGAAAAGATCAAAGGATCTTCTTGAGATCCTTTTTTTCT
GCGCGTAATCTGCTGCTTGCAAACAAAAAAACCACCGCTACCAGCGGTGGTTTGTTTGCCGGATCAAGAGCTACCAACTCTT
TTTCCGAAGGTAACTGGCTTCAGCAGAGCGCAGATACCAAATACTGTCCTTCTAGTGTAGCCGTAGTTAGGCCACCACTTCA
AGAACTCTGTAGCACCGCCTACATACCTCGCTCTGCTAATCCTGTTACCAGTGGCTGCTGCCAGTGGCGATAAGTCGTGTCT
TACCGGGTTGGACTCAAGACGATAGTTACCGGATAAGGCGCAGCGGTCGGGCTGAACGGGGGGTTCGTGCACACAGCCCAGC
TTGGAGCGAACGACCTACACCGAACTGAGATACCTACAGCGTGAGCTATGAGAAAGCGCCACGCTTCCCGAAGGGAGAAAGG
CGGACAGGTATCCGGTAAGCGGCAGGGTCGGAACAGGAGAGCGCACGAGGGAGCTTCCAGGGGGAAACGCCTGGTATCTTTA
TAGTCCTGTCGGGTTTCGCCACCTCTGACTTGAGCGTCGATTTTTGTGATGCTCGTCAGGGGGGCGGAGCCTATGGAAAAAC
GCCAGCAACGCGGCCTTTTTACGGTTCCTGGCCTTTTGCTGGCCTTTTGCTCACATGTTCTTTCCTGCGTTATCCCCTGATT
CTGTGGATAACCGTATTACCGCCTTTGAGTGAGCTGATACCGCTCGCCGCAGCCGAACGACCGAGCGCAGCGAGTCAGTGAG
CGAGGAAGCGGTAGAGCGCCTGATGCGGTATTTTCTCCTTACGCATCTGTGCGGTATTTCACACCGCATAGGGTCATGGCTG
CGCCCCGACACCCGCCAACACCCGCTGACGCGCCCTGACGGGCTTGTCTGCTCCCGGCATCCGCTTACAGACAAGCTGTGAC
CGTGTCCGGGAGCTGCATGTGTCAGAGGTTTTCACCGTCATCACCGAAACGCGCGAGGCAGAAGGAGATGGCGCCCAACAGT
CCCCCGGCCACGGGGCCTGCCACCATACCCACGCCGAAACAAGCGCTCATGAGCCCGAAGTGGCGAGCCCGATCTTCCCCAT
CGGTGATGTCGGCGATATAGGCGCCAGCAACCGCACCTGTGGCGCCGGTGATGCCGGCCACGATGCGTCCGGCGTAGAGGAT
CTGCTCATGTTTGACAGCTTATCATCGATTTATATTCCCCAGAACATCAGGTTAATGGCGTTTTTGATGTCATTTTCGCGGT
GGCTGAGATCAGCCACTTCTTCCCCGATAACGGAGACTGGCACACTGGCCATATCGGTGGTCATCATGCGCCAGCTTTCATC
CCCGATATGCACCACCGGGTAAAGTTCACGGGAGACTTTATCTGACAGCAGACGTGCACTGGCCAGGGGGATCACCATCCGT
CGCCCGGGCGTGTCAATAATATCACTCTGTACATCCACAAACAGACGATAACGGCTCTCTCTTTTATAGGTGTAAACCTTAA
ACTGCATAGCGCACCGCAAAGTTAAGAAACC

### Adaptor Strategy

*Combining iFLinkC-EZ with Golden Gate to Assemble Polycistronic Expression Constructs*

Large poly-cistronic expression constructs were assembled from individual translational units pre-assembled using iFLinkC-EZ. The destination plasmid pZT7-GoldenGate was based on pZHiX with an AMP resistance gene and ccdB for counterselection while individual translational units were assembled in pZ with a KAN resistant backbone. The total volume of a Golden Gate mediated assembly reaction was 25 µL. Adaptors for Golden Gate mediated assembly were introduced *via* iFLinkC-EZ at the 5' and 3' of each translation unit in $pZ^{KAN}$. pX vectors containing adaptors are equivalent to pL vectors. Adaptors are organized in roman numerals where I' + I, II' + II, III' + III, IV' + IV, X' and X denote pairs for DNA assembly. For Golden Gate mediated DNA assembly in pZT7-GoldenGate, the most 5' and the most 3' translation units strictly require a I and X' adaptor module. The BsmbI Golden Gate assembly reaction was composed of plasmid DNA featuring 100 ng each of the relevant translational unit, 2.5 µL T4 DNA Ligase Buffer (10×), 1 µL T4 DNA Ligase (400 units/µL) and 1 µL BsmbI-v2 (10 units/µL) before being subject to the temperature cycling protocol outlined in **Tab. S3**. Upon completion of the Golden Gate cycling protocol, 10 µL reaction mix was transformed into 100 µL $RbCl_2$ competent cells DH10B cells and plated on LB agar plates (+ AMP). Any undigested pZT7-GoldenGate with an AMP resistance was negatively selected using ccdB while pZ with a KAN resistance coding for individual translation units were negatively selected *via* a change in the antibiotic resistance from kanamycin to ampicillin.

**Table S3.** Temperature Cycling Protocol for BsmBI mediated Golden Gate assembly

| **Step** | **Temperature** | **Duration** | **Cycles** |
|---|---|---|---|
| Restriction digest | 42 °C | 5 min | 20 |
| T4 DNA Ligase | 16 °C | 5 min | |
| Restriction digest at opt. temperature for BsmBI | 55 °C | 10 min | 1 |
| Add 0.5 µL T5 Exonuclease | 37 °C | Pause | 1 |
| Degrade linear DANN | 37 °C | 30 min | 1 |
| Heat inactivate enzymes | 80 °C | 20 min | 1 |

**Destination plasmid pZT7-GoldenGate**

GG I' site *(BsmBI recognition* and cleavage), ccdB, GG X site (*BsmBI recognition* and cleavage), AmpR

GGTGCATGCAAGGAGATGGTAGAGGATCGAGATCTCGATCCCGCGAAATTAATACGACTCACTATAGGGAGAGGAATTGTGA
GCGGATAACAATTCCCCTCTAGAAATAATCATAG*GAGACG*TTTAAATTAAAGAGGAGAAATACATTATGCAGTTTAAGGTTT
ACACCTATAAAAGAGAGAGCCGTTATCGTCTGTTTGTGGATGTACAGAGTGATATTATTGACACGCCCGGGCGACGGATGGT
GATCCCCCTGGCCAGTGCACGTCTGCTGTCAGATAAAGTCTCCCGTGAACTTTACCCGGTGGTGCATATCGGGGATGAAAGC
TGGCGCATGATGACCACCGATATGGCCAGTGTGCCGGTCTCCGTTATCGGGGAAGAAGTGGCTGATCTCAGCCACCGCGAAA
ATGACATCAAAAACGCCATTAACCTGATGTTCTGGGGAATATAACAT*CGTCTCC*TCGATAAAGAGCGTCGACCTGCAGGCAT
GCAAGCTTGGCTGTTTTGGCGGATGAGAGAAGATTTTCAGCCTGATACAGATTAAATCAGAACGCAGAAGCGGTCTGATAAA
ACAGAATTTGCCTGGCGGCAGTAGCGCGGTGGTCCCACCTGACCCCATGCCGAACTCAGAAGTGAAACGCCGTAGCGCCATG
GTAGTGTGGCCAGAGCCCATGCGAGAGTAGGGAACTGCCAGGCATCAAATAAAACGAAAGGCTCAGTCGAAAGACTGGGCCT
TTCGTTTTATCTGTTGTTTGTCGGTGAACGCTCTCCTGAGTAGGACAAATCCGCCGGGAGCGGATTTGAACGTTGCGAAGCA
ACGGCCCGGAGGGTGGCGGGCAGGACGCCCGCCATAAACTGCCAGGCATCAAATTAAGCAGAAGGCCATCCTGACGGATGGC
CTTTTTGCGTTTCTACAAACTCTTTTGTTTATTTTTCTAAATACATTCAAATATGTATCCGCTCATGAGACAATAACCCTGA
TAAATGCTTCAATAATATTGAAAAAGGAAGTGTATGAGTATTCAACATTTCCGTGTCGCCCTTATTCCCTTTTTTGCGGCAT
TTTGCCTTCCTGTTTTTGCTCACCCAGAAACGCTGGTGAAAGTAAAAGATGCTGAAGATCAGTTGGGTGCACGAGTGGGTTA
CATCGAACTGGATCTCAACAGCGGTAAGATCCTTGAGAGTTTTCGCCCCGAAGAACGTTTTCCAATGATGAGCACTTTTAAA
GTTCTGCTATGTGGCGCGGTATTATCCCGTATTGACGCCGGGCAAGAGCAACTCGGTCGCCGCATACACTATTCTCAGAATG
ACTTGGTTGAGTACTCACCAGTCACAGAAAAGCATCTTACGGATGGCATGACAGTAAGAGAATTATGTAGTGCTGCCATAAC
CATGAGTGATAACACAGCGGCCAACTTACTTCTGACAACGATCGGAGGACCGAAGGAGCTAACCGCTTTTTTGCACAACATG
GGGGATCATGTAACTCGCCTTGATCGTTGGGAACCGGAGCTGAATGAAGCCATACCAAACGACGAGCGTGACACCACGATGC
CTGTAGCGATGGCAACAACGTTGCGCAAACTATTAACTGGCGAACTACTTACTCTAGCTTCCCGGCAACAATTAATAGACTG
GATGGAGGCGGATAAAGTTGCAGGACCACTTCTTCGCTCAGCACTTCCAGCTGGTTGGTTTATTGCTGATAAATCTGGAGCC
GGTGAGCGTGGCTCTCGCGGTATCATTGCAGCACTGGGGCCAGATGGTAAGCCCTCCCGTATCGTAGTTATCTACACGACGG
GGAGTCAGGCAACTATGGATGAACGAAATAGACAGATCGCTGAGATAGGTGCCTCACTGATTAAGCATTGGTAACTGTCAGA
CCAAGTTTACTCATATATACTTTAGATTGATTTGGTCTCACGCGCCCTGTAGCGGCGCATTAAGCGCGGCGGGTGTGGTGGT
TACGCGCAGCGTGACCGCTACACTTGCCAGCGCCCTAGCGCCCGCTCCTTTCGCTTTCTTCCCTTCCTTTCTCGCCACGTTC
GCCGGCTTTCCCCGTCAAGCTCTAAATCGGGGGCTCCCTTTAGGGTTCCGATTTAGTGCTTTACGGCACCTCGACCCCAAAA
AACTTGATTTGGGTGATGGTTCACGTAGTGGGCCATCGCCCTGATAGACGGTTTTTCGCCCTTTGACGTTGGAGTCCACGTT
CTTTAATAGTGGACTCTTGTTCCAAACTGGAACAACACTCAACCCTATCTCGGGCTATTCTTTTGATTTATAAGGGATTTTG
CCGATTTCGGCCTATTGGTTAAAAAATGAGCTGATTTAACAAAAATTTAACGCGAATTTTAACAAAATATTAACGTTTACAA
TTTAAAAGGATCTAGGTGAAGATCCTTTTTGATAATCTCATGACCAAAATCCCTTAACGTGAGTTTTCGTTCCACTGAGCGT
CAGACCCCGTAGAAAAGATCAAAGGATCTTCTTGAGATCCTTTTTTTCTGCGCGTAATCTGCTGCTTGCAAACAAAAAAACC
ACCGCTACCAGCGGTGGTTTGTTTGCCGGATCAAGAGCTACCAACTCTTTTTCCGAAGGTAACTGGCTTCAGCAGAGCGCAG
ATACCAAATACTGTCCTTCTAGTGTAGCCGTAGTTAGGCCACCACTTCAAGAACTCTGTAGCACCGCCTACATACCTCGCTC
TGCTAATCCTGTTACCAGTGGCTGCTGCCAGTGGCGATAAGTCGTGTCTTACCGGGTTGGACTCAAGACGATAGTTACCGGA
TAAGGCGCAGCGGTCGGGCTGAACGGGGGGTTCGTGCACACAGCCCAGCTTGGAGCGAACGACCTACACCGAACTGAGATAC
CTACAGCGTGAGCTATGAGAAAGCGCCACGCTTCCCGAAGGGAGAAAGGCGGACAGGTATCCGGTAAGCGGCAGGGTCGGAA
CAGGAGAGCGCACGAGGGAGCTTCCAGGGGGAAACGCCTGGTATCTTTATAGTCCTGTCGGGTTTCGCCACCTCTGACTTGA
GCGTCGATTTTTGTGATGCTCGTCAGGGGGGCGGAGCCTATGGAAAAACGCCAGCAACGCGGCCTTTTTACGGTTCCTGGCC
TTTTGCTGGCCTTTTGCTCACATGTTCTTTCCTGCGTTATCCCCTGATTCTGTGGATAACCGTATTACCGCCTTTGAGTGAG
CTGATACCGCTCGCCGCAGCCGAACGACCGAGCGCAGCGAGTCAGTGAGCGAGGAAGCGGTAGAGCGCCTGATGCGGTATTT
TCTCCTTACGCATCTGTGCGGGAGATCCCGGTGCCTAATGAGTGAGCTAACTTACATTAATTGCGTTGCGCTCATTGACCGC
TTTCCAGTCGGGAAACCTGTCGTGCCAGCTGCATTAATGAATCGGCCAACGCGCGGGGAGAGGCGGTTTGCGTATTGGGCGC
CAGGGTGGTTTTTCTTTTCACCAGTGAAACGGGCAACAGCTGATTGCCCTTCACCGCCTGGCCCTGAGAGAGTTGCAGCAAG
CGGTCCACGCTGGTTTGCCCCAGCAGGCGAAAATCCTGTTTGATGGTGGTTAACGGCGGGATATAACATGAGCTGTCCTCGG
TATCGTCGTATCCCACTACCGAGATGTCCGCACCAACGCGCAGCCCGGACTCGGTAATGGCGCGCATTGCGCCCAGCGCCAT
CTGATCGTTGGCAACCAGCATCGCGGTTGGAACGATGCCCTCATTCAGCATTTGCATGGTTTGTTGAAAACCGGACATGGCA
CTAAAGTCGCCTTCCCGTTCCGCTATCGGCTGAATTTGATTGCGAGTGAGATATTTATGCCAGCCAGCCAGACGCAGACGCG
CCGAGACAGAACTTAATGGGCCCGCTAACAGCGCGATTTGCTGGTGACCCAATGCGACCAGATGCTCCACGCCCAGTCGCGT
ACCATCTTCATGGGAGAAAATAATACTGTTGATGGGTGTCTGGTCAGAGACATCAAGAAATAACGCCGGAACATTAGTGCAG
GCAGCTTCCACAGCAATGGCATCCTGGTCATCCAGCGGATAGTTAATGATCAGCCCACTGACGCGTTGCGCGAGAAGATTGT
GCACCGCCGCTTTACAGGCTTCGACGCCGCTTCGTTCTACCATCGACACCACCACGCTGGCACCCAGTTGATCGGCGCGAGA
TTTAATCGCCGCGACAATTTGCGACGGCGCGTGCAGGGCCAGACTGGAGGTGGCAACGCCAATCAGCAACGACTGTTTGCCC
GCCAGTTGTTGTGCCACGCGGTTGGGAATGTAATTCAGCTCCGCCATCGCCGCTTCCACTTTTTCCCGCGTTTTCGCAGAAA
CGTGGCTGGCCTGGTTCACCACGCGGGAAACGGTCTGATAAGAGACACCGGCATACTCTGCGACATCGTATAACGTTACTGG
TTTCACATTCACCACCCTGAATTGACTCTCTTCCGGGCGCTATCATGCCATACCGCGAAAGGTTTTGCGCCATTCGATGGTG
TCCGGGATCTCGACGCTCTCCCTTATGCGACTCCTGCATTAGG

#### *Adaptor 1* ###

Strictly required at the 5’ end of the most 5’ translation unit for recombination into pZT7-GoldenGate.

**pZ-MBP-TU(I)-sRBS |** MBP, GG I site (*BsmBI*, BsmBI), sRBS, BtsI, KanR, **BbsI**,

GAATATTGGGTTTAGTCTTGTTTCATAATTGTTGCAATGAAACGCGGTGAAACATTGCCTGAAACGTTAACTGAAACGCATA
TTTGCGGATTAGTTCATGACTTTATCTCTAACAAATTGAAATTAAACATTTAATTTTATTAAGGCAATTGTGGCACACCCCT
TGCTTTGTCTTTATCAACGCAAATAACAAGTTGATAACAAGCTAGCAGGAGGAATTCCATATGGGGAAAATCGAAGAAGGTA
AACTGGTAATCTGGATTAACGGCGATAAAGGCTATAACGGTCTGGCTGAAGTCGGTAAGAAATTCGAGAAAGATACCGGAAT
TAAAGTCACCGTTGAGCATCCGGATAAACTGGAAGAGAAATTCCCACAGGTTGCGGCAACTGGCGATGGCCCTGACATTATC
TTCTGGGCACACGACCGCTTTGGTGGCTACGCTCAATCTGGCCTGTTGGCTGAAATCACCCCGGACAAAGCGTTCCAGGACA
AGCTGTATCCGTTTACCTGGGATGCCGTACGTTACAACGGCAAGCTGATTGCTTACCCGATCGCTGTTGAAGCGTTATCGCT
GATTTATAACAAAGATCTGCTGCCGAACCCGCCAAAAACCTGGGAAGAGATCCCGGCGCTGGATAAAGAACTGAAAGCGAAA
GGTAAGAGCGCGCTGATGTTCAACCTGCAAGAACCGTACTTCACCTGGCCGCTGATTGCTGCTGACGGGGGTTATGCGTTCA
AGTATGAAAACGGCAAGTACGACATTAAAGACGTGGGCGTGGATAACGCTGGCGCGAAAGCGGGTCTGACCTTCCTGGTTGA
CCTGATTAAAAACAAACACATGAATGCAGACACCGATTACTCCATCGCAGAAGCTGCCTTTAATAAAGGCGAAACAGCGATG
ACCATCAACGGCCCGTGGGCATGGTCCAACATCGACACCAGCAAAGTGAATTATGGTGTAACGGTACTGCCGACCTTCAAGG
GTCAACCATCCAAACCGTTCGTTGGCGTGCTGAGCGCAGGTATTAACGCCGCCAGTCCGAACAAAGAGCTGGCAAAAGAGTT
CCTCGAAAACTATCTGCTGACTGATGAAGGTCTGGAAGCGGTTAATAAAGACAAACCGCTGGGTGCCGTAGCGCTGAAGTCT
TACGAGGAAGAGTTGGTGAAAGATCCGCGTATTGCCGCCACTATGGAAAACGCCCAGAAAGGTGAAATCATGCCGAACATCC
CGCAGATGTCCGCTTTCTGGTATGCCGTGCGTACTGCGGTGATCAACGCCGCCAGCGGTCGTCAGACTGTCGATGAAGCCCT
GAAAGACGCGCAGACTGGGTAA*CGTCTC*ACATAATTTAAAAAGAGGAGAAAGTACATTATGGGGCACTGCTGAACTAGTCTG
ATACAGTCGACCTGCAGGCATGCAAGCTTGGCTGTTTTGGCGGATGAGAGAAGATTTTCAGCCTGATACAGATTAAATCAGA
ACGCAGAAGCGGTCTGATAAAACAGAATTTGCCTGGCGGCAGTAGCGCGGTGGTCCCACCTGACCCCATGCCGAACTCAGAA
GTGAAACGCCGTAGCGCCGATGGTAGTGTGGCCAGAGCCCATGCGAGAGTAGGGAACTGCCAGGCATCAAATAAAACGAAAG
GCTCAGTCGAAAGACTGGGCCTTTCGTTTTATCTGTTGTTTGTCGGTGAACGCTCTCCTGAGTAGGACAAATCCGCCGGGAG
CGGATTTGAACGTTGCGAAGCAACGGCCCGGAGGGTGGCGGGCAGGACGCCCGCCATAAACTGCCAGGCATCAAATTAAGCA
GAAGGCCATCCTGACGGATGGCCTTTTTGCGTTTCTACAAACTCTCTGATCCTTCAACTCAGCAAAAGTTCGATTTATTCAA
CAAAGCCACGTTGTGTCTCAAAATCTCTGATGTTACATTGCACAAGATAAAAATATATCATCATGAACAATAAAACTGTCTG
CTTACATAAACAGTAATACAAGGGGTGTTATGAGCCATATTCAACGGGAAACGTCTTGCTCTAGGCCGCGATTAAATTCCAA
CATGGATGCTGATTTATATGGGTATAAATGGGCTCGCGATAATGTCGGGCAATCAGGTGCGACAATCTATCGATTGTATGGG
AAGCCCGATGCGCCAGAGTTGTTTCTGAAACATGGCAAAGGTAGCGTTGCCAATGATGTTACAGATGAGATGGTCAGACTAA
ACTGGCTGACGGAATTTATGCCTCTGCCGACCATCAAGCATTTTATCCGTACTCCTGATGATGCATGGTTACTCACCACGGC
GATCCCAGGAAAACAGCATTCCAGGTATTAGAAGAATATCCTGATTCAGGTGAAAATATTGTTGATGCGCTGGCGGTGTTC
CTGCGCCGGTTGCATTCGATTCCTGTTTGTAATTGTCCTTTTAACAGCGACCGCGTATTTCGTCTGGCTCAGGCGCAATCAC
GAATGAATAACGGTTTGGTTGATGCGAGTGATTTTGATGACGAGCGTAATGGCTGGCCTGTTGAACAAGTCTGGAAAGAAAT
GCATAAACTTTTGCCATTCTCACCGGATTCAGTCGTCACTCATGGTGATTTCTCACTTGATAACCTTATTTTTGACGAGGGG
AAATTAATAGGTTGTATTGATGTTGGACGAGTCGGAATCGCAGACCGATACCAGGATCTTGCCATCCTATGGAACTGCCTCG
GTGAGTTTTCTCCTTCATTACAGAAACGGCTTTTTCAAAAATATGGTATTGATAATCCTGATATGAATAAATTGCAGTTTCA
TTTGATGCTCGATGAGTTTTTCTAACTGTCAGACCAAGTTTACTCATATATACTTTAGATTGATTT**GAAGAC**TACGCGCCCT
GTAGCGGCGCATTAAGCGCGGCGGGTGTGGTGGTTACGCGCAGCGTGACCGCTACACTTGCCAGCGCCCTAGCGCCCGCTCC
TTTCGCTTTCTTCCCTTCCTTTCTCGCCACGTTCGCCGGCTTTCCCCGTCAAGCTCTAAATCGGGGGCTCCCTTTAGGGTTC
CGATTTAGTGCTTTACGGCACCTCGACCCCAAAAAACTTGATTTGGGTGATGGTTCACGTAGTGGGCCATCGCCCTGATAGA
CGGTTTTTCGCCCTTTGACGTTGGAGTCCACGTTCTTTAATAGTGGACTCTTGTTCCAAACTGGAACAACACTCAACCCTAT
CTCGGGCTATTCTTTTGATTTATAAGGGATTTTGCCGATTTCGGCCTATTGGTTAAAAAATGAGCTGATTTAACAAAAATTT
AACGCGAATTTTAACAAAATATTAACGTTTACAATTTAAAAGGATCTAGGTGAAGATCCTTTTTGATAATCTCATGACCAAA
ATCCCTTAACGTGAGTTTTCGTTCCACTGAGCGTCAGACCCCGTAGAAAAGATCAAAGGATCTTCTTGAGATCCTTTTTTTC
TGCGCGTAATCTGCTGCTTGCAAACAAAAAAACCACCGCTACCAGCGGTGGTTTGTTTGCCGGATCAAGAGCTACCAACTCT
TTTTCCGAAGGTAACTGGCTTCAGCAGAGCGCAGATACCAAATACTGTCCTTCTAGTGTAGCCGTAGTTAGGCCACCACTTC
AAGAACTCTGTAGCACCGCCTACATACCTCGCTCTGCTAATCCTGTTACCAGTGGCTGCTGCCAGTGGCGATAAGTCGTGTC
TTACCGGGTTGGACTCAAGACGATAGTTACCGGATAAGGCGCAGCGGTCGGGCTGAACGGGGGGTTCGTGCACACAGCCCAG
CTTGGAGCGAACGACCTACACCGAACTGAGATACCTACAGCGTGAGCTATGAGAAAGCGCCACGCTTCCCGAAGGGAGAAAG
GCGGACAGGTATCCGGTAAGCGGCAGGGTCGGAACAGGAGAGCGCACGAGGGAGCTTCCAGGGGGAAACGCCTGGTATCTTT
ATAGTCCTGTCGGGTTTCGCCACCTCTGACTTGAGCGTCGATTTTTGTGATGCTCGTCAGGGGGGCGGAGCCTATGGAAAAA
CGCCAGCAACGCGGCCTTTTTACGGTTCCTGGCCTTTTGCTGGCCTTTTGCTCACATGTTCTTTCCTGCGTTATCCCCTGAT
TCTGTGGATAACCGTATTACCGCCTTTGAGTGAGCTGATACCGCTCGCCGCAGCCGAACGACCGAGCGCAGCGAGTCAGTGA
GCGAGGAAGCGGTAGAGCGCCTGATGCGGTATTTTCTCCTTACGCATCTGTGCGGTATTTCACACCGCATAGGGTCATGGCT

GCGCCCCGACACCCGCCAACACCCGCTGACGCGCCCTGACGGGCTTGTCTGCTCCCGGCATCCGCTTACAGACAAGCTGTGA
CCGTGTCCGGGAGCTGCATGTGTCAGAGGTTTTCACCGTCATCACCGAAACGCGCGAGGCAGAAGGAGATGGCGCCCAACAG
TCCCCCGGCCACGGGGCCTGCCACCATACCCACGCCGAAACAAGCGCTCATGAGCCCGAAGTGGCGAGCCCGATCTTCCCCA
TCGGTGATGTCGGCGATATAGGCGCCAGCAACCGCACCTGTGGCGCCGGTGATGCCGGCCACGATGCGTCCGGCGTAGAGGA
TCTGCTCATGTTTGACAGCTTATCATCGATTCAGCTTTTCAGCCGCCGCCAGAACGTCGTCCGGCTGATGCCTAAATAATTC
GCCGCTGCTGTTTTATCGCCATTAAATTTCTCCAGTGCCTGTTGTGGTGTCAGTAAGCGTGGAGCGGGAGTTTTCGCCGACT
CGCGCGCCAGTTCCGGCAGTAGCAGCTGCAAAAATTGCGGCGTTAAATCCGGCGTCGGTTCCACACTTAAAAATAGCGCCAG
TCGCTCCATCATATTGCGCAGTTCACGAATATTGCCCGGCCAGTCGTAGTGCACCAGCACGGTTTCGCTTGCCTGTAATCCC
TGGCGTAATGCGGCAGAAAACGGGGTGGAGAGCGCCGCCAGAGACACTTTCAAAAAGCTTTCCGCCAGTGGCAGAATATCCG
CCACCCGCTCGCGCAGCGGTGGCAATTGCAGACGCAAAATACTCAGCCGATAAAACAGGTCACGGCGAAACTGCCCTTGCCG
CATATCTTCTTCCAGATTACAGTGAGTGGCGCTAATGACCCGCACATCCACCGGAACAGGCTGATGCCCGCCGACGCGGGTG
ACCTCTTTTTCTTCCAGCACCCGCAGCAGCCGGGTCTGCAACGGCAGCGGCATTTCGCCAATCTCATCGAGAAACAGCGTAC
CTCCGTGGGCGATTTCAAACAGCCCGGCGCGACCGCCGCGTCGCGAGCCGGTAAACGCCCCTTCCTCATAGCCAAACAGTTC
TGCTTCCAGCAGCGATTCGGCAATCGCCCCGCAGTTGACTGCAACAAACGGATGCGACTTTTTGCCCTGTCGCGCATCGTGG
CGGGCAAAATATTCCCGATGAATCGCCTGGGCCGCCAGCTCTTTGCCCGTCCCCGTTTCCCCCTCAATCAACACCGCTGCAC
TGGAGCGGGCATACAGCAAAATAGTCTGCCGTACTTGTTCCATCTGTGGTGATTGACCGAGCATATCGCCCAGCACGTAACG
AGTTCTCAGGGCGTTGCGGGTGGCATCGTGAGTGTTATGGCGTAACGACATGCGCGTCATATCCAGCGCATCGCTGAACGCC
TGGCGCACGGTGGCGGCGGAATAGATAAAAATTCCGGTCATTCCGGCTTCTTCTGCCAGATCGGTAATCAGCCCCGCGCCGA
CCACCGCTTCGGTGCCGTTAGCTTTTAGCTCGTTAATCTGCCCGCGTGCATCTTCTTCGGTAATGTAGCTACGTTGATCGAG
GCGCAAATTAAAGGTTTTTTGAAACGCCACCAGTGCCGGAATAGTTTCCTGATAAGTGACAACGCCGATCGAGCTGGTGAGT
TTTCCGGCTTTTGCCAGTGCCTGTAACACATCGTAGCCGCTCGGTTTAATCAAAATAACTGGCACTGACAGGCGGCTTTTCA
GGTACGCGCCGTTAGATCCAGCGGCGATGATGGCGTCACAGCGTTCGTTTGCCAGTTTCTTGTGGATGTAGGTCACCGCTTT
TTCAAAGCCAAGCTGGATAGGGGTAATGTTCGCCAGGTGATCAAACTCGAGGCTGATATCGCGAAACAGCTCGAACAGGCGC
GTTACAGATACCGTCCAGATAACCGGTTTGTCGTCATTAAGCCGTGGTGGATGTGCCATAGCGCACCGCAAAGTTAAGAAAC
C

**### *Adaptor 2* ###**

**pX-TU(II') |** SapI, GG II' site (*BsmBI*, BsmBI), BtsI, KanR, **BbsI**, ccdB

GAATATTGGGTTTAGTCTTGTTTCATAATTGTTGCAATGAAACGCGGTGAAACATTGCCTGAAACGTTAACTGAAACGCATA
TTTGCGGATTAGTTCATGACTTTATCTCTAACAAATTGAAATTAAACATTTAATTTTATTAAGGCAATTGTGGCACACCCCT
TGCTTTGTCTTTATCAACGCAAATAACAAGTTGATAACAAGCTAGCAGGAGGAATTCCATATGGGCTCTTCAGGGTAAGGAG
T*GAGACG*TGGGGCACTGCTGAACTAGTCTGATACAGTCGACCTGCAGGCATGCAAGCTTGGCTGTTTTGGCGGATGAGAGAA
GATTTTCAGCCTGATACAGATTAAATCAGAACGCAGAAGCGGTCTGATAAAACAGAATTTGCCTGGCGGCAGTAGCGCGGTG
GTCCCACCTGACCCCATGCCGAACTCAGAAGTGAAACGCCGTAGCGCCGATGGTAGTGTGGCCAGAGCCCATGCGAGAGTAG
GGAACTGCCAGGCATCAAATAAAACGAAAGGCTCAGTCGAAAGACTGGGCCTTTCGTTTTATCTGTTGTTTGTCGGTGAACG
CTCTCCTGAGTAGGACAAATCCGCCGGGAGCGGATTTGAACGTTGCGAAGCAACGGCCCGGAGGGTGGCGGGCAGGACGCCC
GCCATAAACTGCCAGGCATCAAATTAAGCAGAAGGCCATCCTGACGGATGGCCTTTTTGCGTTTCTACAAACTCTCTGATCC
TTCAACTCAGCAAAAGTTCGATTTATTCAACAAAGCCACGTTGTGTCTCAAAATCTCTGATGTTACATTGCACAAGATAAAA
ATATATCATCATGAACAATAAAACTGTCTGCTTACATAAACAGTAATACAAGGGGTGTTATGAGCCATATTCAACGGGAAAC
GTCTTGCTCTAGGCCGCGATTAAATTCCAACATGGATGCTGATTTATATGGGTATAAATGGGCTCGCGATAATGTCGGGCAA
TCAGGTGCGACAATCTATCGATTGTATGGGAAGCCCGATGCGCCAGAGTTGTTTCTGAAACATGGCAAAGGTAGCGTTGCCA
ATGATGTTACAGATGAGATGGTCAGACTAAACTGGCTGACGGAATTTATGCCTCTGCCGACCATCAAGCATTTTATCCGTAC
TCCTGATGATGCATGGTTACTCACCACGGCGATCCCAGGGAAAACAGCATTCCAGGTATTAGAAGAATATCCTGATTCAGGT
GAAAATATTGTTGATGCGCTGGCGGTGTTCCTGCGCCGGTTGCATTCGATTCCTGTTTGTAATTGTCCTTTTAACAGCGACC
GCGTATTTCGTCTGGCTCAGGCGCAATCACGAATGAATAACGGTTTGGTTGATGCGAGTGATTTTGATGACGAGCGTAATGG
CTGGCCTGTTGAACAAGTCTGGAAAGAAATGCATAAACTTTTGCCATTCTCACCGGATTCAGTCGTCACTCATGGTGATTTC
TCACTTGATAACCTTATTTTTGACGAGGGGAAATTAATAGGTTGTATTGATGTTGGACGAGTCGGAATCGCAGACCGATACC
AGGATCTTGCCATCCTATGGAACTGCCTCGGTGAGTTTTCTCCTTCATTACAGAAACGGCTTTTTCAAAAATATGGTATTGA
TAATCCTGATATGAATAAATTGCAGTTTCATTTGATGCTCGATGAGTTTTTCTAACTGTCAGACCAAGTTTACTCATATATA
CTTTAGATTGATTT**GAAGAC**TACGCGCCCTGTAGCGGCGCATTAAGCGCGGCGGGTGTGGTGGTTACGCGCAGCGTGACCGC
TACACTTGCCAGCGCCCTAGCGCCCGCTCCTTTCGCTTTCTTCCCTTCCTTTCTCGCCACGTTCGCCGGCTTTCCCCGTCAA
GCTCTAAATCGGGGGCTCCCTTTAGGGTTCCGATTTAGTGCTTTACGGCACCTCGACCCCAAAAAACTTGATTTGGGTGATG
GTTCACGTAGTGGGCCATCGCCCTGATAGACGGTTTTTCGCCCTTTGACGTTGGAGTCCACGTTCTTTAATAGTGGACTCTT
GTTCCAAACTGGAACAACACTCAACCCTATCTCGGGCTATTCTTTTGATTTATAAGGGATTTTGCCGATTTCGGCCTATTGG
TTAAAAAATGAGCTGATTTAACAAAAATTTAACGCGAATTTTAACAAAATATTAACGTTTACAATTTAAAAGGATCTAGGTG
AAGATCCTTTTTGATAATCTCATGACCAAAATCCCTTAACGTGAGTTTTCGTTCCACTGAGCGTCAGACCCCGTAGAAAAGA
TCAAAGGATCTTCTTGAGATCCTTTTTTTCTGCGCGTAATCTGCTGCTTGCAAACAAAAAAACCACCGCTACCAGCGGTGGT
TTGTTTGCCGGATCAAGAGCTACCAACTCTTTTTCCGAAGGTAACTGGCTTCAGCAGAGCGCAGATACCAAATACTGTCCTT
CTAGTGTAGCCGTAGTTAGGCCACCACTTCAAGAACTCTGTAGCACCGCCTACATACCTCGCTCTGCTAATCCTGTTACCAG
TGGCTGCTGCCAGTGGCGATAAGTCGTGTCTTACCGGGTTGGACTCAAGACGATAGTTACCGGATAAGGCGCAGCGGTCGGG
CTGAACGGGGGGTTCGTGCACACAGCCCAGCTTGGAGCGAACGACCTACACCGAACTGAGATACCTACAGCGTGAGCTATGA
GAAAGCGCCACGCTTCCCGAAGGGAGAAAGGCGGACAGGTATCCGGTAAGCGGCAGGGTCGGAACAGGAGAGCGCACGAGGG
AGCTTCCAGGGGGAAACGCCTGGTATCTTTATAGTCCTGTCGGGTTTCGCCACCTCTGACTTGAGCGTCGATTTTTGTGATG
CTCGTCAGGGGGGCGGAGCCTATGGAAAAACGCCAGCAACGCGGCCTTTTTACGGTTCCTGGCCTTTTGCTGGCCTTTTGCT
CACATGTTCTTTCCTGCGTTATCCCCTGATTCTGTGGATAACCGTATTACCGCCTTTGAGTGAGCTGATACCGCTCGCCGCA
GCCGAACGACCGAGCGCAGCGAGTCAGTGAGCGAGGAAGCGGTAGAGCGCCTGATGCGGTATTTTCTCCTTACGCATCTGTG
CGGTATTTCACACCGCATAGGGTCATGGCTGCGCCCCGACACCCGCCAACACCCGCTGACGCGCCCTGACGGGCTTGTCTGC
TCCCGGCATCCGCTTACAGACAAGCTGTGACCGTGTCCGGGAGCTGCATGTGTCAGAGGTTTTCACCGTCATCACCGAAACG
CGCGAGGCAGAAGGAGATGGCGCCCAACAGTCCCCCGGCCACGGGGCCTGCCACCATACCCACGCCGAAACAAGCGCTCATG
AGCCCGAAGTGGCGAGCCCGATCTTCCCCATCGGTGATGTCGGCGATATAGGCGCCAGCAACCGCACCTGTGGCGCCGGTGA
TGCCGGCCACGATGCGTCCGGCGTAGAGGATCTGCTCATGTTTGACAGCTTATCATCGATTTATATTCCCCAGAACATCAGG
TTAATGGCGTTTTTGATGTCATTTTCGCGGTGGCTGAGATCAGCCACTTCTTCCCCGATAACGGAGACTGGCACACTGGCCA
TATCGGTGGTCATCATGCGCCAGCTTTCATCCCCGATATGCACCACCGGGTAAAGTTCACGGGAGACTTTATCTGACAGCAG
ACGTGCACTGGCCAGGGGGATCACCATCCGTCGCCCGGGCGTGTCAATAATATCACTCTGTACATCCACAAACAGACGATAA
CGGCTCTCTCTTTTATAGGTGTAAACCTTAAACTGCATAGCGCACCGCAAAGTTAAGAAACC

**pZ-MBP-TU(II)-sRBS |** MBP, GG II site (*BsmBI*, BsmBI), sRBS, BtsI, KanR, **BbsI**

GAATATTGGGTTTAGTCTTGTTTCATAATTGTTGCAATGAAACGCGGTGAAACATTGCCTGAAACGTTAACTGAAACGCATA
TTTGCGGATTAGTTCATGACTTTATCTCTAACAAATTGAAATTAAACATTTAATTTTATTAAGGCAATTGTGGCACACCCCT
TGCTTTGTCTTTATCAACGCAAATAACAAGTTGATAACAAGCTAGCAGGAGGAATTCCATATGGGGAAAATCGAAGAAGGTA
AACTGGTAATCTGGATTAACGGCGATAAAGGCTATAACGGTCTGGCTGAAGTCGGTAAGAAATTCGAGAAAGATACCGGAAT
TAAAGTCACCGTTGAGCATCCGGATAAACTGGAAGAGAAATTCCCACAGGTTGCGGCAACTGGCGATGGCCCTGACATTATC
TTCTGGGCACACGACCGCTTTGGTGGCTACGCTCAATCTGGCCTGTTGGCTGAAATCACCCCGGACAAAGCGTTCCAGGACA
AGCTGTATCCGTTTACCTGGGATGCCGTACGTTACAACGGCAAGCTGATTGCTTACCCGATCGCTGTTGAAGCGTTATCGCT
GATTTATAACAAAGATCTGCTGCCGAACCCGCCAAAAACCTGGGAAGAGATCCCGGCGCTGGATAAAGAACTGAAAGCGAAA
GGTAAGAGCGCGCTGATGTTCAACCTGCAAGAACCGTACTTCACCTGGCCGCTGATTGCTGCTGACGGGGGTTATGCGTTCA
AGTATGAAAACGGCAAGTACGACATTAAAGACGTGGGCGTGGATAACGCTGGCGCGAAAGCGGGTCTGACCTTCCTGGTTGA
CCTGATTAAAAACAAACACATGAATGCAGACACCGATTACTCCATCGCAGAAGCTGCCTTTAATAAAGGCGAAACAGCGATG
ACCATCAACGGCCCGTGGGCATGGTCCAACATCGACACCAGCAAAGTGAATTATGGTGTAACGGTACTGCCGACCTTCAAGG
GTCAACCATCCAAACCGTTCGTTGGCGTGCTGAGCGCAGGTATTAACGCCGCCAGTCCGAACAAAGAGCTGGCAAAAGAGTT
CCTCGAAAACTATCTGCTGACTGATGAAGGTCTGGAAGCGGTTAATAAAGACAAACCGCTGGGTGCCGTAGCGCTGAAGTCT
TACGAGGAAGAGTTGGTGAAAGATCCGCGTATTGCCGCCACTATGGAAAACGCCCAGAAAGGTGAAATCATGCCGAACATCC
CGCAGATGTCCGCTTTCTGGTATGCCGTGCGTACTGCGGTGATCAACGCCGCCAGCGGTCGTCAGACTGTCGATGAAGCCCT
GAAAGACGCGCAGACTGGGTAA*CGTCTC*AGGAGATTTAAAAAGAGGAGAAAGTACATTATGGGGCACTGCTGAACTAGTCTG
ATACAGTCGACCTGCAGGCATGCAAGCTTGGCTGTTTTGGCGGATGAGAGAAGATTTTCAGCCTGATACAGATTAAATCAGA
ACGCAGAAGCGGTCTGATAAAACAGAATTTGCCTGGCGGCAGTAGCGCGGTGGTCCCACCTGACCCCATGCCGAACTCAGAA
GTGAAACGCCGTAGCGCCGATGGTAGTGTGGCCAGAGCCCATGCGAGAGTAGGGAACTGCCAGGCATCAAATAAAACGAAAG
GCTCAGTCGAAAGACTGGGCCTTTCGTTTTATCTGTTGTTTGTCGGTGAACGCTCTCCTGAGTAGGACAAATCCGCCGGGAG
CGGATTTGAACGTTGCGAAGCAACGGCCCGGAGGGTGGCGGGCAGGACGCCCGCCATAAACTGCCAGGCATCAAATTAAGCA
GAAGGCCATCCTGACGGATGGCCTTTTTGCGTTTCTACAAACTCTCTGATCCTTCAACTCAGCAAAAGTTCGATTTATTCAA
CAAAGCCACGTTGTGTCTCAAAATCTCTGATGTTACATTGCACAAGATAAAAATATATCATCATGAACAATAAAACTGTCTG
CTTACATAAACAGTAATACAAGGGGTGTTATGAGCCATATTCAACGGGAAACGTCTTGCTCTAGGCCGCGATTAAATTCCAA
CATGGATGCTGATTTATATGGGTATAAATGGGCTCGCGATAATGTCGGGCAATCAGGTGCGACAATCTATCGATTGTATGGG
AAGCCCGATGCGCCAGAGTTGTTTCTGAAACATGGCAAAGGTAGCGTTGCCAATGATGTTACAGATGAGATGGTCAGACTAA
ACTGGCTGACGGAATTTATGCCTCTGCCGACCATCAAGCATTTTATCCGTACTCCTGATGATGCATGGTTACTCACCACGGC
GATCCCAGGGAAAACAGCATTCCAGGTATTAGAAGAATATCCTGATTCAGGTGAAAATATTGTTGATGCGCTGGCGGTGTTC
CTGCGCCGGTTGCATTCGATTCCTGTTTGTAATTGTCCTTTTAACAGCGACCGCGTATTTCGTCTGGCTCAGGCGCAATCAC
GAATGAATAACGGTTTGGTTGATGCGAGTGATTTTGATGACGAGCGTAATGGCTGGCCTGTTGAACAAGTCTGGAAAGAAAT
GCATAAACTTTTGCCATTCTCACCGGATTCAGTCGTCACTCATGGTGATTTCTCACTTGATAACCTTATTTTTGACGAGGGG
AAATTAATAGGTTGTATTGATGTTGGACGAGTCGGAATCGCAGACCGATACCAGGATCTTGCCATCCTATGGAACTGCCTCG
GTGAGTTTTCTCCTTCATTACAGAAACGGCTTTTTCAAAAATATGGTATTGATAATCCTGATATGAATAAATTGCAGTTTCA
TTTGATGCTCGATGAGTTTTTCTAACTGTCAGACCAAGTTTACTCATATATACTTTAGATTGATTT**GAAGAC**TACGCGCCCT
GTAGCGGCGCATTAAGCGCGGCGGGTGTGGTGGTTACGCGCAGCGTGACCGCTACACTTGCCAGCGCCCTAGCGCCCGCTCC
TTTCGCTTTCTTCCCTTCCTTTCTCGCCACGTTCGCCGGCTTTCCCCGTCAAGCTCTAAATCGGGGGCTCCCTTTAGGGTTC
CGATTTAGTGCTTTACGGCACCTCGACCCCAAAAAACTTGATTTGGGTGATGGTTCACGTAGTGGGCCATCGCCCTGATAGA
CGGTTTTTCGCCCTTTGACGTTGGAGTCCACGTTCTTTAATAGTGGACTCTTGTTCCAAACTGGAACAACACTCAACCCTAT
CTCGGGCTATTCTTTTGATTTATAAGGGATTTTGCCGATTTCGGCCTATTGGTTAAAAAATGAGCTGATTTAACAAAAATTT
AACGCGAATTTTAACAAAATATTAACGTTTACAATTTAAAAGGATCTAGGTGAAGATCCTTTTTGATAATCTCATGACCAAA
ATCCCTTAACGTGAGTTTTCGTTCCACTGAGCGTCAGACCCCGTAGAAAAGATCAAAGGATCTTCTTGAGATCCTTTTTTTC
TGCGCGTAATCTGCTGCTTGCAAACAAAAAAACCACCGCTACCAGCGGTGGTTTGTTTGCCGGATCAAGAGCTACCAACTCT
TTTTCCGAAGGTAACTGGCTTCAGCAGAGCGCAGATACCAAATACTGTCCTTCTAGTGTAGCCGTAGTTAGGCCACCACTTC
AAGAACTCTGTAGCACCGCCTACATACCTCGCTCTGCTAATCCTGTTACCAGTGGCTGCTGCCAGTGGCGATAAGTCGTGTC
TTACCGGGTTGGACTCAAGACGATAGTTACCGGATAAGGCGCAGCGGTCGGGCTGAACGGGGGGTTCGTGCACACAGCCCAG
CTTGGAGCGAACGACCTACACCGAACTGAGATACCTACAGCGTGAGCTATGAGAAAGCGCCACGCTTCCCGAAGGGAGAAAG
GCGGACAGGTATCCGGTAAGCGGCAGGGTCGGAACAGGAGAGCGCACGAGGGAGCTTCCAGGGGGAAACGCCTGGTATCTTT
ATAGTCCTGTCGGGTTTCGCCACCTCTGACTTGAGCGTCGATTTTTGTGATGCTCGTCAGGGGGGCGGAGCCTATGGAAAAA
CGCCAGCAACGCGGCCTTTTTACGGTTCCTGGCCTTTTGCTGGCCTTTTGCTCACATGTTCTTTCCTGCGTTATCCCCTGAT
TCTGTGGATAACCGTATTACCGCCTTTGAGTGAGCTGATACCGCTCGCCGCAGCCGAACGACCGAGCGCAGCGAGTCAGTGA
GCGAGGAAGCGGTAGAGCGCCTGATGCGGTATTTTCTCCTTACGCATCTGTGCGGTATTTCACACCGCATAGGGTCATGGCT
GCGCCCCGACACCCGCCAACACCCGCTGACGCGCCCTGACGGGCTTGTCTGCTCCCGGCATCCGCTTACAGACAAGCTGTGA
CCGTGTCCGGGAGCTGCATGTGTCAGAGGTTTTCACCGTCATCACCGAAACGCGCGAGGCAGAAGGAGATGGCGCCCAACAG
TCCCCCGGCCACGGGGCCTGCCACCATACCCACGCCGAAACAAGCGCTCATGAGCCCGAAGTGGCGAGCCCGATCTTCCCCA
TCGGTGATGTCGGCGATATAGGCGCCAGCAACCGCACCTGTGGCGCCGGTGATGCCGGCCACGATGCGTCCGGCGTAGAGGA
TCTGCTCATGTTTGACAGCTTATCATCGATTCAGCTTTTCAGCCGCCGCCAGAACGTCGTCCGGCTGATGCCTAAATAATTC

GCCGCTGCTGTTTTATCGCCATTAAATTTCTCCAGTGCCTGTTGTGGTGTCAGTAAGCGTGGAGCGGGAGTTTTCGCCGACT
CGCGCGCCAGTTCCGGCAGTAGCAGCTGCAAAAATTGCGGCGTTAAATCCGGCGTCGGTTCCACACTTAAAAATAGCGCCAG
TCGCTCCATCATATTGCGCAGTTCACGAATATTGCCCGGCCAGTCGTAGTGCACCAGCACGGTTTCGCTTGCCTGTAATCCC
TGGCGTAATGCGGCAGAAAACGGGGTGGAGAGCGCCGCCAGAGACACTTTCAAAAAGCTTTCCGCCAGTGGCAGAATATCCG
CCACCCGCTCGCGCAGCGGTGGCAATTGCAGACGCAAAATACTCAGCCGATAAAACAGGTCACGGCGAAACTGCCCTTGCCG
CATATCTTCTTCCAGATTACAGTGAGTGGCGCTAATGACCCGCACATCCACCGGAACAGGCTGATGCCCGCCGACGCGGGTG
ACCTCTTTTTCTTCCAGCACCCGCAGCAGCCGGGTCTGCAACGGCAGCGGCATTTCGCCAATCTCATCGAGAAACAGCGTAC
CTCCGTGGGCGATTTCAAACAGCCCGGCGCGACCGCCGCGTCGCGAGCCGGTAAACGCCCCTTCCTCATAGCCAAACAGTTC
TGCTTCCAGCAGCGATTCGGCAATCGCCCCGCAGTTGACTGCAACAAACGGATGCGACTTTTTGCCCTGTCGCGCATCGTGG
CGGGCAAAATATTCCCGATGAATCGCCTGGGCCGCCAGCTCTTTGCCCGTCCCCGTTTCCCCCTCAATCAACACCGCTGCAC
TGGAGCGGGCATACAGCAAAATAGTCTGCCGTACTTGTTCCATCTGTGGTGATTGACCGAGCATATCGCCCAGCACGTAACG
AGTTCTCAGGGCGTTGCGGGTGGCATCGTGAGTGTTATGGCGTAACGACATGCGCGTCATATCCAGCGCATCGCTGAACGCC
TGGCGCACGGTGGCGGCGGAATAGATAAAAATTCCGGTCATTCCGGCTTCTTCTGCCAGATCGGTAATCAGCCCCGCGCCGA
CCACCGCTTCGGTGCCGTTAGCTTTTAGCTCGTTAATCTGCCCGCGTGCATCTTCTTCGGTAATGTAGCTACGTTGATCGAG
GCGCAAATTAAAGGTTTTTTGAAACGCCACCAGTGCCGGAATAGTTTCCTGATAAGTGACAACGCCGATCGAGCTGGTGAGT
TTTCCGGCTTTTGCCAGTGCCTGTAACACATCGTAGCCGCTCGGTTTAATCAAAATAACTGGCACTGACAGGCGGCTTTTCA
GGTACGCGCCGTTAGATCCAGCGGCGATGATGGCGTCACAGCGTTCGTTTGCCAGTTTCTTGTGGATGTAGGTCACCGCTTT
TTCAAAGCCAAGCTGGATAGGGGTAATGTTCGCCAGGTGATCAAACTCGAGGCTGATATCGCGAAACAGCTCGAACAGGCGC
GTTACAGATACCGTCCAGATAACCGGTTTGTCGTCATTAAGCCGTGGTGGATGTGCCATAGCGCACCGCAAAGTTAAGAAAC
C

#### *Adaptor 3* ###

**pX-TU(III') |** SapI, GG III' site (*BsmBI*, BsmBI), BtsI, KanR, **BbsI**, ccdB

GAATATTGGGTTTAGTCTTGTTTCATAATTGTTGCAATGAAACGCGGTGAAACATTGCCTGAAACGTTAACTGAAACGCATA
TTTGCGGATTAGTTCATGACTTTATCTCTAACAAATTGAAATTAAACATTTAATTTTATTAAGGCAATTGTGGCACACCCCT
TGCTTTGTCTTTATCAACGCAAATAACAAGTTGATAACAAGCTAGCAGGAGGAATTCCATATGGGCTCTTCAGGGTAATCGC
T*GAGACG*TGGGGCACTGCTGAACTAGTCTGATACAGTCGACCTGCAGGCATGCAAGCTTGGCTGTTTTGGCGGATGAGAGAA
GATTTTCAGCCTGATACAGATTAAATCAGAACGCAGAAGCGGTCTGATAAAACAGAATTTGCCTGGCGGCAGTAGCGCGGTG
GTCCCACCTGACCCCATGCCGAACTCAGAAGTGAAACGCCGTAGCGCCGATGGTAGTGTGGCCAGAGCCCATGCGAGAGTAG
GGAACTGCCAGGCATCAAATAAAACGAAAGGCTCAGTCGAAAGACTGGGCCTTTCGTTTTATCTGTTGTTTGTCGGTGAACG
CTCTCCTGAGTAGGACAAATCCGCCGGGAGCGGATTTGAACGTTGCGAAGCAACGGCCCGGAGGGTGGCGGGCAGGACGCCC
GCCATAAACTGCCAGGCATCAAATTAAGCAGAAGGCCATCCTGACGGATGGCCTTTTTGCGTTTCTACAAACTCTCTGATCC
TTCAACTCAGCAAAAGTTCGATTTATTCAACAAAGCCACGTTGTGTCTCAAAATCTCTGATGTTACATTGCACAAGATAAAA
ATATATCATCATGAACAATAAAACTGTCTGCTTACATAAACAGTAATACAAGGGGTGTTATGAGCCATATTCAACGGGAAAC
GTCTTGCTCTAGGCCGCGATTAAATTCCAACATGGATGCTGATTTATATGGGTATAAATGGGCTCGCGATAATGTCGGGCAA
TCAGGTGCGACAATCTATCGATTGTATGGGAAGCCCGATGCGCCAGAGTTGTTTCTGAAACATGGCAAAGGTAGCGTTGCCA
ATGATGTTACAGATGAGATGGTCAGACTAAACTGGCTGACGGAATTTATGCCTCTGCCGACCATCAAGCATTTTATCCGTAC
TCCTGATGATGCATGGTTACTCACCACGGCGATCCCAGGGAAAACAGCATTCCAGGTATTAGAAGAATATCCTGATTCAGGT
GAAAATATTGTTGATGCGCTGGCGGTGTTCCTGCGCCGGTTGCATTCGATTCCTGTTTGTAATTGTCCTTTTAACAGCGACC
GCGTATTTCGTCTGGCTCAGGCGCAATCACGAATGAATAACGGTTTGGTTGATGCGAGTGATTTTGATGACGAGCGTAATGG
CTGGCCTGTTGAACAAGTCTGGAAAGAAATGCATAAACTTTTGCCATTCTCACCGGATTCAGTCGTCACTCATGGTGATTTC
TCACTTGATAACCTTATTTTTGACGAGGGGAAATTAATAGGTTGTATTGATGTTGGACGAGTCGGAATCGCAGACCGATACC
AGGATCTTGCCATCCTATGGAACTGCCTCGGTGAGTTTTCTCCTTCATTACAGAAACGGCTTTTTCAAAAATATGGTATTGA
TAATCCTGATATGAATAAATTGCAGTTTCATTTGATGCTCGATGAGTTTTTCTAACTGTCAGACCAAGTTTACTCATATATA
CTTTAGATTGATTT**GAAGAC**TACGCGCCCTGTAGCGGCGCATTAAGCGCGGCGGGTGTGGTGGTTACGCGCAGCGTGACCGC
TACACTTGCCAGCGCCCTAGCGCCCGCTCCTTTCGCTTTCTTCCCTTCCTTTCTCGCCACGTTCGCCGGCTTTCCCCGTCAA
GCTCTAAATCGGGGGCTCCCTTTAGGGTTCCGATTTAGTGCTTTACGGCACCTCGACCCCAAAAAACTTGATTTGGGTGATG
GTTCACGTAGTGGGCCATCGCCCTGATAGACGGTTTTTCGCCCTTTGACGTTGGAGTCCACGTTCTTTAATAGTGGACTCTT
GTTCCAAACTGGAACAACACTCAACCCTATCTCGGGCTATTCTTTTGATTTATAAGGGATTTTGCCGATTTCGGCCTATTGG
TTAAAAAATGAGCTGATTTAACAAAAATTTAACGCGAATTTTAACAAAATATTAACGTTTACAATTTAAAAGGATCTAGGTG
AAGATCCTTTTTGATAATCTCATGACCAAAATCCCTTAACGTGAGTTTTCGTTCCACTGAGCGTCAGACCCCGTAGAAAAGA
TCAAAGGATCTTCTTGAGATCCTTTTTTTCTGCGCGTAATCTGCTGCTTGCAAACAAAAAAACCACCGCTACCAGCGGTGGT
TTGTTTGCCGGATCAAGAGCTACCAACTCTTTTTCCGAAGGTAACTGGCTTCAGCAGAGCGCAGATACCAAATACTGTCCTT
CTAGTGTAGCCGTAGTTAGGCCACCACTTCAAGAACTCTGTAGCACCGCCTACATACCTCGCTCTGCTAATCCTGTTACCAG
TGGCTGCTGCCAGTGGCGATAAGTCGTGTCTTACCGGGTTGGACTCAAGACGATAGTTACCGGATAAGGCGCAGCGGTCGGG
CTGAACGGGGGGTTCGTGCACACAGCCCAGCTTGGAGCGAACGACCTACACCGAACTGAGATACCTACAGCGTGAGCTATGA
GAAAGCGCCACGCTTCCCGAAGGGAGAAAGGCGGACAGGTATCCGGTAAGCGGCAGGGTCGGAACAGGAGAGCGCACGAGGG
AGCTTCCAGGGGGAAACGCCTGGTATCTTTATAGTCCTGTCGGGTTTCGCCACCTCTGACTTGAGCGTCGATTTTTGTGATG
CTCGTCAGGGGGGCGGAGCCTATGGAAAAACGCCAGCAACGCGGCCTTTTTACGGTTCCTGGCCTTTTGCTGGCCTTTTGCT
CACATGTTCTTTCCTGCGTTATCCCCTGATTCTGTGGATAACCGTATTACCGCCTTTGAGTGAGCTGATACCGCTCGCCGCA
GCCGAACGACCGAGCGCAGCGAGTCAGTGAGCGAGGAAGCGGTAGAGCGCCTGATGCGGTATTTTCTCCTTACGCATCTGTG
CGGTATTTCACACCGCATAGGGTCATGGCTGCGCCCCGACACCCGCCAACACCCGCTGACGCGCCCTGACGGGCTTGTCTGC
TCCCGGCATCCGCTTACAGACAAGCTGTGACCGTGTCCGGGAGCTGCATGTGTCAGAGGTTTTCACCGTCATCACCGAAACG
CGCGAGGCAGAAGGAGATGGCGCCCAACAGTCCCCCGGCCACGGGGCCTGCCACCATACCCACGCCGAAACAAGCGCTCATG
AGCCCGAAGTGGCGAGCCCGATCTTCCCCATCGGTGATGTCGGCGATATAGGCGCCAGCAACCGCACCTGTGGCGCCGGTGA
TGCCGGCCACGATGCGTCCGGCGTAGAGGATCTGCTCATGTTTGACAGCTTATCATCGATTTATATTCCCCAGAACATCAGG
TTAATGGCGTTTTTGATGTCATTTTCGCGGTGGCTGAGATCAGCCACTTCTTCCCCGATAACGGAGACTGGCACACTGGCCA
TATCGGTGGTCATCATGCGCCAGCTTTCATCCCCGATATGCACCACCGGGTAAAGTTCACGGAGACTTTATCTGACAGCAG
ACGTGCACTGGCCAGGGGGATCACCATCCGTCGCCCGGGCGTGTCAATAATATCACTCTGTACATCCACAAACAGACGATAA
CGGCTCTCTCTTTTATAGGTGTAAACCTTAAACTGCATAGCGCACCGCAAAGTTAAGAAACC

**pZ-MBP-TU(III)-sRBS |** MBP, GG III site (*BsmBI*, BsmBI), sRBS, BtsI, KanR, **BbsI**

GAATATTGGGTTTAGTCTTGTTTCATAATTGTTGCAATGAAACGCGGTGAAACATTGCCTGAAACGTTAACTGAAACGCATA
TTTGCGGATTAGTTCATGACTTTATCTCTAACAAATTGAAATTAAACATTTAATTTTATTAAGGCAATTGTGGCACACCCCT
TGCTTTGTCTTTATCAACGCAAATAACAAGTTGATAACAAGCTAGCAGGAGGAATTCCATATGGGGAAAATCGAAGAAGGTA
AACTGGTAATCTGGATTAACGGCGATAAAGGCTATAACGGTCTGGCTGAAGTCGGTAAGAAATTCGAGAAAGATACCGGAAT
TAAAGTCACCGTTGAGCATCCGGATAAACTGGAAGAGAAATTCCCACAGGTTGCGGCAACTGGCGATGGCCCTGACATTATC
TTCTGGGCACACGACCGCTTTGGTGGCTACGCTCAATCTGGCCTGTTGGCTGAAATCACCCCGGACAAAGCGTTCCAGGACA
AGCTGTATCCGTTTACCTGGGATGCCGTACGTTACAACGGCAAGCTGATTGCTTACCCGATCGCTGTTGAAGCGTTATCGCT
GATTTATAACAAAGATCTGCTGCCGAACCCGCCAAAAACCTGGGAAGAGATCCCGGCGCTGGATAAAGAACTGAAAGCGAAA
GGTAAGAGCGCGCTGATGTTCAACCTGCAAGAACCGTACTTCACCTGGCCGCTGATTGCTGCTGACGGGGGTTATGCGTTCA
AGTATGAAAACGGCAAGTACGACATTAAAGACGTGGGCGTGGATAACGCTGGCGCGAAAGCGGGTCTGACCTTCCTGGTTGA
CCTGATTAAAAACAAACACATGAATGCAGACACCGATTACTCCATCGCAGAAGCTGCCTTTAATAAAGGCGAAACAGCGATG
ACCATCAACGGCCCGTGGGCATGGTCCAACATCGACACCAGCAAAGTGAATTATGGTGTAACGGTACTGCCGACCTTCAAGG
GTCAACCATCCAAACCGTTCGTTGGCGTGCTGAGCGCAGGTATTAACGCCGCCAGTCCGAACAAAGAGCTGGCAAAAGAGTT
CCTCGAAAACTATCTGCTGACTGATGAAGGTCTGGAAGCGGTTAATAAAGACAAACCGCTGGGTGCCGTAGCGCTGAAGTCT
TACGAGGAAGAGTTGGTGAAAGATCCGCGTATTGCCGCCACTATGGAAAACGCCCAGAAAGGTGAAATCATGCCGAACATCC
CGCAGATGTCCGCTTTCTGGTATGCCGTGCGTACTGCGGTGATCAACGCCGCCAGCGGTCGTCAGACTGTCGATGAAGCCCT
GAAAGACGCGCAGACTGGGTAA*CGTCTCA*TCGCATTTAAAAAGAGGAGAAAGTACATTATGGGGCACTGCTGAACTAGTCTG
ATACAGTCGACCTGCAGGCATGCAAGCTTGGCTGTTTTGGCGGATGAGAGAAGATTTTCAGCCTGATACAGATTAAATCAGA
ACGCAGAAGCGGTCTGATAAAACAGAATTTGCCTGGCGGCAGTAGCGCGGTGGTCCCACCTGACCCCATGCCGAACTCAGAA
GTGAAACGCCGTAGCGCCGATGGTAGTGTGGCCAGAGCCCATGCGAGAGTAGGGAACTGCCAGGCATCAAATAAAACGAAAG
GCTCAGTCGAAAGACTGGGCCTTTCGTTTTATCTGTTGTTTGTCGGTGAACGCTCTCCTGAGTAGGACAAATCCGCCGGGAG
CGGATTTGAACGTTGCGAAGCAACGGCCCGGAGGGTGGCGGGCAGGACGCCCGCCATAAACTGCCAGGCATCAAATTAAGCA
GAAGGCCATCCTGACGGATGGCCTTTTTGCGTTTCTACAAACTCTCTGATCCTTCAACTCAGCAAAAGTTCGATTTATTCAA
CAAAGCCACGTTGTGTCTCAAAATCTCTGATGTTACATTGCACAAGATAAAAATATATCATCATGAACAATAAAACTGTCTG
CTTACATAAACAGTAATACAAGGGGTGTTATGAGCCATATTCAACGGGAAACGTCTTGCTCTAGGCCGCGATTAAATTCCAA
CATGGATGCTGATTTATATGGGTATAAATGGGCTCGCGATAATGTCGGGCAATCAGGTGCGACAATCTATCGATTGTATGGG
AAGCCCGATGCGCCAGAGTTGTTTCTGAAACATGGCAAAGGTAGCGTTGCCAATGATGTTACAGATGAGATGGTCAGACTAA
ACTGGCTGACGGAATTTATGCCTCTGCCGACCATCAAGCATTTTATCCGTACTCCTGATGATGCATGGTTACTCACCACGGC
GATCCCAGGGAAAACAGCATTCCAGGTATTAGAAGAATATCCTGATTCAGGTGAAAATATTGTTGATGCGCTGGCGGTGTTC
CTGCGCCGGTTGCATTCGATTCCTGTTTGTAATTGTCCTTTTAACAGCGACCGCGTATTTCGTCTGGCTCAGGCGCAATCAC
GAATGAATAACGGTTTGGTTGATGCGAGTGATTTTGATGACGAGCGTAATGGCTGGCCTGTTGAACAAGTCTGGAAAGAAAT
GCATAAACTTTTGCCATTCTCACCGGATTCAGTCGTCACTCATGGTGATTTCTCACTTGATAACCTTATTTTTGACGAGGGG
AAATTAATAGGTTGTATTGATGTTGGACGAGTCGGAATCGCAGACCGATACCAGGATCTTGCCATCCTATGGAACTGCCTCG
GTGAGTTTTCTCCTTCATTACAGAAACGGCTTTTTCAAAAATATGGTATTGATAATCCTGATATGAATAAATTGCAGTTTCA
TTTGATGCTCGATGAGTTTTTCTAACTGTCAGACCAAGTTTACTCATATATACTTTAGATTGATTT**GAAGAC**TACGCGCCCT
GTAGCGGCGCATTAAGCGCGGCGGGTGTGGTGGTTACGCGCAGCGTGACCGCTACACTTGCCAGCGCCCTAGCGCCCGCTCC
TTTCGCTTTCTTCCCTTCCTTTCTCGCCACGTTCGCCGGCTTTCCCCGTCAAGCTCTAAATCGGGGGCTCCCTTTAGGGTTC
CGATTTAGTGCTTTACGGCACCTCGACCCCAAAAAACTTGATTTGGGTGATGGTTCACGTAGTGGGCCATCGCCCTGATAGA
CGGTTTTTCGCCCTTTGACGTTGGAGTCCACGTTCTTTAATAGTGGACTCTTGTTCCAAACTGGAACAACACTCAACCCTAT
CTCGGGCTATTCTTTTGATTTATAAGGGATTTTGCCGATTTCGGCCTATTGGTTAAAAAATGAGCTGATTTAACAAAAATTT
AACGCGAATTTTAACAAAATATTAACGTTTACAATTTAAAAGGATCTAGGTGAAGATCCTTTTTGATAATCTCATGACCAAA
ATCCCTTAACGTGAGTTTTCGTTCCACTGAGCGTCAGACCCCGTAGAAAAGATCAAAGGATCTTCTTGAGATCCTTTTTTTC
TGCGCGTAATCTGCTGCTTGCAAACAAAAAAACCACCGCTACCAGCGGTGGTTTGTTTGCCGGATCAAGAGCTACCAACTCT
TTTTCCGAAGGTAACTGGCTTCAGCAGAGCGCAGATACCAAATACTGTCCTTCTAGTGTAGCCGTAGTTAGGCCACCACTTC
AAGAACTCTGTAGCACCGCCTACATACCTCGCTCTGCTAATCCTGTTACCAGTGGCTGCTGCCAGTGGCGATAAGTCGTGTC
TTACCGGGTTGGACTCAAGACGATAGTTACCGGATAAGGCGCAGCGGTCGGGCTGAACGGGGGGTTCGTGCACACAGCCCAG
CTTGGAGCGAACGACCTACACCGAACTGAGATACCTACAGCGTGAGCTATGAGAAAGCGCCACGCTTCCCGAAGGGAGAAAG
GCGGACAGGTATCCGGTAAGCGGCAGGGTCGGAACAGGAGAGCGCACGAGGGAGCTTCCAGGGGGAAACGCCTGGTATCTTT
ATAGTCCTGTCGGGTTTCGCCACCTCTGACTTGAGCGTCGATTTTTGTGATGCTCGTCAGGGGGGCGGAGCCTATGGAAAAA
CGCCAGCAACGCGGCCTTTTTACGGTTCCTGGCCTTTTGCTGGCCTTTTGCTCACATGTTCTTTCCTGCGTTATCCCCTGAT
TCTGTGGATAACCGTATTACCGCCTTTGAGTGAGCTGATACCGCTCGCCGCAGCCGAACGACCGAGCGCAGCGAGTCAGTGA
GCGAGGAAGCGGTAGAGCGCCTGATGCGGTATTTTCTCCTTACGCATCTGTGCGGTATTTCACACCGCATAGGGTCATGGCT
GCGCCCCGACACCCGCCAACACCCGCTGACGCGCCCTGACGGGCTTGTCTGCTCCCGGCATCCGCTTACAGACAAGCTGTGA
CCGTGTCCGGGAGCTGCATGTGTCAGAGGTTTTCACCGTCATCACCGAAACGCGCGAGGCAGAAGGAGATGGCGCCCAACAG
TCCCCCGGCCACGGGGCCTGCCACCATACCCACGCCGAAACAAGCGCTCATGAGCCCGAAGTGGCGAGCCCGATCTTCCCCA
TCGGTGATGTCGGCGATATAGGCGCCAGCAACCGCACCTGTGGCGCCGGTGATGCCGGCCACGATGCGTCCGGCGTAGAGGA
TCTGCTCATGTTTGACAGCTTATCATCGATTCAGCTTTTCAGCCGCCGCCAGAACGTCGTCCGGCTGATGCCTAAATAATTC

GCCGCTGCTGTTTTATCGCCATTAAATTTCTCCAGTGCCTGTTGTGGTGTCAGTAAGCGTGGAGCGGGAGTTTTCGCCGACT
CGCGCGCCAGTTCCGGCAGTAGCAGCTGCAAAAATTGCGGCGTTAAATCCGGCGTCGGTTCCACACTTAAAAATAGCGCCAG
TCGCTCCATCATATTGCGCAGTTCACGAATATTGCCCGGCCAGTCGTAGTGCACCAGCACGGTTTCGCTTGCCTGTAATCCC
TGGCGTAATGCGGCAGAAAACGGGGTGGAGAGCGCCGCCAGAGACACTTTCAAAAAGCTTTCCGCCAGTGGCAGAATATCCG
CCACCCGCTCGCGCAGCGGTGGCAATTGCAGACGCAAAATACTCAGCCGATAAAACAGGTCACGGCGAAACTGCCCTTGCCG
CATATCTTCTTCCAGATTACAGTGAGTGGCGCTAATGACCCGCACATCCACCGGAACAGGCTGATGCCCGCCGACGCGGGTG
ACCTCTTTTTCTTCCAGCACCCGCAGCAGCCGGGTCTGCAACGGCAGCGGCATTTCGCCAATCTCATCGAGAAACAGCGTAC
CTCCGTGGGCGATTTCAAACAGCCCGGCGCGACCGCCGCGTCGCGAGCCGGTAAACGCCCCTTCCTCATAGCCAAACAGTTC
TGCTTCCAGCAGCGATTCGGCAATCGCCCCGCAGTTGACTGCAACAAACGGATGCGACTTTTTGCCCTGTCGCGCATCGTGG
CGGGCAAAATATTCCCGATGAATCGCCTGGGCCGCCAGCTCTTTGCCCGTCCCCGTTTCCCCCTCAATCAACACCGCTGCAC
TGGAGCGGGCATACAGCAAAATAGTCTGCCGTACTTGTTCCATCTGTGGTGATTGACCGAGCATATCGCCCAGCACGTAACG
AGTTCTCAGGGCGTTGCGGGTGGCATCGTGAGTGTTATGGCGTAACGACATGCGCGTCATATCCAGCGCATCGCTGAACGCC
TGGCGCACGGTGGCGGCGGAATAGATAAAAATTCCGGTCATTCCGGCTTCTTCTGCCAGATCGGTAATCAGCCCCGCGCCGA
CCACCGCTTCGGTGCCGTTAGCTTTTAGCTCGTTAATCTGCCCGCGTGCATCTTCTTCGGTAATGTAGCTACGTTGATCGAG
GCGCAAATTAAAGGTTTTTTGAAACGCCACCAGTGCCGGAATAGTTTCCTGATAAGTGACAACGCCGATCGAGCTGGTGAGT
TTTCCGGCTTTTGCCAGTGCCTGTAACACATCGTAGCCGCTCGGTTTAATCAAAATAACTGGCACTGACAGGCGGCTTTTCA
GGTACGCGCCGTTAGATCCAGCGGCGATGATGGCGTCACAGCGTTCGTTTGCCAGTTTCTTGTGGATGTAGGTCACCGCTTT
TTCAAAGCCAAGCTGGATAGGGGTAATGTTCGCCAGGTGATCAAACTCGAGGCTGATATCGCGAAACAGCTCGAACAGGCGC
GTTACAGATACCGTCCAGATAACCGGTTTGTCGTCATTAAGCCGTGGTGGATGTGCCATAGCGCACCGCAAAGTTAAGAAAC
C

#### *Adaptor 4* ###

**pZ-MBP-TU(IV)-sRBS | **MBP, GG IV site (*BsmBI*, BsmBI), sRBS, BtsI, KanR, **BbsI**

GAATATTGGGTTTAGTCTTGTTTCATAATTGTTGCAATGAAACGCGGTGAAACATTGCCTGAAACGTTAACTGAAACGCATA
TTTGCGGATTAGTTCATGACTTTATCTCTAACAAATTGAAATTAAACATTTAATTTTATTAAGGCAATTGTGGCACACCCCT
TGCTTTGTCTTTATCAACGCAAATAACAAGTTGATAACAAGCTAGCAGGAGGAATTCCATATGGGGAAAATCGAAGAAGGTA
AACTGGTAATCTGGATTAACGGCGATAAAGGCTATAACGGTCTGGCTGAAGTCGGTAAGAAATTCGAGAAAGATACCGGAAT
TAAAGTCACCGTTGAGCATCCGGATAAACTGGAAGAGAAATTCCCACAGGTTGCGGCAACTGGCGATGGCCCTGACATTATC
TTCTGGGCACACGACCGCTTTGGTGGCTACGCTCAATCTGGCCTGTTGGCTGAAATCACCCCGGACAAAGCGTTCCAGGACA
AGCTGTATCCGTTTACCTGGGATGCCGTACGTTACAACGGCAAGCTGATTGCTTACCCGATCGCTGTTGAAGCGTTATCGCT
GATTTATAACAAAGATCTGCTGCCGAACCCGCCAAAAACCTGGGAAGAGATCCCGGCGCTGGATAAAGAACTGAAAGCGAAA
GGTAAGAGCGCGCTGATGTTCAACCTGCAAGAACCGTACTTCACCTGGCCGCTGATTGCTGCTGACGGGGGTTATGCGTTCA
AGTATGAAAACGGCAAGTACGACATTAAAGACGTGGGCGTGGATAACGCTGGCGCGAAAGCGGGTCTGACCTTCCTGGTTGA
CCTGATTAAAAACAAACACATGAATGCAGACACCGATTACTCCATCGCAGAAGCTGCCTTTAATAAAGGCGAAACAGCGATG
ACCATCAACGGCCCGTGGGCATGGTCCAACATCGACACCAGCAAAGTGAATTATGGTGTAACGGTACTGCCGACCTTCAAGG
GTCAACCATCCAAACCGTTCGTTGGCGTGCTGAGCGCAGGTATTAACGCCGCCAGTCCGAACAAAGAGCTGGCAAAAGAGTT
CCTCGAAAACTATCTGCTGACTGATGAAGGTCTGGAAGCGGTTAATAAAGACAAACCGCTGGGTGCCGTAGCGCTGAAGTCT
TACGAGGAAGAGTTGGTGAAAGATCCGCGTATTGCCGCCACTATGGAAAACGCCCAGAAAGGTGAAATCATGCCGAACATCC
CGCAGATGTCCGCTTTCTGGTATGCCGTGCGTACTGCGGTGATCAACGCCGCCAGCGGTCGTCAGACTGTCGATGAAGCCCT
GAAAGACGCGCAGACTGGGTAA*CGTCTC*ACGAAATTTAAAAAGAGGAGAAAGTACATTATGGGGCACTGCTGAACTAGTCTG
ATACAGTCGACCTGCAGGCATGCAAGCTTGGCTGTTTTGGCGGATGAGAGAAGATTTTCAGCCTGATACAGATTAAATCAGA
ACGCAGAAGCGGTCTGATAAAACAGAATTTGCCTGGCGGCAGTAGCGCGGTGGTCCCACCTGACCCCATGCCGAACTCAGAA
GTGAAACGCCGTAGCGCCGATGGTAGTGTGGCCAGAGCCCATGCGAGAGTAGGGAACTGCCAGGCATCAAATAAAACGAAAG
GCTCAGTCGAAAGACTGGGCCTTTCGTTTTATCTGTTGTTTGTCGGTGAACGCTCTCCTGAGTAGGACAAATCCGCCGGGAG
CGGATTTGAACGTTGCGAAGCAACGGCCCGGAGGGTGGCGGGCAGGACGCCCGCCATAAACTGCCAGGCATCAAATTAAGCA
GAAGGCCATCCTGACGGATGGCCTTTTTGCGTTTCTACAAACTCTCTGATCCTTCAACTCAGCAAAAGTTCGATTTATTCAA
CAAAGCCACGTTGTGTCTCAAAATCTCTGATGTTACATTGCACAAGATAAAAATATATCATCATGAACAATAAAACTGTCTG
CTTACATAAACAGTAATACAAGGGGTGTTATGAGCCATATTCAACGGGAAACGTCTTGCTCTAGGCCGCGATTAAATTCCAA
CATGGATGCTGATTTATATGGGTATAAATGGGCTCGCGATAATGTCGGGCAATCAGGTGCGACAATCTATCGATTGTATGGG
AAGCCCGATGCGCCAGAGTTGTTTCTGAAACATGGCAAAGGTAGCGTTGCCAATGATGTTACAGATGAGATGGTCAGACTAA
ACTGGCTGACGGAATTTATGCCTCTGCCGACCATCAAGCATTTTATCCGTACTCCTGATGATGCATGGTTACTCACCACGGC
GATCCCAGGGAAAACAGCATTCCAGGTATTAGAAGAATATCCTGATTCAGGTGAAAATATTGTTGATGCGCTGGCGGTGTTC
CTGCGCCGGTTGCATTCGATTCCTGTTTGTAATTGTCCTTTTAACAGCGACCGCGTATTTCGTCTGGCTCAGGCGCAATCAC
GAATGAATAACGGTTTGGTTGATGCGAGTGATTTTGATGACGAGCGTAATGGCTGGCCTGTTGAACAAGTCTGGAAAGAAAT
GCATAAACTTTTGCCATTCTCACCGGATTCAGTCGTCACTCATGGTGATTTCTCACTTGATAACCTTATTTTTGACGAGGGG
AAATTAATAGGTTGTATTGATGTTGGACGAGTCGGAATCGCAGACCGATACCAGGATCTTGCCATCCTATGGAACTGCCTCG
GTGAGTTTTCTCCTTCATTACAGAAACGGCTTTTTCAAAAATATGGTATTGATAATCCTGATATGAATAAATTGCAGTTTCA
TTTGATGCTCGATGAGTTTTTCTAACTGTCAGACCAAGTTTACTCATATATACTTTAGATTGATTT**GAAGAC**TACGCGCCCT
GTAGCGGCGCATTAAGCGCGGCGGGTGTGGTGGTTACGCGCAGCGTGACCGCTACACTTGCCAGCGCCCTAGCGCCCGCTCC
TTTCGCTTTCTTCCCTTCCTTTCTCGCCACGTTCGCCGGCTTTCCCCGTCAAGCTCTAAATCGGGGGCTCCCTTTAGGGTTC
CGATTTAGTGCTTTACGGCACCTCGACCCCAAAAAACTTGATTTGGGTGATGGTTCACGTAGTGGGCCATCGCCCTGATAGA
CGGTTTTTCGCCCTTTGACGTTGGAGTCCACGTTCTTTAATAGTGGACTCTTGTTCCAAACTGGAACAACACTCAACCCTAT
CTCGGGCTATTCTTTTGATTTATAAGGGATTTTGCCGATTTCGGCCTATTGGTTAAAAAATGAGCTGATTTAACAAAAATTT
AACGCGAATTTTAACAAAATATTAACGTTTACAATTTAAAAGGATCTAGGTGAAGATCCTTTTTGATAATCTCATGACCAAA
ATCCCTTAACGTGAGTTTTCGTTCCACTGAGCGTCAGACCCCGTAGAAAAGATCAAAGGATCTTCTTGAGATCCTTTTTTTC
TGCGCGTAATCTGCTGCTTGCAAACAAAAAAACCACCGCTACCAGCGGTGGTTTGTTTGCCGGATCAAGAGCTACCAACTCT
TTTTCCGAAGGTAACTGGCTTCAGCAGAGCGCAGATACCAAATACTGTCCTTCTAGTGTAGCCGTAGTTAGGCCACCACTTC
AAGAACTCTGTAGCACCGCCTACATACCTCGCTCTGCTAATCCTGTTACCAGTGGCTGCTGCCAGTGGCGATAAGTCGTGTC
TTACCGGGTTGGACTCAAGACGATAGTTACCGGATAAGGCGCAGCGGTCGGGCTGAACGGGGGGTTCGTGCACACAGCCCAG
CTTGGAGCGAACGACCTACACCGAACTGAGATACCTACAGCGTGAGCTATGAGAAAGCGCCACGCTTCCCGAAGGGAGAAAG
GCGGACAGGTATCCGGTAAGCGGCAGGGTCGGAACAGGAGAGCGCACGAGGGAGCTTCCAGGGGGAAACGCCTGGTATCTTT
ATAGTCCTGTCGGGTTTCGCCACCTCTGACTTGAGCGTCGATTTTTGTGATGCTCGTCAGGGGGGCGGAGCCTATGGAAAAA
CGCCAGCAACGCGGCCTTTTTACGGTTCCTGGCCTTTTGCTGGCCTTTTGCTCACATGTTCTTTCCTGCGTTATCCCCTGAT
TCTGTGGATAACCGTATTACCGCCTTTGAGTGAGCTGATACCGCTCGCCGCAGCCGAACGACCGAGCGCAGCGAGTCAGTGA
GCGAGGAAGCGGTAGAGCGCCTGATGCGGTATTTTCTCCTTACGCATCTGTGCGGTATTTCACACCGCATAGGGTCATGGCT
GCGCCCCGACACCCGCCAACACCCGCTGACGCGCCCTGACGGGCTTGTCTGCTCCCGGCATCCGCTTACAGACAAGCTGTGA
CCGTGTCCGGGAGCTGCATGTGTCAGAGGTTTTCACCGTCATCACCGAAACGCGCGAGGCAGAAGGAGATGGCGCCCAACAG
TCCCCCGGCCACGGGGCCTGCCACCATACCCACGCCGAAACAAGCGCTCATGAGCCCGAAGTGGCGAGCCCGATCTTCCCCA

TCGGTGATGTCGGCGATATAGGCGCCAGCAACCGCACCTGTGGCGCCGGTGATGCCGGCCACGATGCGTCCGGCGTAGAGGA
TCTGCTCATGTTTGACAGCTTATCATCGATTCAGCTTTTCAGCCGCCGCCAGAACGTCGTCCGGCTGATGCCTAAATAATTC
GCCGCTGCTGTTTTATCGCCATTAAATTTCTCCAGTGCCTGTTGTGGTGTCAGTAAGCGTGGAGCGGGAGTTTTCGCCGACT
CGCGCGCCAGTTCCGGCAGTAGCAGCTGCAAAAATTGCGGCGTTAAATCCGGCGTCGGTTCCACACTTAAAAATAGCGCCAG
TCGCTCCATCATATTGCGCAGTTCACGAATATTGCCCGGCCAGTCGTAGTGCACCAGCACGGTTTCGCTTGCCTGTAATCCC
TGGCGTAATGCGGCAGAAAACGGGGTGGAGAGCGCCGCCAGAGACACTTTCAAAAAGCTTTCCGCCAGTGGCAGAATATCCG
CCACCCGCTCGCGCAGCGGTGGCAATTGCAGACGCAAAATACTCAGCCGATAAAACAGGTCACGGCGAAACTGCCCTTGCCG
CATATCTTCTTCCAGATTACAGTGAGTGGCGCTAATGACCCGCACATCCACCGGAACAGGCTGATGCCCGCCGACGCGGGTG
ACCTCTTTTTCTTCCAGCACCCGCAGCAGCCGGGTCTGCAACGGCAGCGGCATTTCGCCAATCTCATCGAGAAACAGCGTAC
CTCCGTGGGCGATTTCAAACAGCCCGGCGCGACCGCCGCGTCGCGAGCCGGTAAACGCCCCTTCCTCATAGCCAAACAGTTC
TGCTTCCAGCAGCGATTCGGCAATCGCCCCGCAGTTGACTGCAACAAACGGATGCGACTTTTTGCCCTGTCGCGCATCGTGG
CGGGCAAAATATTCCCGATGAATCGCCTGGGCCGCCAGCTCTTTGCCCGTCCCCGTTTCCCCCTCAATCAACACCGCTGCAC
TGGAGCGGGCATACAGCAAAATAGTCTGCCGTACTTGTTCCATCTGTGGTGATTGACCGAGCATATCGCCCAGCACGTAACG
AGTTCTCAGGGCGTTGCGGGTGGCATCGTGAGTGTTATGGCGTAACGACATGCGCGTCATATCCAGCGCATCGCTGAACGCC
TGGCGCACGGTGGCGGCGGAATAGATAAAAATTCCGGTCATTCCGGCTTCTTCTGCCAGATCGGTAATCAGCCCCGCGCCGA
CCACCGCTTCGGTGCCGTTAGCTTTTAGCTCGTTAATCTGCCCGCGTGCATCTTCTTCGGTAATGTAGCTACGTTGATCGAG
GCGCAAATTAAAGGTTTTTTGAAACGCCACCAGTGCCGGAATAGTTTCCTGATAAGTGACAACGCCGATCGAGCTGGTGAGT
TTTCCGGCTTTTGCCAGTGCCTGTAACACATCGTAGCCGCTCGGTTTAATCAAAATAACTGGCACTGACAGGCGGCTTTTCA
GGTACGCGCCGTTAGATCCAGCGGCGATGATGGCGTCACAGCGTTCGTTTGCCAGTTTCTTGTGGATGTAGGTCACCGCTTT
TTCAAAGCCAAGCTGGATAGGGGTAATGTTCGCCAGGTGATCAAACTCGAGGCTGATATCGCGAAACAGCTCGAACAGGCGC
GTTACAGATACCGTCCAGATAACCGGTTTGTCGTCATTAAGCCGTGGTGGATGTGCCATAGCGCACCGCAAAGTTAAGAAAC
C

**pX-TU(IV')** | SapI, GG IV' site (*BsmBI*, BsmBI), BtsI, KanR, **BbsI**, ccdB

GAATATTGGGTTTAGTCTTGTTTCATAATTGTTGCAATGAAACGCGGTGAAACATTGCCTGAAACGTTAACTGAAACGCATA
TTTGCGGATTAGTTCATGACTTTATCTCTAACAAATTGAAATTAAACATTTAATTTTATTAAGGCAATTGTGGCACACCCCT
TGCTTTGTCTTTATCAACGCAAATAACAAGTTGATAACAAGCTAGCAGGAGGAATTCCATATGGGCTCTTCAGGGTAACGAA
T*GAGACG*TGGGGCACTGCTGAACTAGTCTGATACAGTCGACCTGCAGGCATGCAAGCTTGGCTGTTTTGGCGGATGAGAGAA
GATTTTCAGCCTGATACAGATTAAATCAGAACGCAGAAGCGGTCTGATAAAACAGAATTTGCCTGGCGGCAGTAGCGCGGTG
GTCCCACCTGACCCCATGCCGAACTCAGAAGTGAAACGCCGTAGCGCCGATGGTAGTGTGGCCAGAGCCCATGCGAGAGTAG
GGAACTGCCAGGCATCAAATAAAACGAAAGGCTCAGTCGAAAGACTGGGCCTTTCGTTTTATCTGTTGTTTGTCGGTGAACG
CTCTCCTGAGTAGGACAAATCCGCCGGGAGCGGATTTGAACGTTGCGAAGCAACGGCCCGGAGGGTGGCGGGCAGGACGCCC
GCCATAAACTGCCAGGCATCAAATTAAGCAGAAGGCCATCCTGACGGATGGCCTTTTTGCGTTTCTACAAACTCTCTGATCC
TTCAACTCAGCAAAAGTTCGATTTATTCAACAAAGCCACGTTGTGTCTCAAAATCTCTGATGTTACATTGCACAAGATAAAA
ATATATCATCATGAACAATAAAACTGTCTGCTTACATAAACAGTAATACAAGGGGTGTTATGAGCCATATTCAACGGGAAAC
GTCTTGCTCTAGGCCGCGATTAAATTCCAACATGGATGCTGATTTATATGGGTATAAATGGGCTCGCGATAATGTCGGGCAA
TCAGGTGCGACAATCTATCGATTGTATGGGAAGCCCGATGCGCCAGAGTTGTTTCTGAAACATGGCAAAGGTAGCGTTGCCA
ATGATGTTACAGATGAGATGGTCAGACTAAACTGGCTGACGGAATTTATGCCTCTGCCGACCATCAAGCATTTTATCCGTAC
TCCTGATGATGCATGGTTACTCACCACGGCGATCCCAGGGAAAACAGCATTCCAGGTATTAGAAGAATATCCTGATTCAGGT
GAAAATATTGTTGATGCGCTGGCGGTGTTCCTGCGCCGGTTGCATTCGATTCCTGTTTGTAATTGTCCTTTTAACAGCGACC
GCGTATTTCGTCTGGCTCAGGCGCAATCACGAATGAATAACGGTTTGGTTGATGCGAGTGATTTTGATGACGAGCGTAATGG
CTGGCCTGTTGAACAAGTCTGGAAAGAAATGCATAAACTTTTGCCATTCTCACCGGATTCAGTCGTCACTCATGGTGATTTC
TCACTTGATAACCTTATTTTTGACGAGGGGAAATTAATAGGTTGTATTGATGTTGGACGAGTCGGAATCGCAGACCGATACC
AGGATCTTGCCATCCTATGGAACTGCCTCGGTGAGTTTTCTCCTTCATTACAGAAACGGCTTTTTCAAAAATATGGTATTGA
TAATCCTGATATGAATAAATTGCAGTTTCATTTGATGCTCGATGAGTTTTTCTAACTGTCAGACCAAGTTTACTCATATATA
CTTTAGATTGATTT**GAAGAC**TACGCGCCCTGTAGCGGCGCATTAAGCGCGGCGGGTGTGGTGGTTACGCGCAGCGTGACCGC
TACACTTGCCAGCGCCCTAGCGCCCGCTCCTTTCGCTTTCTTCCCTTCCTTTCTCGCCACGTTCGCCGGCTTTCCCCGTCAA
GCTCTAAATCGGGGGCTCCCTTTAGGGTTCCGATTTAGTGCTTTACGGCACCTCGACCCCAAAAAACTTGATTTGGGTGATG
GTTCACGTAGTGGGCCATCGCCCTGATAGACGGTTTTTCGCCCTTTGACGTTGGAGTCCACGTTCTTTAATAGTGGACTCTT
GTTCCAAACTGGAACAACACTCAACCCTATCTCGGGCTATTCTTTTGATTTATAAGGGATTTTGCCGATTTCGGCCTATTGG
TTAAAAAATGAGCTGATTTAACAAAAATTTAACGCGAATTTTAACAAAATATTAACGTTTACAATTTAAAAGGATCTAGGTG
AAGATCCTTTTTGATAATCTCATGACCAAAATCCCTTAACGTGAGTTTTCGTTCCACTGAGCGTCAGACCCCGTAGAAAAGA
TCAAAGGATCTTCTTGAGATCCTTTTTTTCTGCGCGTAATCTGCTGCTTGCAAACAAAAAAACCACCGCTACCAGCGGTGGT
TTGTTTGCCGGATCAAGAGCTACCAACTCTTTTTCCGAAGGTAACTGGCTTCAGCAGAGCGCAGATACCAAATACTGTCCTT
CTAGTGTAGCCGTAGTTAGGCCACCACTTCAAGAACTCTGTAGCACCGCCTACATACCTCGCTCTGCTAATCCTGTTACCAG
TGGCTGCTGCCAGTGGCGATAAGTCGTGTCTTACCGGGTTGGACTCAAGACGATAGTTACCGGATAAGGCGCAGCGGTCGGG
CTGAACGGGGGGTTCGTGCACACAGCCCAGCTTGGAGCGAACGACCTACACCGAACTGAGATACCTACAGCGTGAGCTATGA
GAAAGCGCCACGCTTCCCGAAGGGAGAAAGGCGGACAGGTATCCGGTAAGCGGCAGGGTCGGAACAGGAGAGCGCACGAGGG

AGCTTCCAGGGGGAAACGCCTGGTATCTTTATAGTCCTGTCGGGTTTCGCCACCTCTGACTTGAGCGTCGATTTTTGTGATG
CTCGTCAGGGGGGCGGAGCCTATGGAAAAACGCCAGCAACGCGGCCTTTTTACGGTTCCTGGCCTTTTGCTGGCCTTTTGCT
CACATGTTCTTTCCTGCGTTATCCCCTGATTCTGTGGATAACCGTATTACCGCCTTTGAGTGAGCTGATACCGCTCGCCGCA
GCCGAACGACCGAGCGCAGCGAGTCAGTGAGCGAGGAAGCGGTAGAGCGCCTGATGCGGTATTTTCTCCTTACGCATCTGTG
CGGTATTTCACACCGCATAGGGTCATGGCTGCGCCCCGACACCCGCCAACACCCGCTGACGCGCCCTGACGGGCTTGTCTGC
TCCCGGCATCCGCTTACAGACAAGCTGTGACCGTGTCCGGGAGCTGCATGTGTCAGAGGTTTTCACCGTCATCACCGAAACG
CGCGAGGCAGAAGGAGATGGCGCCCAACAGTCCCCCGGCCACGGGGCCTGCCACCATACCCACGCCGAAACAAGCGCTCATG
AGCCCGAAGTGGCGAGCCCGATCTTCCCCATCGGTGATGTCGGCGATATAGGCGCCAGCAACCGCACCTGTGGCGCCGGTGA
TGCCGGCCACGATGCGTCCGGCGTAGAGGATCTGCTCATGTTTGACAGCTTATCATCGATTTATATTCCCCAGAACATCAGG
TTAATGGCGTTTTTGATGTCATTTTCGCGGTGGCTGAGATCAGCCACTTCTTCCCCGATAACGGAGACTGGCACACTGGCCA
TATCGGTGGTCATCATGCGCCAGCTTTCATCCCCGATATGCACCACCGGGTAAAGTTCACGGGAGACTTTATCTGACAGCAG
ACGTGCACTGGCCAGGGGGATCACCATCCGTCGCCCGGGCGTGTCAATAATATCACTCTGTACATCCACAAACAGACGATAA
CGGCTCTCTCTTTTATAGGTGTAAACCTTAAACTGCATAGCGCACCGCAAAGTTAAGAAACC

#### ***Adaptor X (End)*** ###

Strictly required at the 3’ end of the most 3’ translation unit for recombination into pZT7-GoldenGate.

**pX-TU(X’) |** SapI, GG X’ site (*BsmBI*, BsmBI), BtsI, KanR, **BbsI**, ccdB

GAATATTGGGTTTAGTCTTGTTTCATAATTGTTGCAATGAAACGCGGTGAAACATTGCCTGAAACGTTAACTGAAACGCATA
TTTGCGGATTAGTTCATGACTTTATCTCTAACAAATTGAAATTAAACATTTAATTTTATTAAGGCAATTGTGGCACACCCCT
TGCTTTGTCTTTATCAACGCAAATAACAAGTTGATAACAAGCTAGCAGGAGGAATTCCATATGGGCTCTTCAGGGTAATCGA
T*GAGACG*TGGGGCACTGCTGAACTAGTCTGATACAGTCGACCTGCAGGCATGCAAGCTTGGCTGTTTTGGCGGATGAGAGAA
GATTTTCAGCCTGATACAGATTAAATCAGAACGCAGAAGCGGTCTGATAAAACAGAATTTGCCTGGCGGCAGTAGCGCGGTG
GTCCCACCTGACCCCATGCCGAACTCAGAAGTGAAACGCCGTAGCGCCGATGGTAGTGTGGCCAGAGCCCATGCGAGAGTAG
GGAACTGCCAGGCATCAAATAAAACGAAAGGCTCAGTCGAAAGACTGGGCCTTTCGTTTTATCTGTTGTTTGTCGGTGAACG
CTCTCCTGAGTAGGACAAATCCGCCGGGAGCGGATTTGAACGTTGCGAAGCAACGGCCCGGAGGGTGGCGGGCAGGACGCCC
GCCATAAACTGCCAGGCATCAAATTAAGCAGAAGGCCATCCTGACGGATGGCCTTTTTGCGTTTCTACAAACTCTCTGATCC
TTCAACTCAGCAAAAGTTCGATTTATTCAACAAAGCCACGTTGTGTCTCAAAATCTCTGATGTTACATTGCACAAGATAAAA
ATATATCATCATGAACAATAAAACTGTCTGCTTACATAAACAGTAATACAAGGGGTGTTATGAGCCATATTCAACGGGAAAC
GTCTTGCTCTAGGCCGCGATTAAATTCCAACATGGATGCTGATTTATATGGGTATAAATGGGCTCGCGATAATGTCGGGCAA
TCAGGTGCGACAATCTATCGATTGTATGGGAAGCCCGATGCGCCAGAGTTGTTTCTGAAACATGGCAAAGGTAGCGTTGCCA
ATGATGTTACAGATGAGATGGTCAGACTAAACTGGCTGACGGAATTTATGCCTCTGCCGACCATCAAGCATTTTATCCGTAC
TCCTGATGATGCATGGTTACTCACCACGGCGATCCCAGGGAAAACAGCATTCCAGGTATTAGAAGAATATCCTGATTCAGGT
GAAAATATTGTTGATGCGCTGGCGGTGTTCCTGCGCCGGTTGCATTCGATTCCTGTTTGTAATTGTCCTTTTAACAGCGACC
GCGTATTTCGTCTGGCTCAGGCGCAATCACGAATGAATAACGGTTTGGTTGATGCGAGTGATTTTGATGACGAGCGTAATGG
CTGGCCTGTTGAACAAGTCTGGAAAGAAATGCATAAACTTTTGCCATTCTCACCGGATTCAGTCGTCACTCATGGTGATTTC
TCACTTGATAACCTTATTTTTGACGAGGGGAAATTAATAGGTTGTATTGATGTTGGACGAGTCGGAATCGCAGACCGATACC
AGGATCTTGCCATCCTATGGAACTGCCTCGGTGAGTTTTCTCCTTCATTACAGAAACGGCTTTTTCAAAAATATGGTATTGA
TAATCCTGATATGAATAAATTGCAGTTTCATTTGATGCTCGATGAGTTTTTCTAACTGTCAGACCAAGTTTACTCATATATA
CTTTAGATTGATTT**GAAGAC**TACGCGCCCTGTAGCGGCGCATTAAGCGCGGCGGGTGTGGTGGTTACGCGCAGCGTGACCGC
TACACTTGCCAGCGCCCTAGCGCCCGCTCCTTTCGCTTTCTTCCCTTCCTTTCTCGCCACGTTCGCCGGCTTTCCCCGTCAA
GCTCTAAATCGGGGGCTCCCTTTAGGGTTCCGATTTAGTGCTTTACGGCACCTCGACCCCAAAAAACTTGATTTGGGTGATG
GTTCACGTAGTGGGCCATCGCCCTGATAGACGGTTTTTCGCCCTTTGACGTTGGAGTCCACGTTCTTTAATAGTGGACTCTT
GTTCCAAACTGGAACAACACTCAACCCTATCTCGGGCTATTCTTTTGATTTATAAGGGATTTTGCCGATTTCGGCCTATTGG
TTAAAAAATGAGCTGATTTAACAAAAATTTAACGCGAATTTTAACAAAATATTAACGTTTACAATTTAAAAGGATCTAGGTG
AAGATCCTTTTTGATAATCTCATGACCAAAATCCCTTAACGTGAGTTTTCGTTCCACTGAGCGTCAGACCCCGTAGAAAAGA
TCAAAGGATCTTCTTGAGATCCTTTTTTTCTGCGCGTAATCTGCTGCTTGCAAACAAAAAAACCACCGCTACCAGCGGTGGT
TTGTTTGCCGGATCAAGAGCTACCAACTCTTTTTCCGAAGGTAACTGGCTTCAGCAGAGCGCAGATACCAAATACTGTCCTT
CTAGTGTAGCCGTAGTTAGGCCACCACTTCAAGAACTCTGTAGCACCGCCTACATACCTCGCTCTGCTAATCCTGTTACCAG
TGGCTGCTGCCAGTGGCGATAAGTCGTGTCTTACCGGGTTGGACTCAAGACGATAGTTACCGGATAAGGCGCAGCGGTCGGG
CTGAACGGGGGGTTCGTGCACACAGCCCAGCTTGGAGCGAACGACCTACACCGAACTGAGATACCTACAGCGTGAGCTATGA
GAAAGCGCCACGCTTCCCGAAGGGAGAAAGGCGGACAGGTATCCGGTAAGCGGCAGGGTCGGAACAGGAGAGCGCACGAGGG
AGCTTCCAGGGGGAAACGCCTGGTATCTTTATAGTCCTGTCGGGTTTCGCCACCTCTGACTTGAGCGTCGATTTTTGTGATG
CTCGTCAGGGGGGCGGAGCCTATGGAAAAACGCCAGCAACGCGGCCTTTTTACGGTTCCTGGCCTTTTGCTGGCCTTTTGCT
CACATGTTCTTTCCTGCGTTATCCCCTGATTCTGTGGATAACCGTATTACCGCCTTTGAGTGAGCTGATACCGCTCGCCGCA
GCCGAACGACCGAGCGCAGCGAGTCAGTGAGCGAGGAAGCGGTAGAGCGCCTGATGCGGTATTTTCTCCTTACGCATCTGTG
CGGTATTTCACACCGCATAGGGTCATGGCTGCGCCCCGACACCCGCCAACACCCGCTGACGCGCCCTGACGGGCTTGTCTGC
TCCCGGCATCCGCTTACAGACAAGCTGTGACCGTGTCCGGGAGCTGCATGTGTCAGAGGTTTTCACCGTCATCACCGAAACG
CGCGAGGCAGAAGGAGATGGCGCCCAACAGTCCCCCGGCCACGGGGCCTGCCACCATACCCACGCCGAAACAAGCGCTCATG
AGCCCGAAGTGGCGAGCCCGATCTTCCCCATCGGTGATGTCGGCGATATAGGCGCCAGCAACCGCACCTGTGGCGCCGGTGA
TGCCGGCCACGATGCGTCCGGCGTAGAGGATCTGCTCATGTTTGACAGCTTATCATCGATTTATATTCCCCAGAACATCAGG
TTAATGGCGTTTTTGATGTCATTTTCGCGGTGGCTGAGATCAGCCACTTCTTCCCCGATAACGGAGACTGGCACACTGGCCA
TATCGGTGGTCATCATGCGCCAGCTTTCATCCCCGATATGCACCACCGGGTAAAGTTCACGGAGACTTTATCTGACAGCAG
ACGTGCACTGGCCAGGGGGATCACCATCCGTCGCCCGGGCGTGTCAATAATATCACTCTGTACATCCACAAACAGACGATAA
CGGCTCTCTCTTTTATAGGTGTAAACCTTAAACTGCATAGCGCACCGCAAAGTTAAGAAACC
